# Character evolution of modern fly-speck fungi and implications for interpreting thyriothecial fossils

**DOI:** 10.1101/2020.03.13.989582

**Authors:** Ludovic Le Renard, André L. Firmino, Olinto L. Pereira, Ruth A. Stockey, Mary. L. Berbee

## Abstract

**PREMISE OF THE STUDY:** Fossils show that fly-speck fungi have been reproducing with small, black thyriothecia on leaf surfaces for ∼250 million years. We analyze morphological characters of extant thyriothecial fungi to develop a phylogenetic framework for interpreting fossil taxa.

**METHODS:** We placed 59 extant fly-speck fungi in a phylogeny of 320 Ascomycota using nuclear ribosomal large and small subunit sequences, including newly determined sequences from nine taxa. We reconstructed ancestral character states using BayesTraits and maximum likelihood after coding 11 morphological characters based on original observations and literature. We analyzed the relationships of three previously published Mesozoic fossils using parsimony and our morphological character matrix, constrained by the molecular phylogeny.

**KEY RESULTS:** Thyriothecia evolved convergently in multiple lineages of superficial, leaf-inhabiting ascomycetes. The radiate and ostiolate scutellum organization is restricted to Dothideomycetes. Scutellum initiation by intercalary septation of a single hypha characterizes Asterinales and Asterotexiales, and initiation by coordinated growth of two or more adjacent hyphae characterizes Aulographaceae (order *incertae sedis*). Scutella in Microthyriales are initiated apically on a lateral hyphal branch. Patterns of hyphal branching in scutella contribute to distinguishing among orders. Parsimony resolves three fossil taxa as Dothideomycetes; one is further resolved as a member of a Microthyriales-Zeloasperisporiales clade within Dothideomycetes.

**CONCLUSIONS:** This is the most comprehensive systematic study of thyriothecial fungi and their relatives to date. Parsimony analysis of the matrix of character states of modern taxa provides an objective basis for interpreting fossils, leading to insights into morphological evolution and geological ages of Dothideomycetes clades.

---

Fly-speck fungi are ascomycetes that appear as minute (50 μm to 2 mm in diameter) black reproductive structures called thyriothecia. Thyriothecia are distinguished by having scutella, which are small, thin, darkly pigmented, shield-like plates that cover spore-producing cells. The margins of scutella are appressed to the surfaces of living or dead leaves or stems of vascular plants (Arnaud, 1917, 1918, 1925; Stevens and Ryan, 1939; Wu et al., 2011b; Wu et al., 2014). Fly-speck fungi are common throughout the world (Arnaud, 1918; Doidge, 1942; Batista, 1959; Batista et al., 1969; Holm and Holm, 1991), especially in tropical rainforests, where they grow on leaf surfaces (Batista, 1959; Hughes, 1976; Hofmann and Piepenbring, 2011; Firmino, 2016). Scutella of fly-speck fungi are also common as fossils (Elsik, 1978; Kalgutkar and Jansonius, 2000). Our interest in these fungi is rooted in a desire to interpret the phylogenetic significance of the hundreds of fossil forms found in palynological preparations and on preserved cuticles of plant fossils (Kalgutkar and Jansonius, 2000; Taylor et al., 2015). Fossil fly-speck scutella have long been classified into form genera and form species that carry minimal phylogenetic information. Phylogenetic classification of the fossils can potentially be improved by analysis of the systematic distribution of scutellum characters among extant fly-speck fungal lineages.

Scutella of thyriothecia are recognizable in part because they develop above the plant cuticle, in contrast to many other kinds of minute ascomycete fruiting bodies that begin within host tissue and break through to the leaf surface only when spores are ready for dispersal. How fly-speck fungi interact with their host plants is unclear, but some appear to be asymptomatic biotrophic parasites (on living hosts) while others are found in leaf litter (Hofmann, 2009; Wu et al., 2011b). Hyphae of some fly-speck fungi enter the host directly through a stoma (Hofmann, 2009; Firmino, 2016). Many other leaf inhabiting fungi, including fly-speck fungi, enter the host using appressoria. Appressoria are specialized hyphal cells that accumulate turgor pressure and direct growth into the host through a penetration peg, a small melanized ring (Parbery and Emmett, 1977). Appressoria then link the superficial fungal mycelium and various hyphal structures inside host tissue such as haustoria, hypostromata made of densely packed or stromatic hyphae, (Hofmann, 2009; Firmino, 2016), or palmettes of flattened, intercellular, dichotomous hyphae (Ducomet, 1907; Arnaud, 1918).

Several aspects of fly-speck fungus biology make it challenging to relate their morphology to phylogeny. First, thyriothecia are small, and so it can be difficult to find enough individuals of the same species for morphological or molecular systematic analysis. Second, fly- speck fungi appear in mixtures with other epiphyllous fungi, which can complicate distinguishing among species (Dilcher, 1965; Phipps and Rember, 2004). Hofmann (2009) and Firmino (2016) noted mixed species of *Asterina* growing together, and Ellis (1976) described closely related, co-occurring species in the collection of *Microthyrium microscopicum* that is the type for its genus. For sequence analysis, species should ideally be cultured. While some fly- speck fungi grow in pure culture (Firmino, 2016), other species do not (Hofmann, 2009); overall, they are poorly represented in culture collections.

A long-held view that thyriothecial fungi were monophyletic (Arnaud, 1918; Doidge, 1919; Clements and Shear, 1931; Gäumann, 1952; Kirk et al., 2008) has been corrected by a series of molecular phylogenetic analyses. Wu et al. (2011b) found a broad sampling of thyriothecial fungi to be scattered across four orders of Dothideomycetes, although with low support from bootstrap values or posterior probabilities. Mapook et al. (2016) and Hernández- Restrepo et al. (2019) showed that species of thyriothecial and thyriothecium-like fungi in Muyocopronales form a strongly supported sister group to Dyfrolomycetales, still in Dothideomycetes but distant from any other thyriothecial fungi. Even more distant is a monophyletic group of two thyriothecial *Micropeltis* species nested in class Lecanoromycetes rather than Dothideomycetes (Hongsanan and Hyde, 2017).

In spite of this recent progress, the challenges involved in collecting sufficient cells from fly-speck fungi remain, and their sequences are still poorly represented in GenBank and in phylogenetic studies. About 150 sequences were available, representing 18 genera, as of January 2019, contrasting with the more than 50 genera documented in recent publications (Hofmann, 2009; Wu et al., 2011a; Wu et al., 2011b; Hongsanan et al., 2014; Wu et al., 2014; Mapook et al., 2016), and close to ∼150 generic names available in families of thyriothecial fungi (Wijayawardene et al., 2014).

To interpret the fossil record of fly-speck fungi, our goals include increasing the sampling of extant representatives and related taxa in the class Dothideomycetes. Our project provides new cultures, herbarium specimens, and sequences into the public domain. Using new and previously published data, our first goal is to infer a molecular phylogeny that samples as widely as possible from the lineages of fly-speck fungi and their possible close relatives. Secondly, we score morphological characters for selected species in each lineage, and reconstruct ancestral morphological character states throughout the phylogeny. Thirdly, we use the molecular tree, comparative morphology, and inferred ancestral character states to discuss the phylogenetic affinities of three fly-speck fossils.

## MATERIALS AND METHODS

### Taxon sampling

We assembled a dataset of 320 ascomycete taxa including species of 59 fly-speck fungi. We obtained sequence data from new collections, from pure cultures where possible, found in Canada, Costa Rica (Permit n°R002-2014-OT-CONAGEBIO) and France.

Pure cultures were established by crushing individual thyriothecia under an inverted compound microscope using insect pins (Austerlitz Insect pins, size 0; Slavkov u Brna, Czech Republic) and then transferring individual spores or groups of spores onto water agar with 1 mM ampicillin and 1 mM kanamycin for germination. Germinating spores were transferred to antibiotic-free nutritive media under a dissecting microscope. The resulting cultures have been deposited in the Westerdijk Institute, Netherlands (Appendix S1, strains labeled with an asterisk).

Sequence data for nuclear ribosomal large subunit (LSU) and small subunit (SSU) DNA from thyriothecial taxa and their closest relatives were recovered from GenBank using a series of BLAST searches (Appendix S1). The use of these two loci was pragmatic; for most of the relatives of thyriothecial fungi including other leaf-inhabiting fungi and lichenized fungi, only ribosomal DNA data were available. We also included published sequences from lichenized fungi or lichen associated fungi that shared the epiphyllous habitat with thyriothecial fungi. We selected outgroups to represent classes related to Dothideomycetes. In addition, near full-length to full-length LSU and SSU data were retrieved from 13 whole genome projects with the help of BLAST searches and using the JGI MycoCosm Portal (Grigoriev et al., 2013) (Appendix S2).

### DNA extraction, amplification, sequencing

Strains successfully isolated in pure culture include *Lembosina* sp. CBS 144007, *Lembosina aulographoides* CBS 143809, *Microthyrium macrosporum* CBS 143810, and cf. *Stomiopeltis* sp. CBS 143811. They were harvested from culture media for genomic DNA extraction using the Qiagen DNeasy Plant Minikit (QIAGEN Inc., Mississauga, Ontario). A pure culture corresponding to the voucher of cf. *Stomiopeltis* sp. UBC-F33041 was isolated and its DNA extracted, but it could not be maintained over repeated transfers to new cultures. The polymerase chain reaction (PCR) reactions were performed using PureTaq Ready-To-Go PCR beads (Amersham Biosciences, Piscataway, New Jersey), following manufacturer instructions. Amplifications were performed using fungal specific rDNA primers reported in James et al. (2006): BMB-BR, NS1, NS3, NS6, NS8, NS20, NS23, NS51, ITS2, LIC1460R, LIC2197, LR0R, LR3R, LR5, LR5R, LR7R, LR8, LR10, LR11, LR10R, LR12, LR13. Uncultured dried specimens *Micropeltis* sp. UBC-F33034, *Scolecopeltidium* spp. UBC-F33033 and UBC-F33035, and Asterotexiaceae sp. UBC-F33036 were used as templates for direct DNA amplification (Lee and Taylor, 1990) of ∼1 kb of the 5’ end of the LSU, a region known to be informative for thyriothecial fungi (Hofmann et al., 2010; Wu et al., 2011b; Guatimosim et al., 2015). A cocktail of 12.5 µL of water and primers at a concentration of 1.0 µM each was pipetted into a PCR reaction tube containing a PureTaq Ready-To-Go PCR bead, and kept on ice. Thyriothecia were: (i) mounted in a drop of sterile water, observed under the compound microscope to ascertain identity, presence of ascospores, and absence of contaminant material; (ii) crushed firmly between a slide and a coverslip, and (iii) ∼10-13 µL of the crushed ascospore material in water was pipetted from the coverslip and the slide into a waiting PCR reaction tube, which was then vortexed briefly. The final volume of liquid in each tube was ∼25 µL, and the final concentration of each primer was 0.5 µM. The PCR reactions ran for 40 cycles with an annealing temperature of 50°C and an elongation temperature of 72°C. Direct amplifications of the 5’ end of the LSU using primers LR0R-LR8 with an annealing temperature of 55°C initially yielded weak PCR product. Weak PCR products were then diluted 100 to 1000 times and re-amplified with internal primers to increase product concentration for sequencing (see Mitchel, 2015, for similar methods).

### Phylogenetic inference

The LSU and SSU sequences were first aligned using the iterative refinement method G-INS-I in MAFFT v7 (Katoh and Standley, 2013). Introns were trimmed out manually and the resulting sequences re-aligned with MAFFT. Ambiguously aligned positions were excluded manually. The resulting concatenated dataset spanned 4552 bp. We ran JModelTest 2 (Darriba et al., 2012) on the CIPRES portal (Miller et al., 2010) and selected the GTR+G model for individual genes. We ran RAxML v.8 (Stamatakis, 2014) on two partitions (one for each gene) and 1000 independent maximum likelihood searches, again running on CIPRES. Branch support was assessed using 1000 bootstrap replicates (BS) (Felsenstein, 1985). Bayesian analysis was performed using MrBayes 3.2 Metropolis Coupled Monte Carlo algorithm (Ronquist et al., 2012). For the Bayesian analysis, we ran four independent searches, each with eight MCMC chains for 160 million generations sampling every 5000 trees, using 128 processors of 2 Gb each on a Compute Canada cluster (cedar.computecanada.ca). Following test runs, the heating parameter was adjusted to 0.03, optimizing the search space so that swap frequencies were between 0.3 and 0.7, as the MrBayes 3.2 manual advises. We discarded 50% of the samples as burn-in. We used Tracer 1.5 to determine that effective sample size exceeded 200 for all parameters, and the R package RWTY (Warren et al., 2017) to verify the convergence of independent runs. We considered clades with >70% likelihood bootstrap support and >0.95 Bayesian posterior probability to be moderately well supported.

Preliminary results suggested that some taxa at various ranks were not monophyletic. To explore the fit of the data to alternative hypotheses, we set monophyly constraints as single individual branches and found the most likely tree given each constraint using 100 independent “thorough” searches in RAxML. We then generated per-site log likelihoods of each likelihood tree given the constraint, and performed an approximately unbiased (AU) test (Shimodaira, 2002) using CONSEL (Shimodaira and Hasegawa, 2001). We rejected hypotheses receiving a probability below 0.05 in the AU test (Appendix S3). We specifically tested in turn the monophyly of (i) a putative clade comprising Asterinales and Asterotexiales, (ii) a putative clade comprising Microthyriales and Zeloasperisporiales, (iii) Radiate thyriothecia, (iv) *Asterina* and (v) *Lembosia*.

### Observation and scoring of morphological characters

Original observations of specimens were particularly important for coding thyriothecial characters that also appear in fossils. Table 1 lists the 26 specimens examined directly for morphological analysis. Scoring for the remaining 294 taxa was based on our interpretations of published illustrations and the literature (Appendix S4). When possible, we used morphological and sequence information from the same specimen (Appendix S4).

**Table 1.**
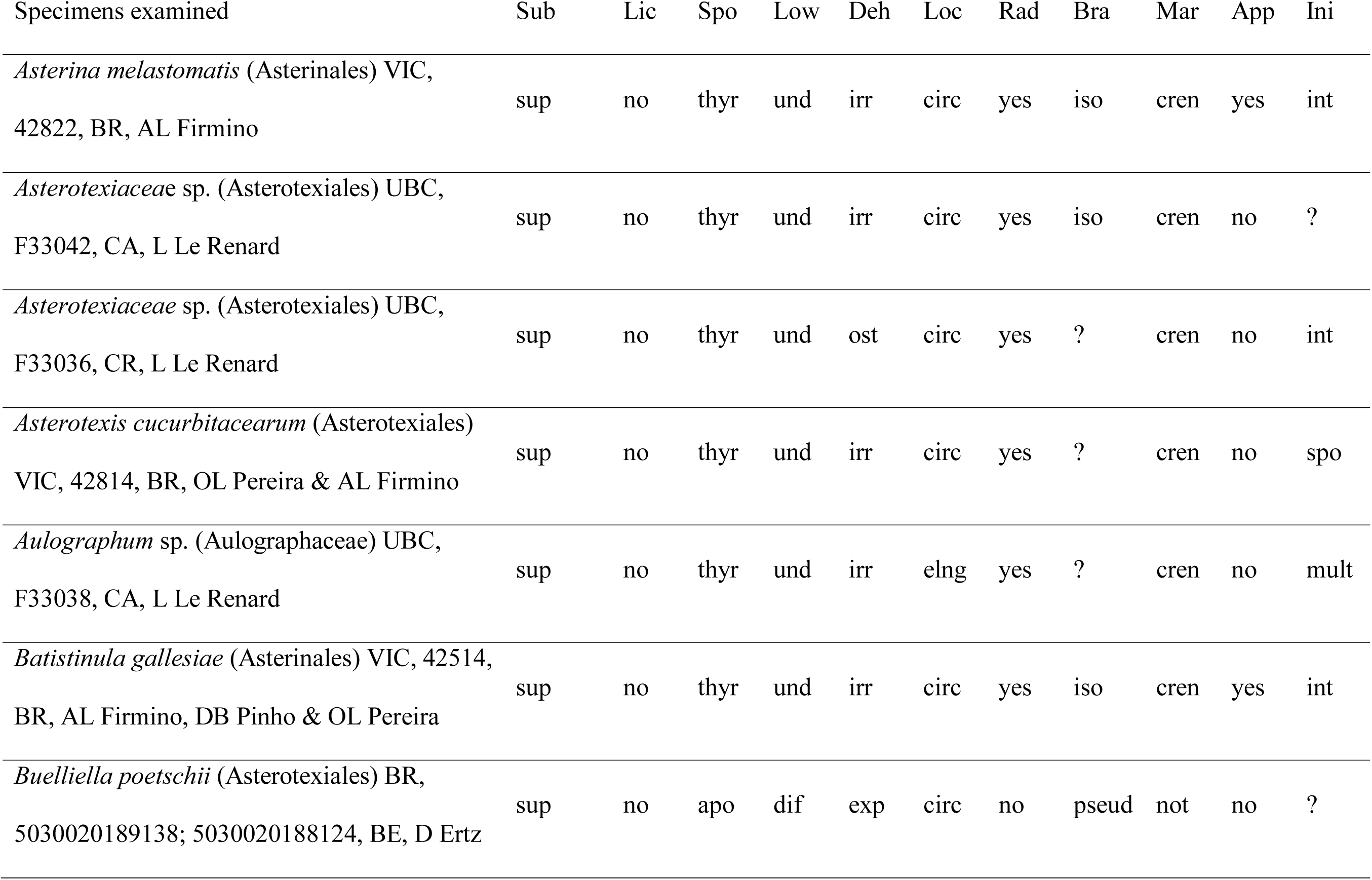
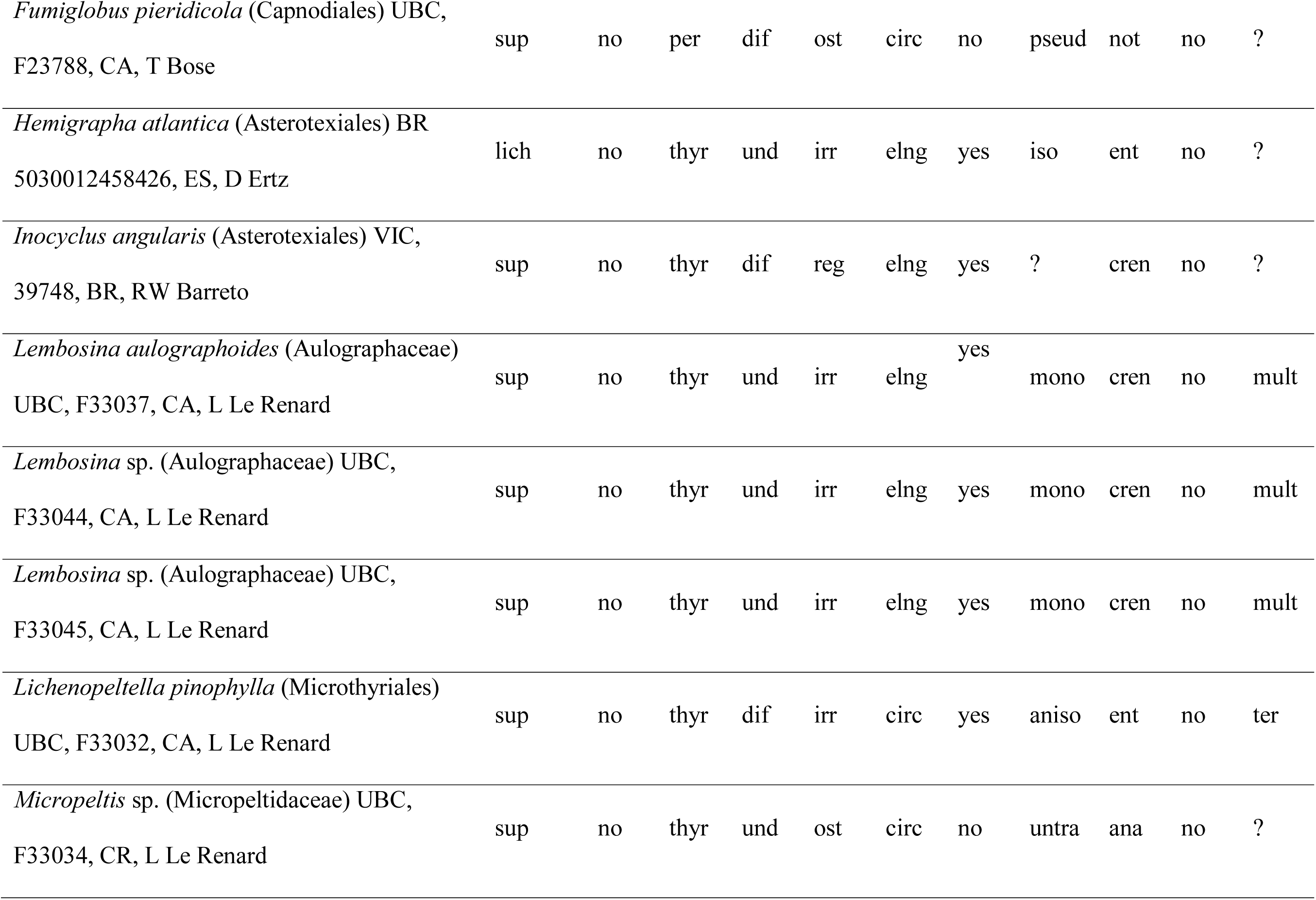
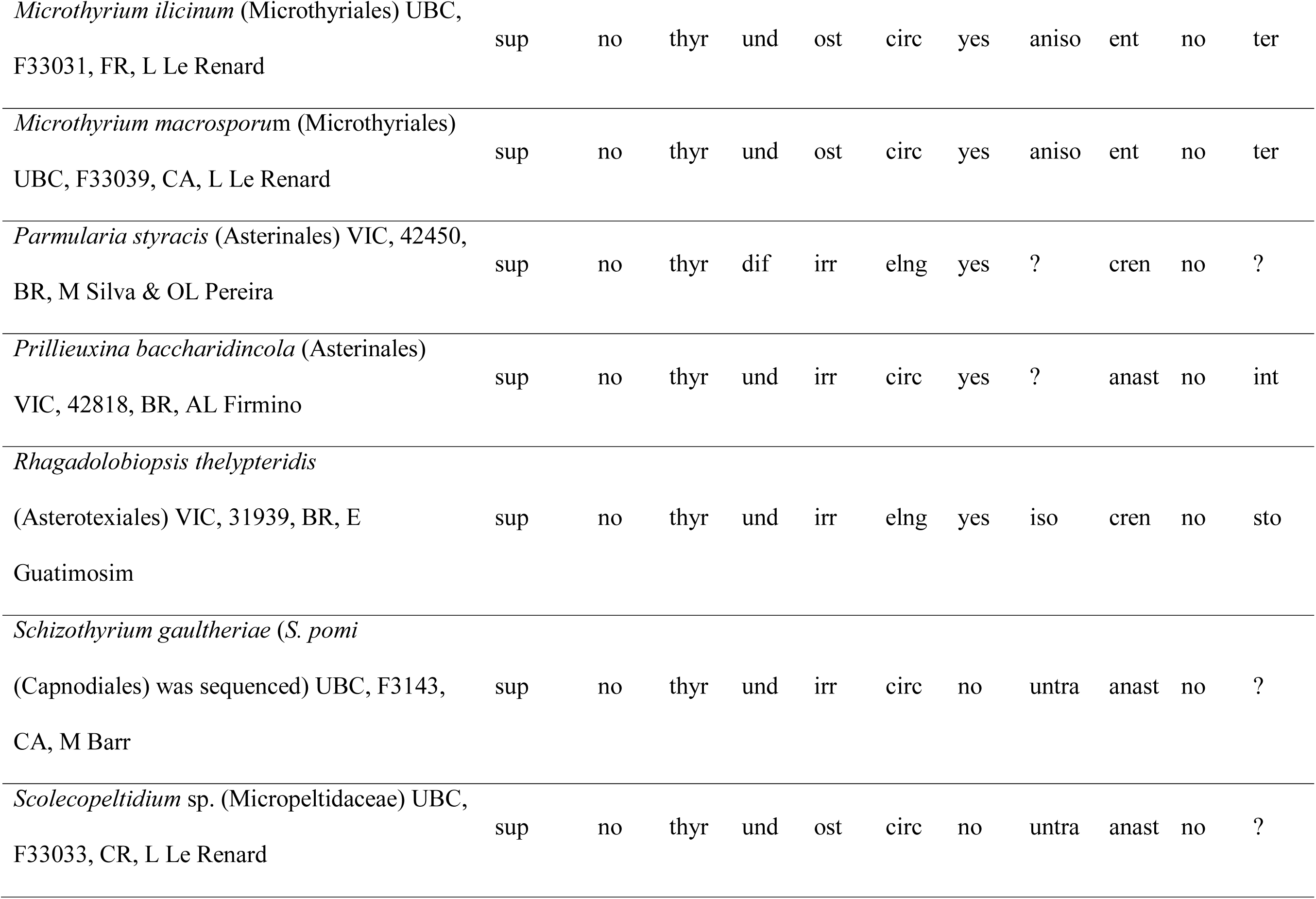
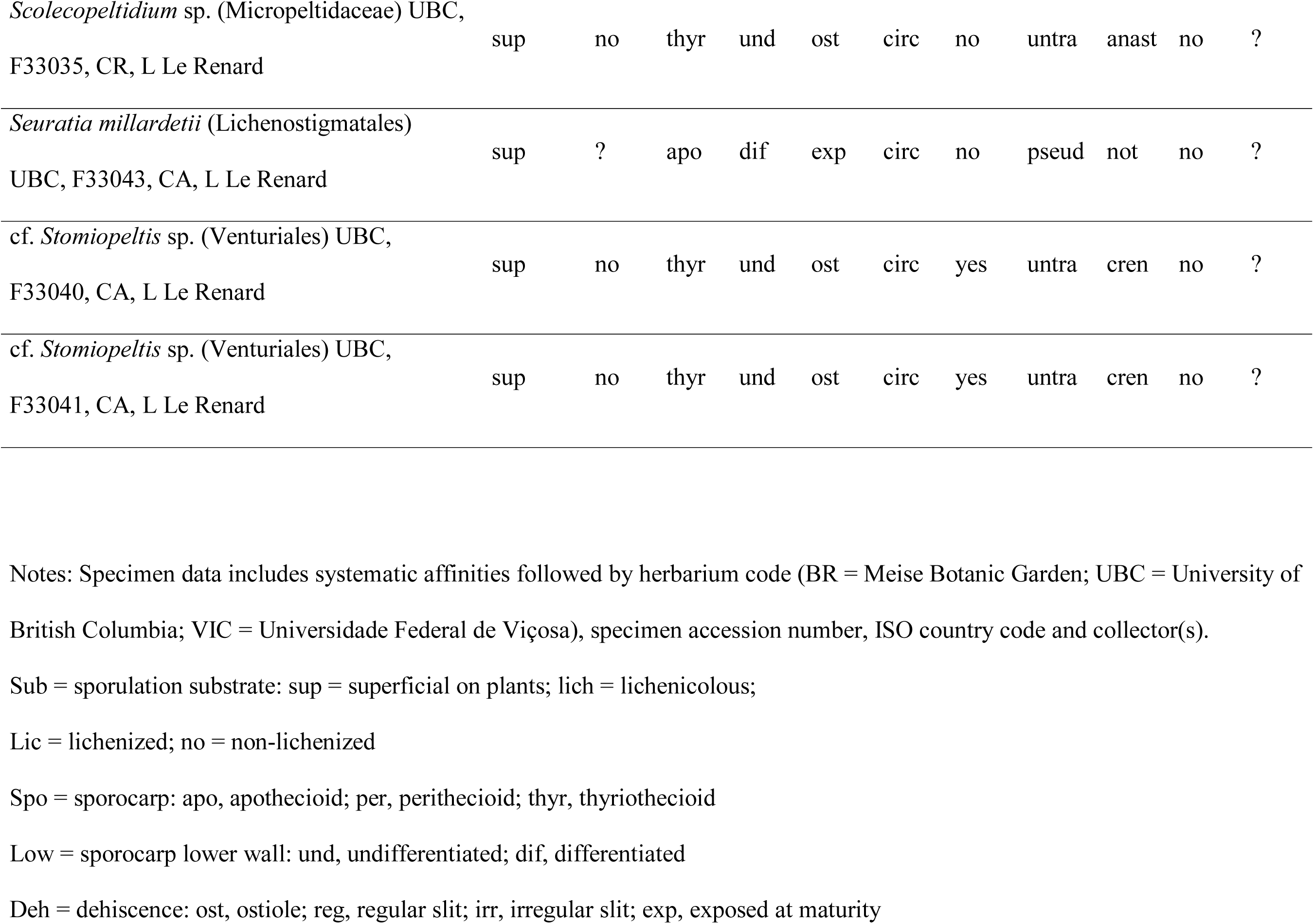
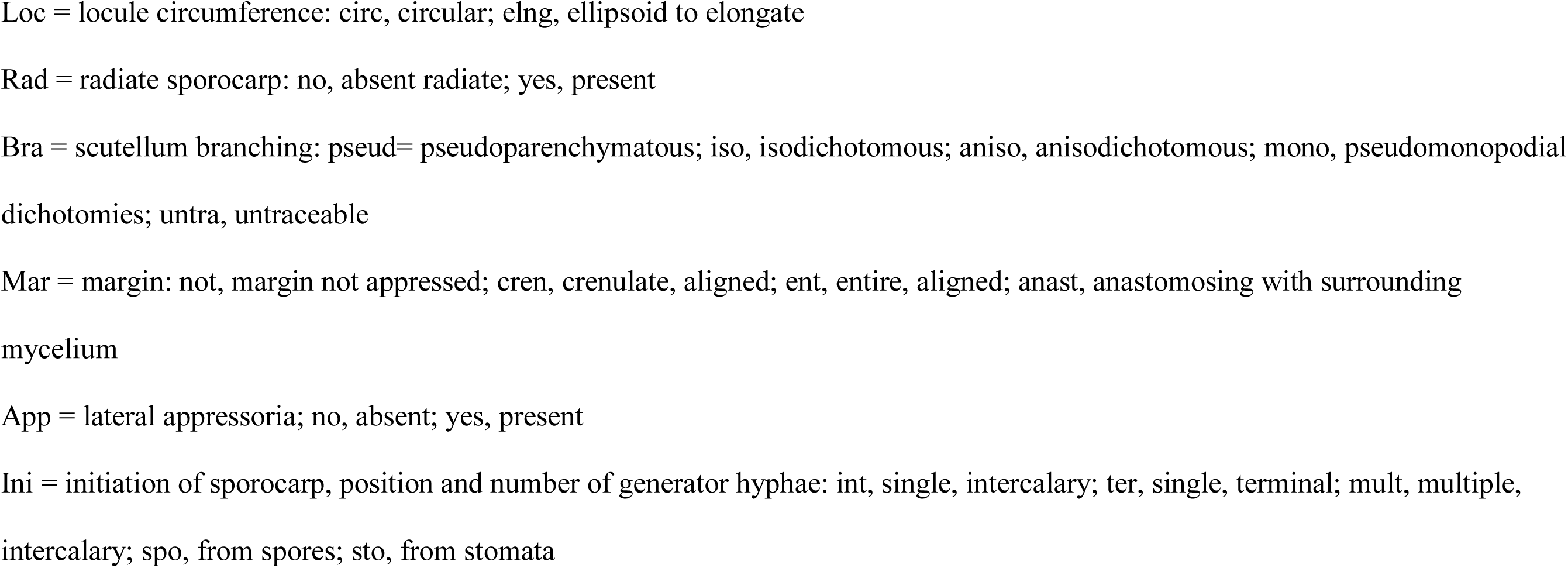
Matrix of specimens and their observed character states.

Table 2 shows our working definitions for scoring 11 morphological or habitat characters, considering sporocarp features of both asexual, conidium-producing structures and sexual ascomata. For direct observation of the hyphal structure of the dorsal surface of scutella, and to look for appressoria, we peeled thyriothecia together with their surface hyphae off of leaf surfaces using nail polish, as in Hosagoudar and Kapoor (1985). We examined thyriothecial features using a Leica DMRB (Leitz, Wetzlar, Germany) differential interference contrast (DIC) microscope, and photographed them at 400x or 1000x with a Leica DFC420 digital color camera. Many scutella were more cone-shaped than strictly flat, and investigating their hyphae required views from multiple focal planes. We attempted to use confocal microscopy to visualize hyphal organization, staining a specimen of *Microthyrium* with 0.1 mg/mL calcofluor white (Sigma, St. Louis, Missouri). The calcofluor revealed internal asci and some septa, but hyphae of the scutellum were obscured by autofluorescence and pigmentation, and we pursued this approach no further. Subsequently, we assembled stacks of bright field or DIC images at different focal planes to capture details of scutella and mycelia, following Harper (2015, p. 64, §4.3.1) and suggestions from Bercovici et al. (2009).

**Table 2.**
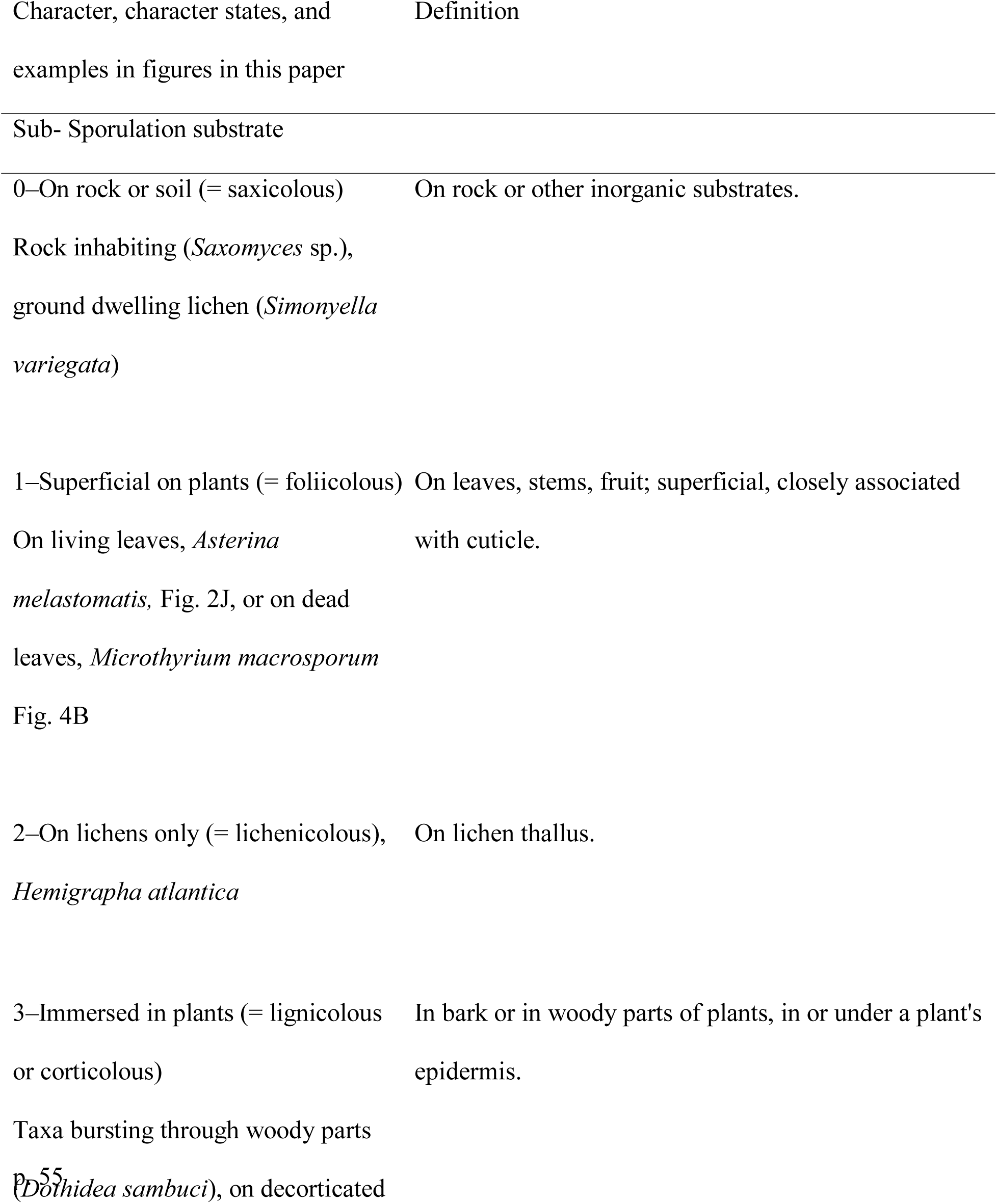
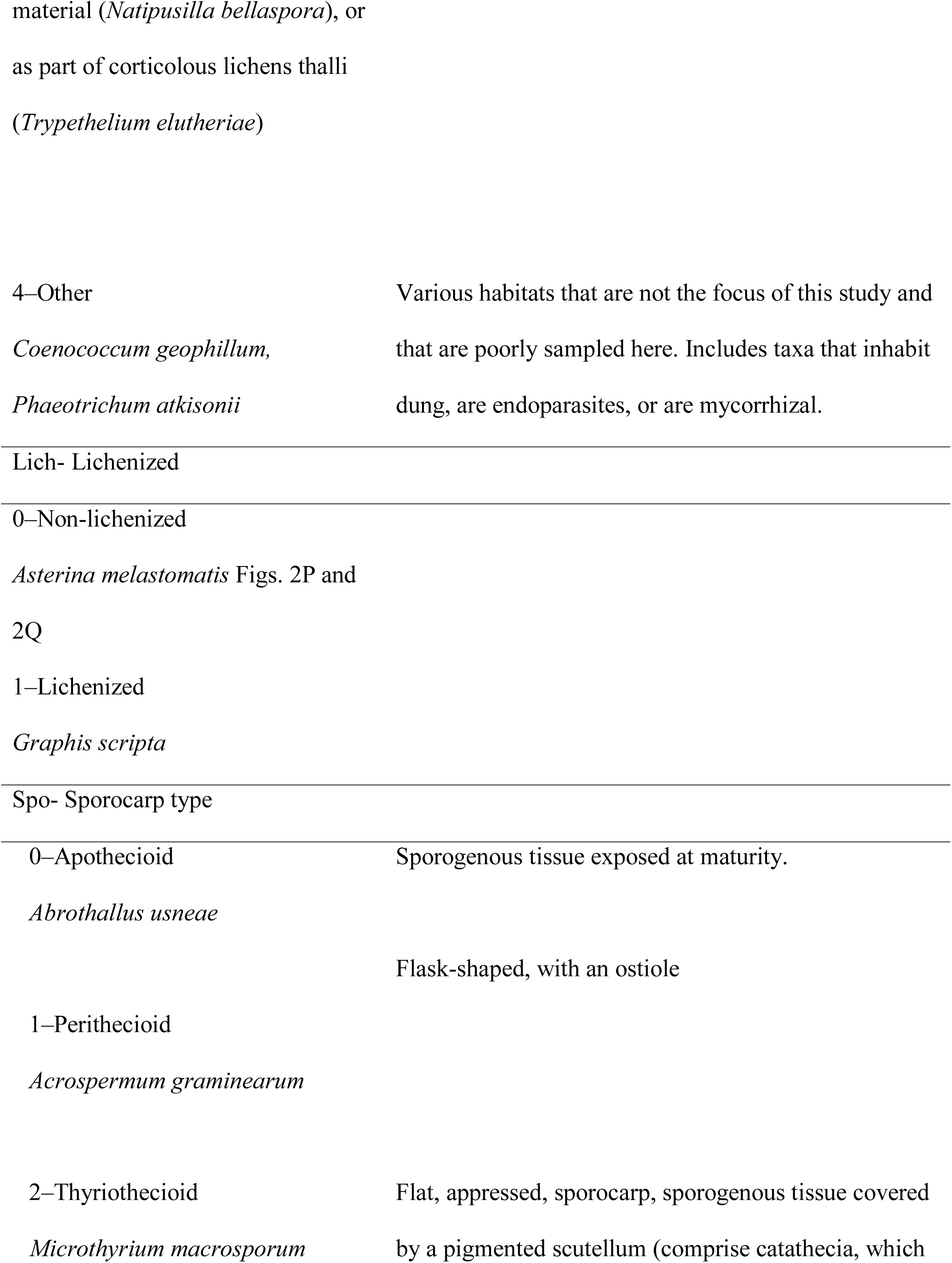
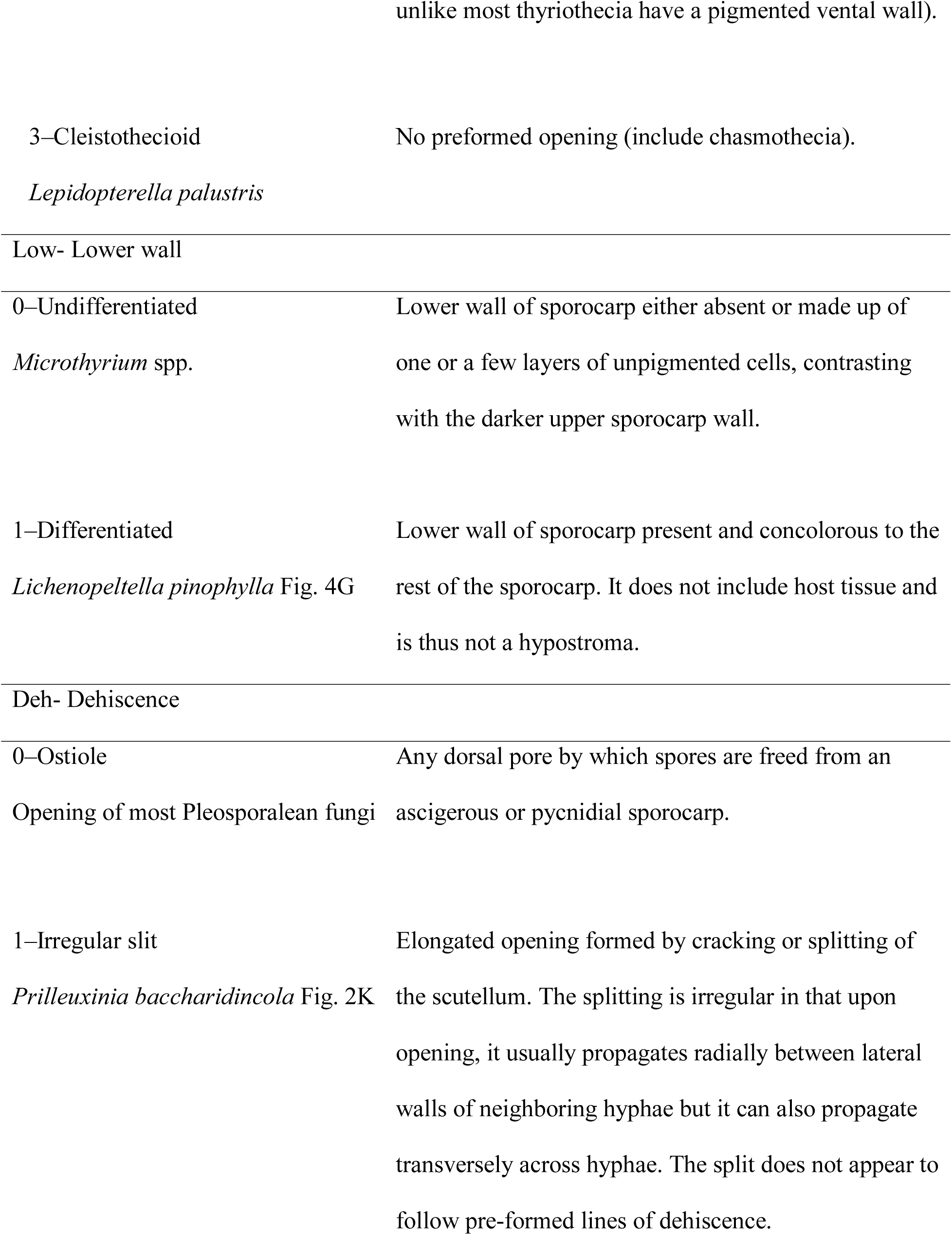
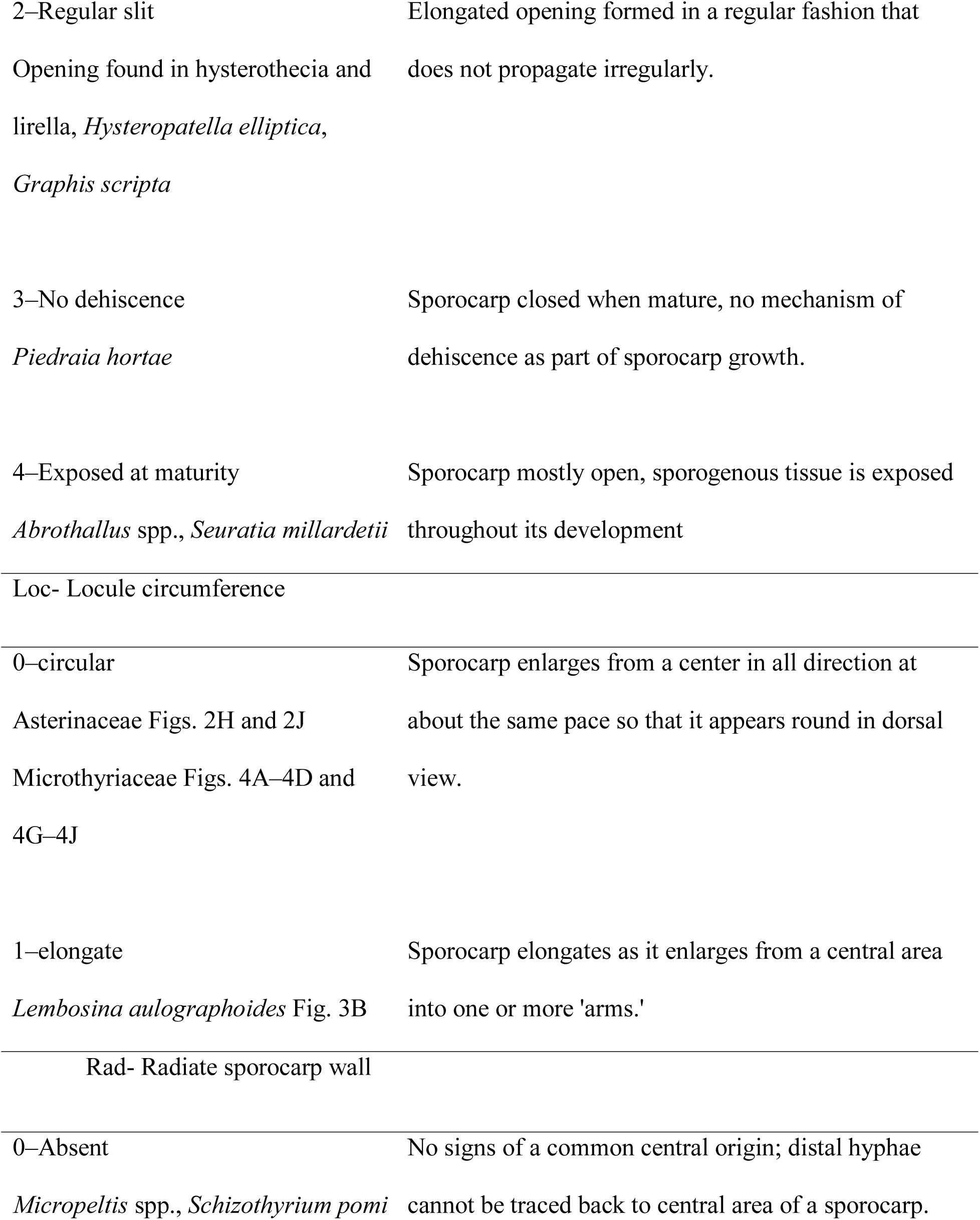
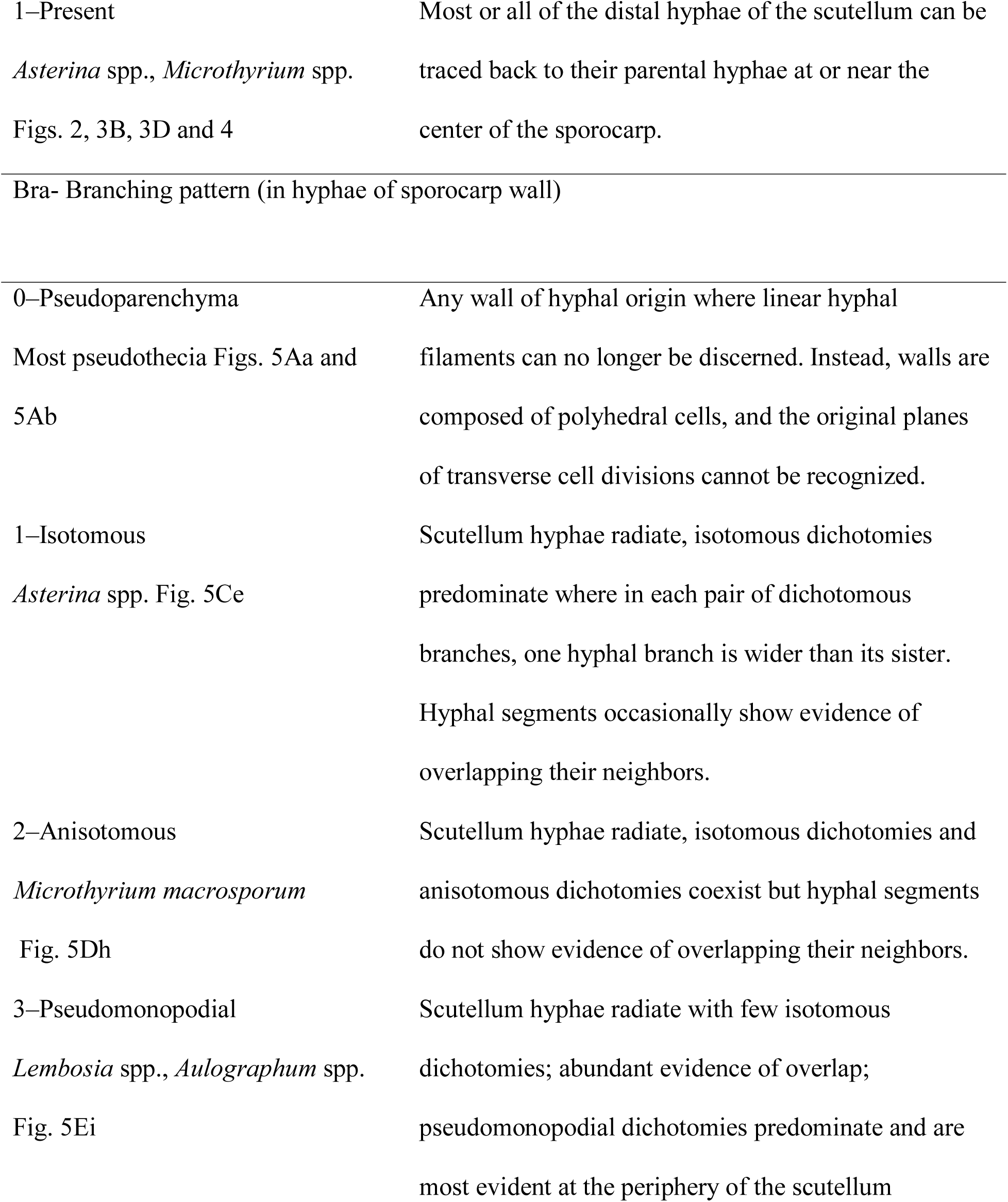
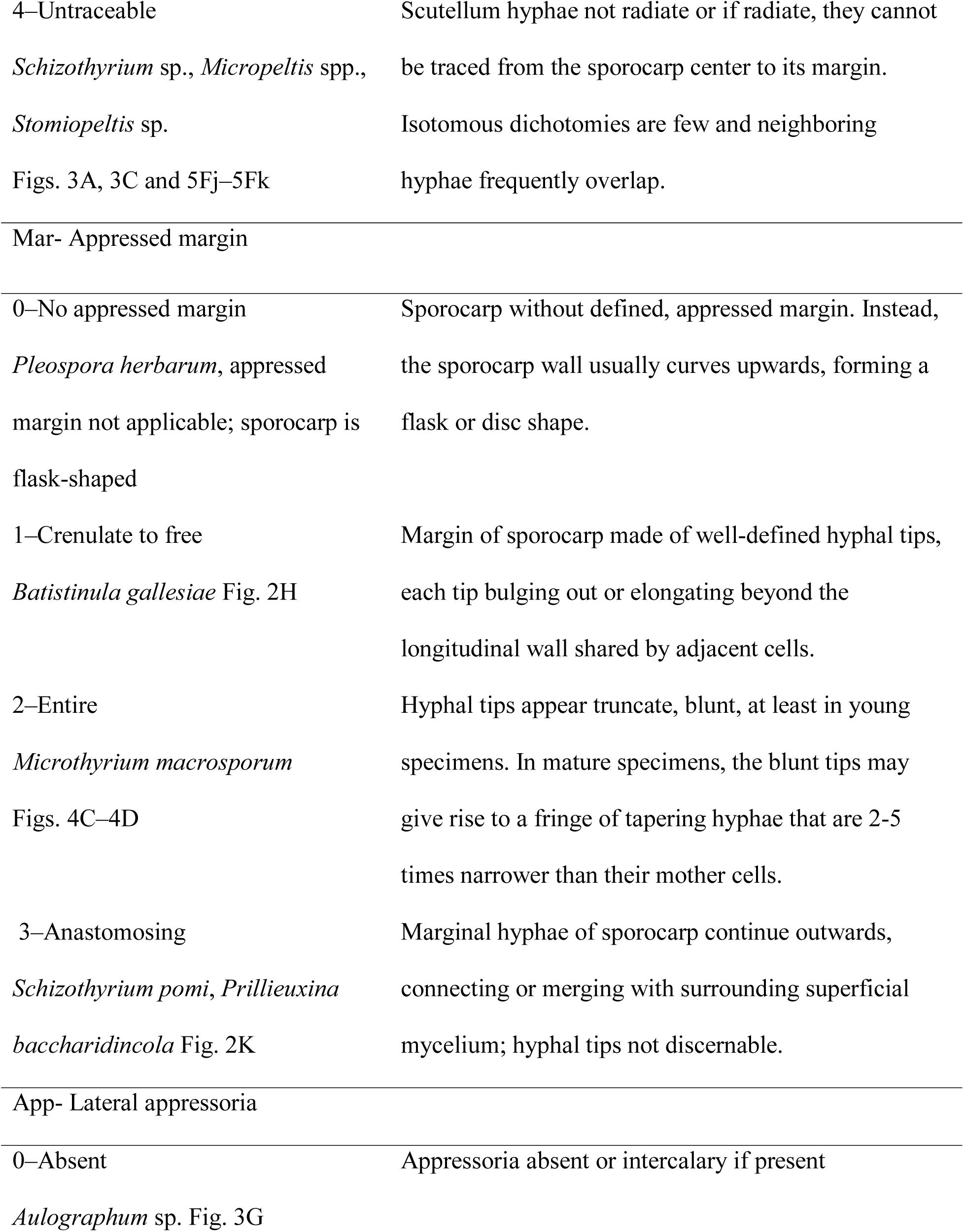
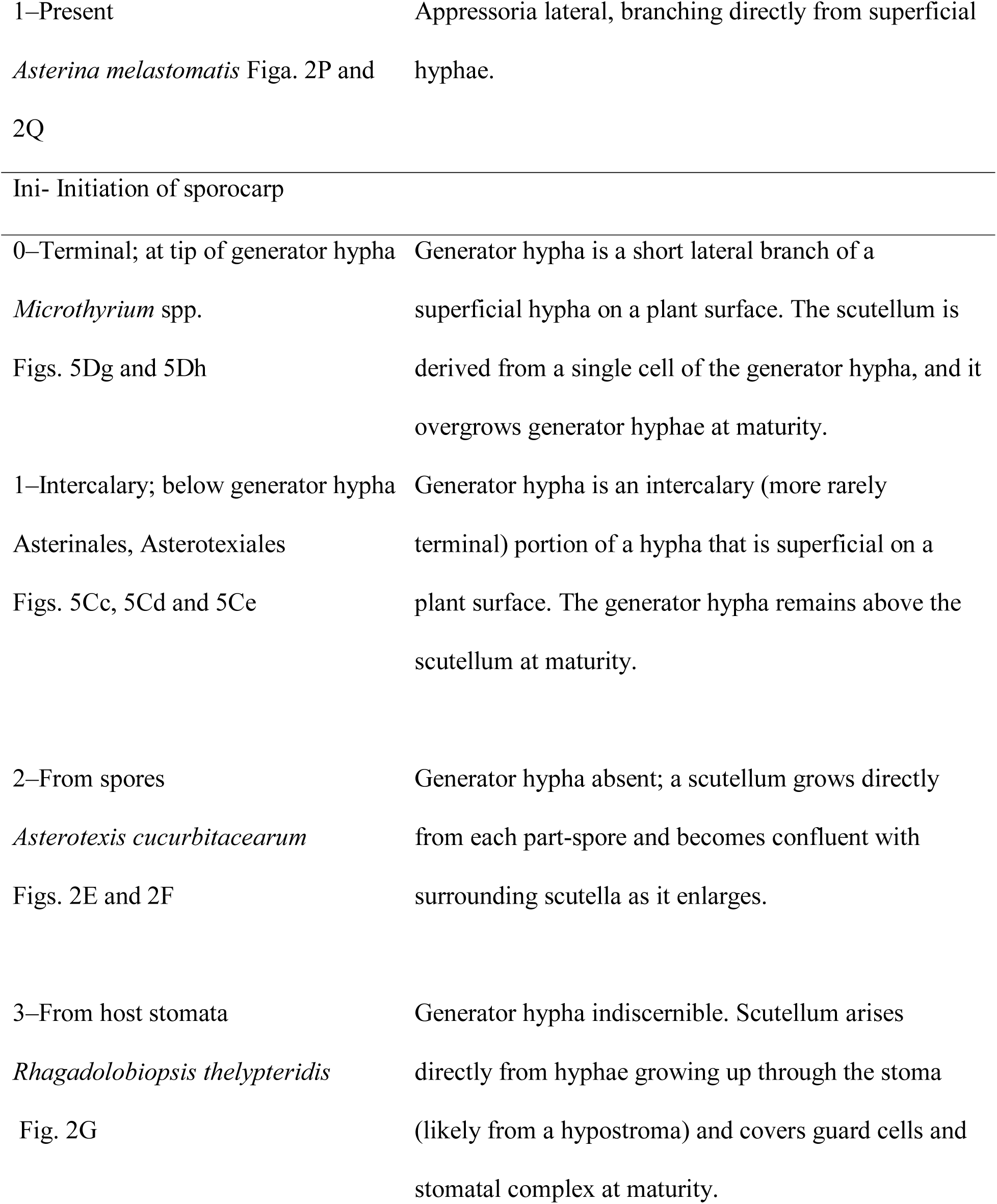
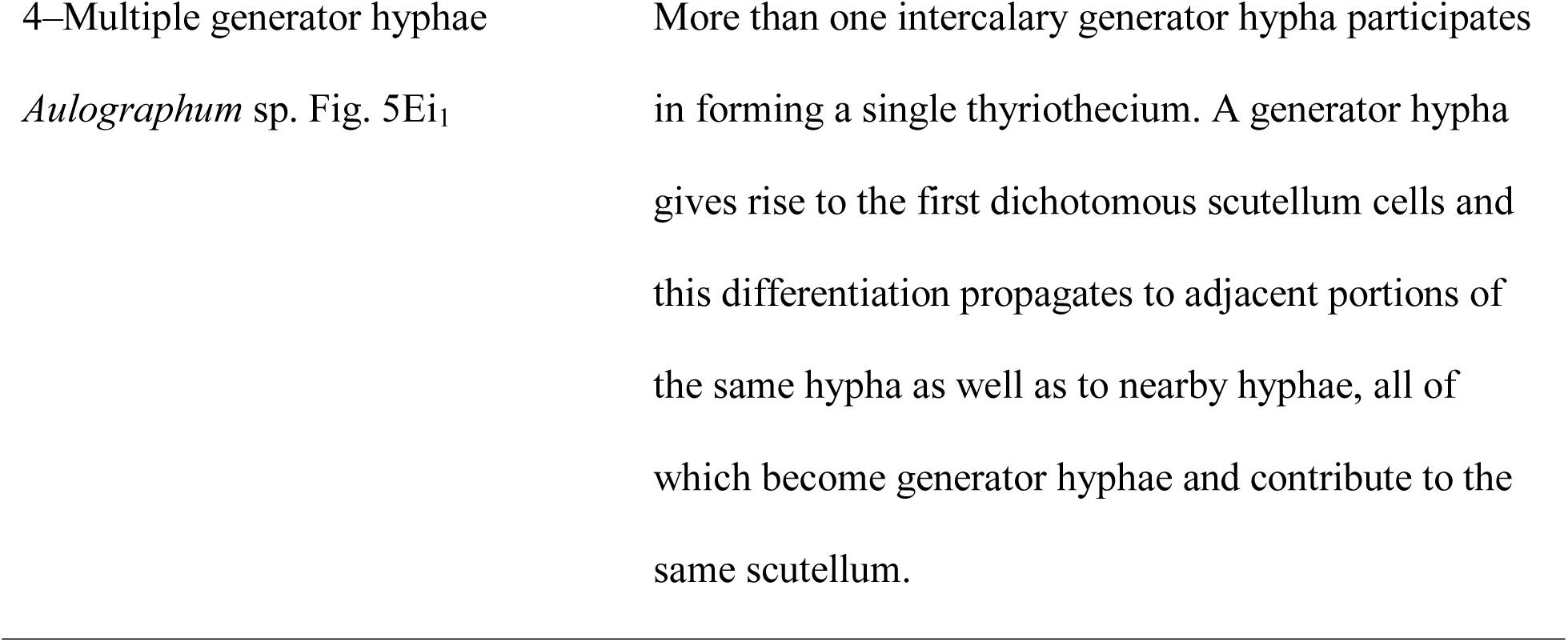
Definitions of characters and their states with reference to examples from illustrations in figures.

Published written descriptions provide some character state information but are rarely detailed enough to allow coding of the more subtle characters. Of the 11 scored characters, 10**–** 100% could be scored per specimen, across the 320 specimens studied (Appendix S4).

Appressoria are an example of a character that is rarely reported or illustrated, visible in surface mycelium of extant and fossil thyriothecial taxa, but rarely noted in publications on other fungi. Six characters specific to sporocarps could be scored for 67**–**79% of the taxa included (Appendix S4). Thyriothecial development could be coded in extant and fossil taxa because different stages are exposed on the surfaces of leaves. Development in non-thyriothecial Dothideomycetes is more difficult to observe, again leading to much missing data (Appendix S4).

### Analysis of morphological characters

We performed maximum likelihood ancestral character state reconstruction on the most likely tree in Mesquite for the scored characters.

Maximum likelihood reconstructions were computed using the Markov k-state 1-parameter (MK1) model (Lewis, 2001). Likelihood ratio tests showed that the MK1 model, which assumes equal rates of forward and reverse transitions between states for each character was more likely than an asymmetrical, 2-parameter model for the five binary characters in our dataset. We reconstructed character states on the single, most likely tree.

To take phylogenetic uncertainty into account, we also performed reconstruction using BayesTraits V3 (Pagel and Meade, 2007). In BayesTraits, we used the Multistate ML search algorithm, which estimates different rates for every state transition, with the option MLtries set to 100, a setting which led to consistent results in preliminary runs. Each character was reconstructed for 208 nodes out of the 319 nodes of our tree. The proportion of each state for each node was calculated from its mean value over the 5000 trees chosen randomly from the among the 32,000 trees in the posterior distribution of trees in chain one of our MrBayes analysis. We considered any character state reconstructions with > 0.7 of both posterior probability (from BayesTraits) and proportional likelihood (maximum likelihood) to be supported.

### Analysis of fossils

We selected three fossils reported as thyriothecial from the literature that were well enough illustrated to be coded for morphology. These include the oldest dispersed thyriothecium-like structure (Mishra et al., 2018), a dispersed *Lichenopeltella*-like form (Monga et al., 2015) for which cell patterns could be analyzed, and *Asterina eocenica* found on a leaf associated with mycelium (Dilcher, 1965). As for extant taxa, we coded character states from the illustrations (Appendix S4).

To infer the phylogenetic relationships of each of the three fossils, we used the most likely, 320-taxon tree from the rDNA analysis as a topological constraint. We analyzed the relationships of the fossils together and then one at a time, using the 11-character morphological data set (Appendix S4). With branch-and-bound searches in PAUP 4.0a166 (Swofford 2003), we found the most parsimonious positions of the fossils, given the rooted constraint tree. We did not further pursue the simultaneous analysis of all three fossils because the result was a poorly resolved, uninformative phylogeny (not shown). Instead, we focus on the results from adding one fossil at a time. Alignments, phylograms, and cladograms are available from TreeBASE (Submission ID: <u>25326</u>).

## RESULTS

### Distribution of thyriothecium forming fungi

Thyriothecium forming fungi appear polyphyletic, as expected (Fig. 1). All but Micropeltidaceae are included in class Dothideomycetes (Fig. 1). Within Dothideomycetes, thyriothecial fungi appear widely distributed, arising from a poorly resolved backbone (Fig. 1; Appendices S5 and S6).

**Figure 1.**
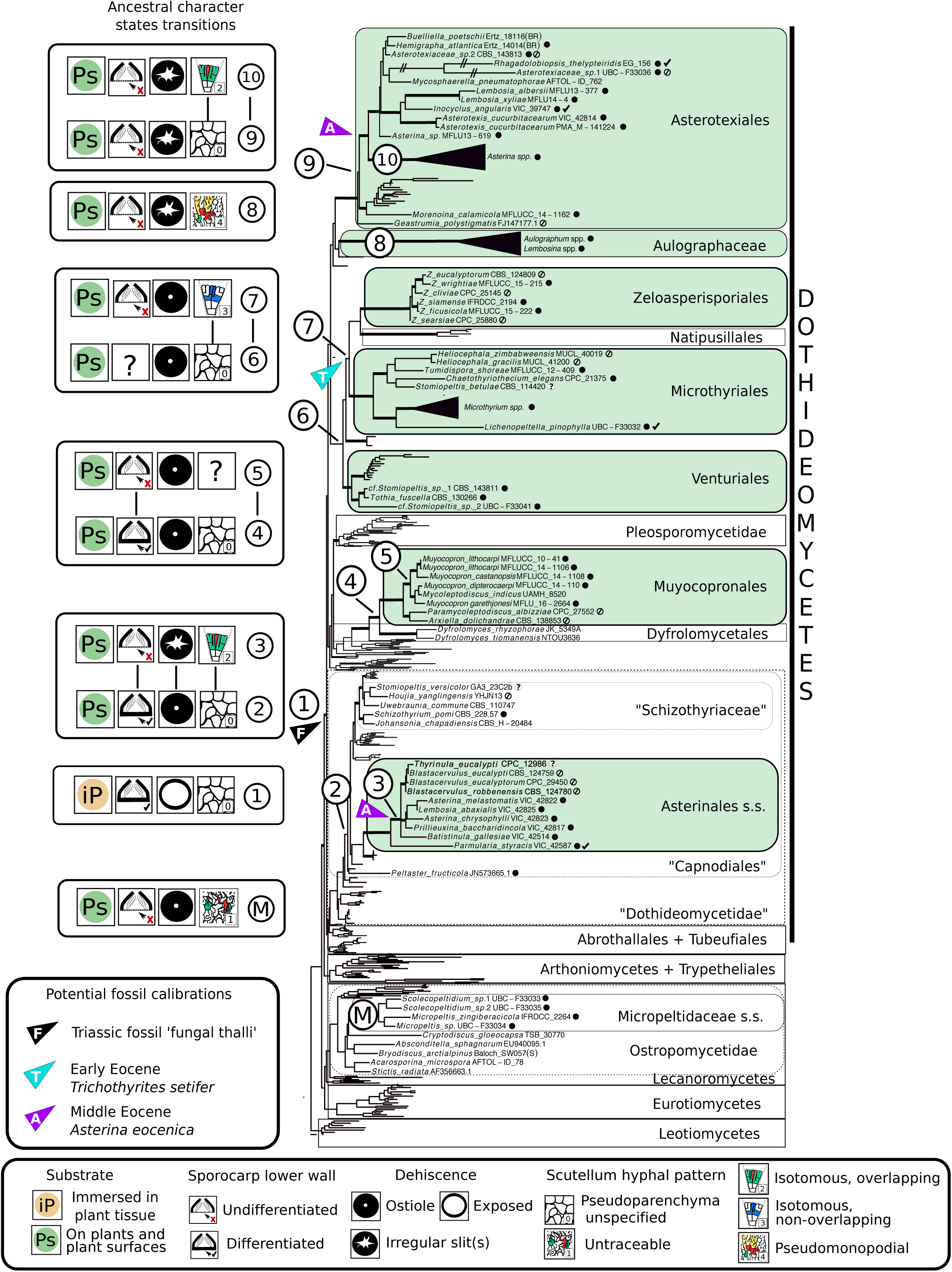
Phylogeny and character state transitions leading to convergent evolution of fly-speck fungal morphology. Numbers on boxes to the left match labeled transitions between reconstructed ancestral states that received > 0.7 proportional likelihood in the maximum likelihood phylogeny to the right. Thickened branches in the tree indicate bootstrap support > 70% and posterior probabilities > 0.95. Arrowheads indicate the most parsimonious consensus positions (for *A. eocenica*, alternative consensus positions) of fossils. Green boxes surround clades that include taxa with radiate thryiothecia. Solid black circles after taxon names indicate thyriothecial species; question marks indicate possible thryiothecia still in need of anatomical analysis; slashed circles indicate taxa known only from their asexual state; and tick marks indicate thyriothecia with differentiated lower walls.

Asterotexiales form a well-supported clade rich in thyriothecial forms, including the type species *Asterotexis cucurbitacearum* (Fig. 1; Appendices S5 and S6). Of the 35 Asterotexiales taxa sampled, 20 have thyriothecial sporocarps and 19 have epiphyllous habitats (Appendix S4). Clade A, one of three clades diverging near the base of the Asterotexiales, includes 19 taxa with thyriothecia, and 18 of the species that are epiphyllous (Appendices S4 and S6). However, Clade A also includes *Buelliella poetschii*, an apothecioid lichen parasite, and *Mycosphaerella pneumatophorae*, which produces a perithecioid ascoma in (rather than superficially on) plant tissue. Note that the “ioid” suffix refers to sporocarp shape, not to development of the ascomata (see character state definitions in Table 2).

Clade B species were recently classified in Asterinales (see Ertz and Diederich, 2015) but here, they group instead with Asterotexiales (Appendices S5 and S6). Clade B includes 14 species, most of which are apothecioid parasites of lichens, although two are corticolous, and one species, *Morenoina calamicola* is thyriothecial and epiphyllous (Appendices S4, S5 and S6). The third clade is represented by *Geastrumia polystigmatis*, an epiphyllous species known only by its asexual state, and it has yet to be formally classified to order. Aulographaceae, represented by six thyriothecial taxa, appears as the sister group of Asterotexiales, but with low branch support.

A clade of five orders including Microthyriales and Venturiales receives 88% bootstrap support and 100% posterior probability (Fig. 1; Appendices S5 and S6). Of the orders, Microthyriales and Zeloasperisporiales are thyriothecial and epiphyllous, but their close relatives in Natipusillales are freshwater cleistothecioid fungi that fruit on decorticated wood (Fig. 1; Appendices S4 and S6). Although Venturiales include a few thyriothecial species, most of its members are perithecioid leaf and stem parasites causing, for example, apple scab and black knot of plum (Fig. 1; Appendices S4 and S6).

Most Capnodiales are perithecioid, but the order also includes thyriothecial species (Fig. 1; Appendix S4). The order Asterinales is nested in Capnodiales with 99% Bayesian posterior probability but with negligible likelihood bootstrap support (Appendix S6). Asterinales is represented here by the type species *Asterina melastomatis*, and by sequences from three collections of *Parmularia styracis*, the type species of Parmulariaceae (Appendices S5 and S6). Asterinales also includes anamorphic foliicolous fungi and *Thyrinula eucalypti*, a plant pathogen that causes leaf spots around its thyriothecia. Two thyriothecial taxa in Capnodiales are not closely related to one another or to Asterinales: *Peltaster fructicola* and *Schizothyrium pomi*. The genus *Stomiopeltis* is polyphyletic and *Stomiopeltis betulae* is resolved in Microthyriales while two species tentatively identified as *Stomiopeltis* form a clade with *Tothia fuscella* in Venturiales (Fig. 1; Appendices S5 and S6).

Muyocopronales represents a strongly supported example of convergent origin of thyriothecia (Fig. 1; Appendices S5 and S6). Muyocopronales thyriothecia have several cell layers of lightly pigmented, pseudoparenchymatous cells below their outer, darkly pigmented scutellum (see illustrations of Mapook et al., 2016b). The thickness of the scutellum makes detailed analysis of the hyphal branching pattern challenging.

Micropeltidaceae provide another strongly supported example of convergent thyriothecial morphology. The family is nested in Ostropomycetidae in Lecanoromycetes rather than Dothideomycetes (Fig. 1; Appendices S5 and S6). The scutellum of Micropeltidaceae is distinctive in that it is ostiolate and formed as a flat, compact reticulum-like network of overlapping hyphae. As illustrated by Hofmann and Piepenbring (2006), the hyphae do not radiate from the center to the margins of the scutellum.

### Character states of most recent common ancestors of thyriothecial clades

*Bayesian vs likelihood results*—Different methods of character state reconstruction are generally congruent, but BayesTraits more frequently gives unresolved ancestral states compared with likelihood (Table 3; Appendices S6 and S7). BayesTraits reconstructs states over a posterior distribution of trees, so topological conflict could have explained low support for states. Surprising, in our analysis, unresolved ancestral states in BayesTraits are often associated with clades that have strong topological support, indicating that conflicting branching order is not the only cause of lack of resolution. Rather, the unresolved nodes are associated with missing morphological data and high estimated rates of character state transitions. BayesTraits estimates the forward and reverse rates separately for each character, and where data are missing, the information content is insufficient to parameterize the separate rates. An example of this is evident in the reconstruction of the scutellum branching pattern in the ancestor of Zeloasperisporiales and Natipusillales (Table 3). For this character, BayesTraits estimated an average forward transition rate from pseudoparenchymatous (0) to anisotomous branching (2) as 4.3 but it estimated the reverse rate as 56. Near the reconstruction, the topology is well supported and most of the surrounding branches receive ≥92% posterior probabilities but BayesTraits shows the reconstructed state as ≥ completely equivocal (0.2 for each of five states) (Appendices S6 and S7). In contrast, the same node is reconstructed as anisotomous with 0.85 proportional likelihood (Appendices S6 and S7) because the MK1 model can take advantage of all available data to estimate the single transition rate that it applies to each character. We consider results to be most reliable when supported by both likelihood and Bayesian reconstructions. However, when the topology is well supported, we consider a reconstruction supported by likelihood to be more useful than an equivocal Bayesian result.

**Table 3.**
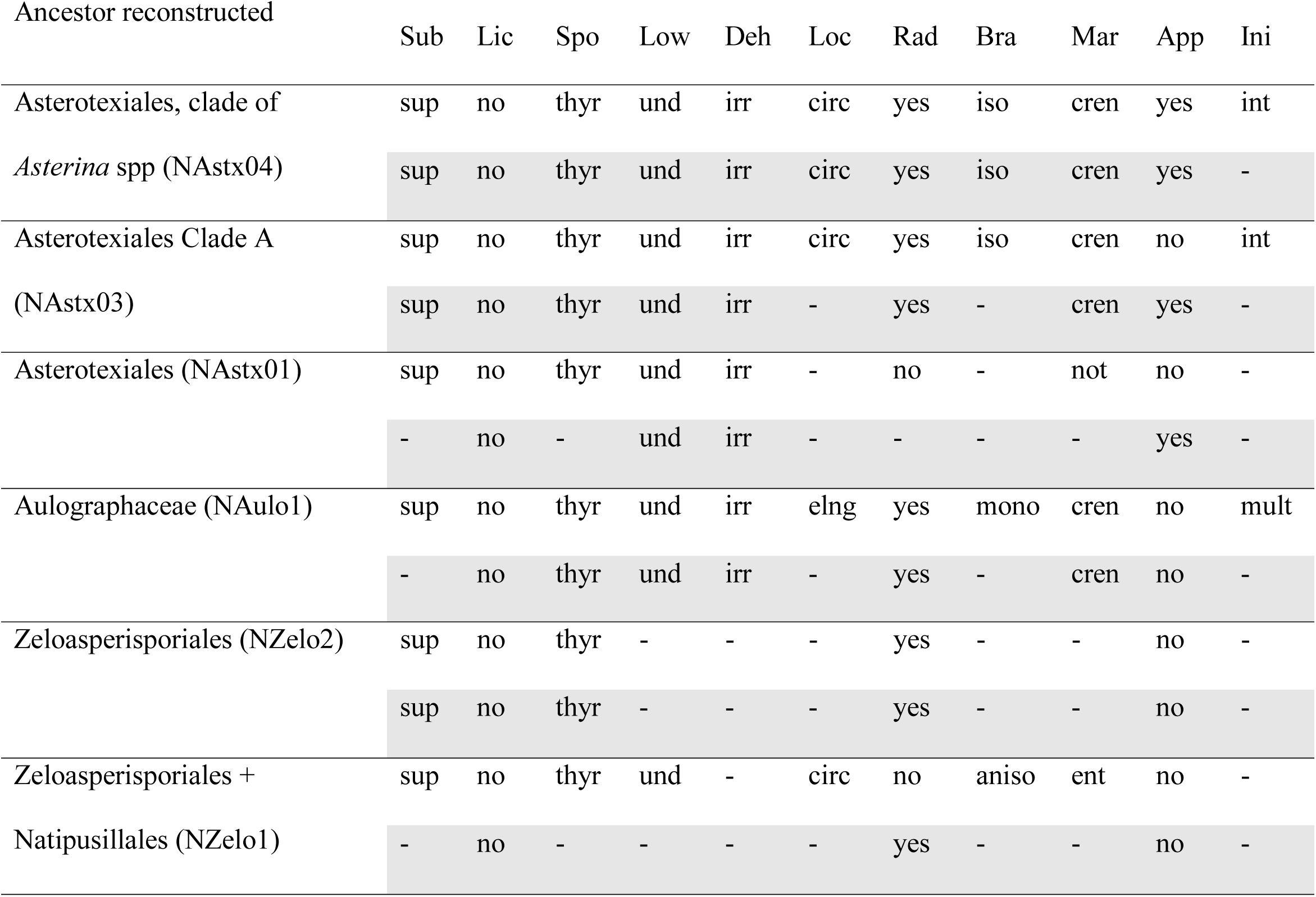
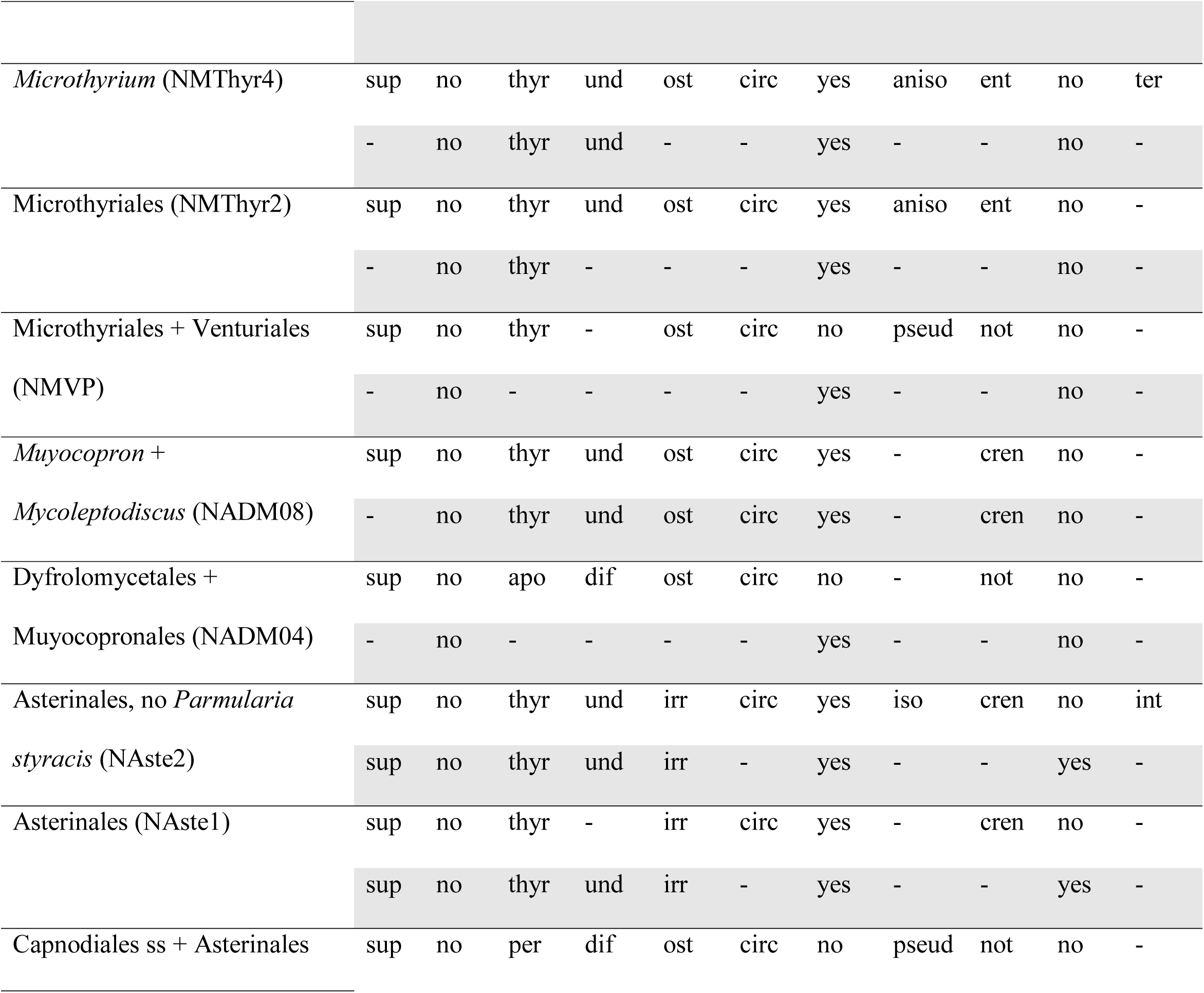
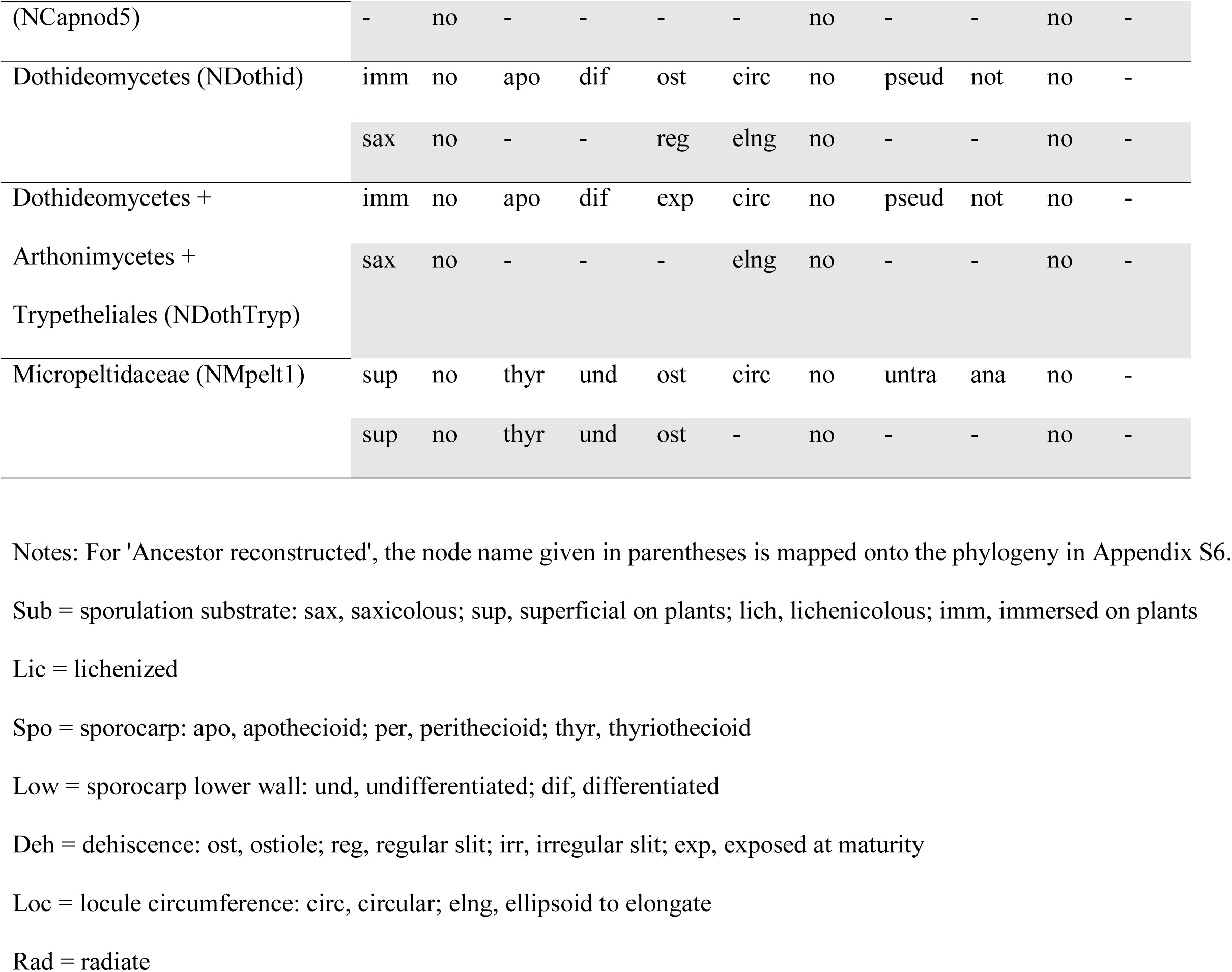
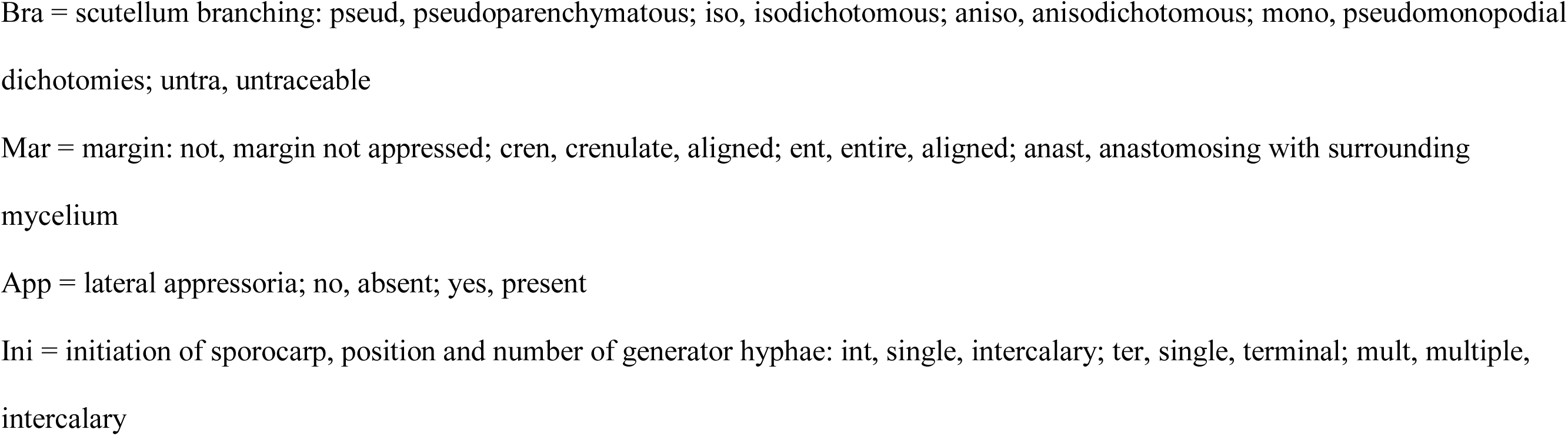
Comparison of ancestral character states at selected nodes as reconstructed from proportional likelihoods in Mesquite (unshaded rows) and from posterior probabilities in BayesTraits (shaded rows).

### Reconstruction of evolutionary origin of character states

#### Sporulation substrate, lichenization, sporocarp type

The sporulation substrate of ancestral Dothideomycetes is reconstructed with likelihood (Fig. 1; Appendix S8) as immersed in plant tissue, or with BayesTraits as being saxicolous (Appendix S8). The most recent common ancestors of thyriothecial lineages appear to have adapted to superficial growth on leaf or twig surfaces (Fig. 1; Appendix S8). In the case of Muyocropronales, adaptations to superficial growth (Appendix S8) preceded origin of thyriothecia. The ancestral Dothideomycetes and the most recent common ancestors of thyriothecial fungi are reconstructed as non-lichenized (Appendix S9), although Micropeltidaceae is nested in the Ostropomycetidae among lichenized fungi. Reconstructions suggest that Dothideomycetes initially produced apothecioid sporocarps (Appendix S10).

Thyriothecial Micropeltidaceae appear to have arisen from apothecioid ancestors while thyriothecial Asterinales, Capnodiales and Muyocopronales appear to have originated from perithecioid ancestors. In three clades, Asterotexiales Clade A, Aulographaceae, and the group of Microthyriales plus Venturiales clade, ancestors are reconstructed as thyriothecial or unresolved (Appendix S10).

#### Lower walls

Unlike most ascomata, thyriothecia usually lack a differentiated lower wall, a layer of pigmented fungal tissue that would separate sporogenous tissue from the substrate (Fig. 1; Appendix S11). Lower walls are hypothesized to be present in the ancestor of Dothideomycetes based on 100% proportional likelihood and 58% support from BayesTraits (Node 1 in Fig. 1; Appendix S11). Out of 59 thyriothecial species included here, only four have a pigmented, differentiated lower wall (Appendices S4 and S11) and ancestral species reconstructed as thyriothecial are also reconstructed as lacking a differentiated lower wall.

#### Dehiscence

The mechanism of spore release from sporocarps varies among the closest relatives of Dothideomycetes, and among thyriothecial taxa (Appendix S12). Likelihood reconstructions suggest that ancestral Dothideomycetes opened with an ostiole (Fig. 1); BayesTraits reconstructions favor opening by a regular slit and show ancestral forms with slits subsequently giving rise to ostiolate clades that diversified further (Table 3; Appendices S6, S7 and S12). A transition to irregular slits is reconstructed somewhere before the most recent common ancestor of Asterotexiales and Aulographaceae (Figs. 1 and 2; Appendix S12). A convergent transition from ostioles to irregular slits appeared along a branch nested in Capnodiales that leads to Asterinales. Developmentally, the irregular slits result from cracking between the adjacent hyphae that make up the scutellum (Fig. 2L, 2M). The precise location of the opening remains unclear until dehiscence. Slit-forming genera sometimes produce ostioles in their asexual or spermatial states (Fig. 2O). In *Aulographaceae*, and possibly also in species of *Lembosia*, the slit appears early in development as a line of lightly pigmented cells (Fig. 3), but the crack follows longitudinal walls and propagates beyond the less pigmented area. Regular slits that follow a pre-determined pattern without further propagation are uncommon among Dothideomycetes. Regular slits appear to have originated twice among thyriothecial taxa, being present in *Parmularia* in Asterinales and in *Inocyclus angularis* in Asterotexiales, but not in their inferred common ancestor (Appendix S12). Regular slits are also found in taxa producing hysterothecia and lirella (such as *Farlowiella charmichaeliana, Hysterographium fraxinii*, *Melaspileopsis* cf. *diplasiospora* (Nyl.) Ertz & Diederich *Stictographa lentiginosa* (Lyell ex Leight.) Mudd, which at maturity may look more like apothecia than thyriothecia (Appendix S12).

**Figure 2.**
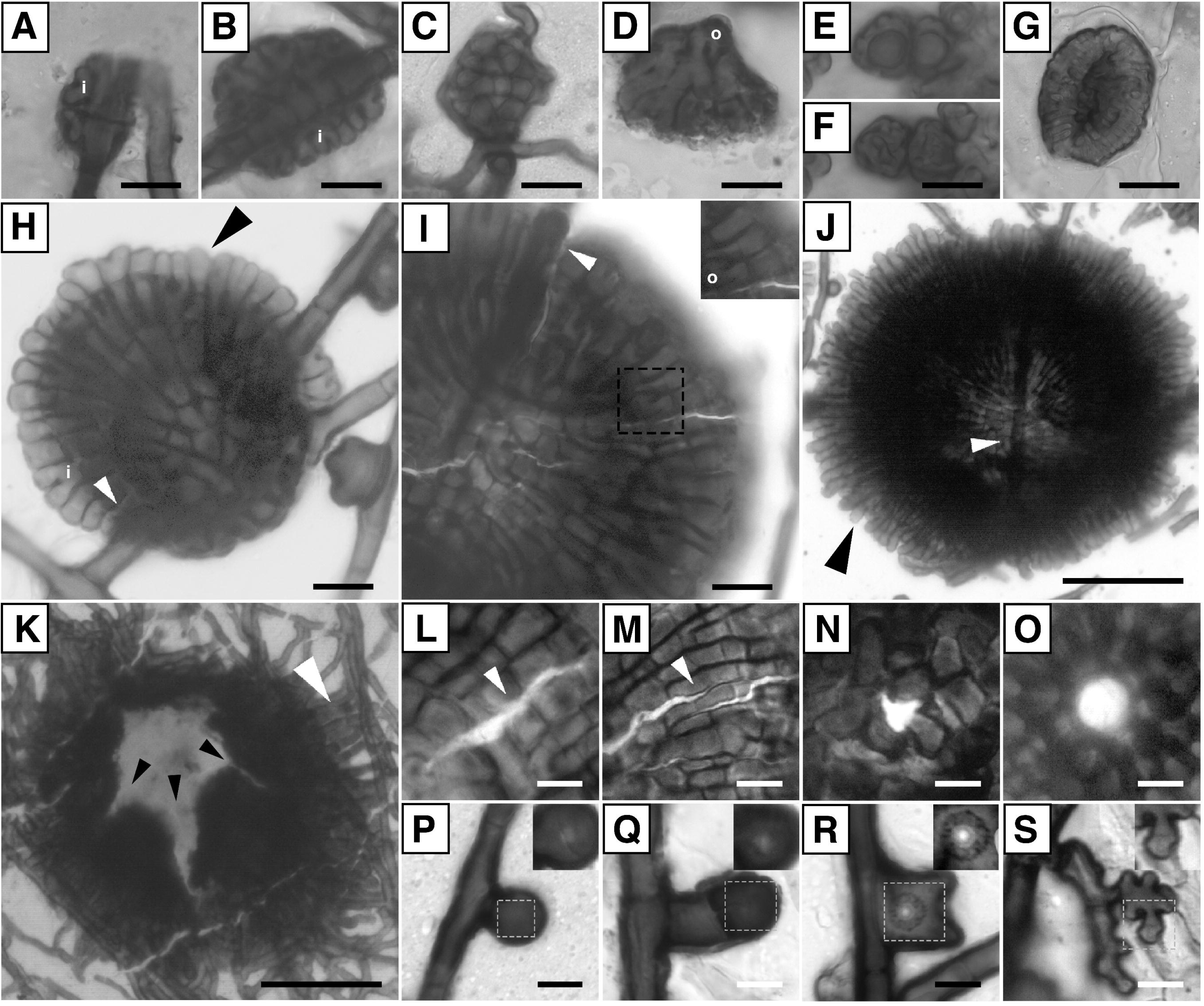
Morphological characters of Asterinales and Asterotexiales in face view. (A–J) Development. Initiation of ascomata in Asterinales and Asterotexiales usually involves intercalary septation of a superficial generator hypha but is sometimes (D) uninterpretable, (E, F) from spores, or (G) from stomata. (H–J) The highly septate generator hypha usually persists above the scutellum throughout development (white arrowhead). (A–D, H–J) The generator hypha gives rise to the scutellum through (i) the formation of new, lateral hyphal tips that branch with isotomous dichotomies, occasionally (D, I, inset) with one tip that overlaps the other (o). (H, J) The appressed scutellum margin shows lightly pigmented tips and is usually crenulate at maturity (black arrowhead). (K) Hyphae of the appressed marginal of scutella *Prillieuxina baccharidincola* instead anastomose with surrounding mycelium (white arrowhead). (I, K, L–O) Dehiscence. (I, K, L–M) Asterinales and Asterotexiales ascomata usually dehisce with irregular slits. (K) Mature scutellum with irregular radial slits, black arrowheads. (L-M) Irregular slits follow the longitudinal walls of scutellum cells (white arrowheads). (N-O) Ostiolate openings are less common. (N) Ostiole in young scutellum. (O) Asexual or spermatial stage, with evenly pigmented cells lining ostiole. (P–S) Lateral appressoria showing characteristic melanized ring at lower focal plane (insets, representing area in the dashed grey boxes). Scales (A–D, H–I) 10 µm; (E–G) 20 µm; (J–K) 50 µm; (L–S) 5 µm. (A, I, J, P, Q) *Asterina melastomatis* VIC 52822, 1, 4, 6, 2 and 2 focal planes (f.p.) respectively; (B) *Asterina chrysophylli* VIC 42823, 4 f.p.; (C, O) Asterotexiaceae sp. CBS 143813, 2 f.p. each; (D) *Hemigrapha atlantica* BR 14014, 1 f.p.; (E, F) *Asterotexis cucurbitacearum* VIC 42814, 1 f.p. each; (G, L) *Rhagadolobiopsis thelypteridis* EG156; (H, R) *Batistinula gallesiae* VIC 42514, 10 and 2 focal planes (f.p.) respectively; (K N), *Prillieuxina baccharidincola* VIC 42817; (M) Asterotexiaceae sp. UBC F33036, 11 f.p.; (S) *Asterina* sp. (VUL. 341b), 2 f.p.

**Figure 3.**
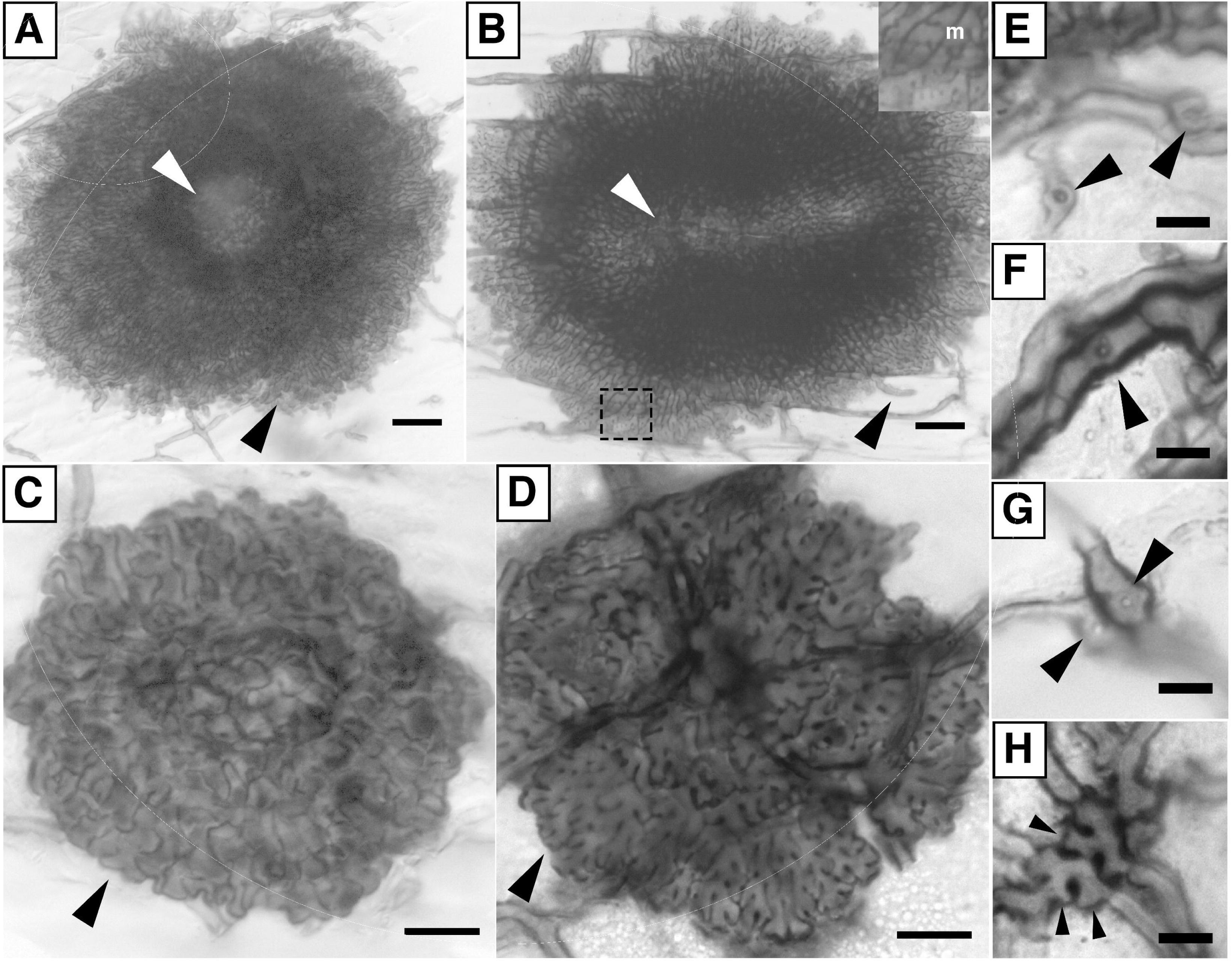
Morphological characters of Aulographaceae and *Stomiopeltis* spp. s.l. (Venturiales) in face view. (A–D) Scutella. (A) Mature *Stomiopeltis*-like scutellum. Hyphae are radiate but untraceable due to frequent overlap of neighboring hyphae. The ostiole is wide (white arrowhead) and at the distal appressed margin, hyphal tips are free (black arrowhead). (B) Mature Aulographaceae scutellum with radiate, pseudomonopodial hyphal branching. Inset (m), a close up of hyphal branching in the area in the dashed grey box. An irregular slit for dehiscence would form in the elongate, lighter area (white arrowhead). The appressed distal margin has free hyphal tips. (C) Early development of a *Stomiopeltis*-like scutellum. The appressed distal margin is irregularly crenulate (arrowhead). (D) Young Aulographaceae scutellum showing multiple, intercalary generator hyphae, pseudomonopodial hyphal branching, and an irregularly crenulate appressed margin (arrowhead). (E–G) Intercalary appressoria with a melanized ring (arrowheads). (H) Initiation of Aulographaceae ascomata; coordinated intercalary septation of multiple generator hyphae (arrowheads) gives rise to scutella. Scales, (A, B) 20 µm; (C, D) 10 µm; (E–H) 5 µm. (A, C, F) *Stomiopeltis* sp. UBC F33041, 5, 6 and 1 stacked focal planes (f.p.) respectively. (E) *Stomiopeltis* sp. UBC F33040, 1 f.p. (B) *Lembosina aulographoides* CBS 143809, 4 f.p.; (D, G, H) *Aulographum* sp. CBS 143545, 1 f.p.

Ancestral ostioles appear to have been retained in Microthyriales and Muyocopronales (Figs. 1 and 4; Appendix S12), but the mode of dehiscence and ancestral states are unknown in Zeloasperisporiales. The common ancestor of Micropeltidaceae probably released spores through an ostiolate thyriothecium which is hypothesized to be derived from the apothecium that is shared by other members of Ostropomycetidae in Lecanoromycetes (Appendix S12).

**Figure 4.**
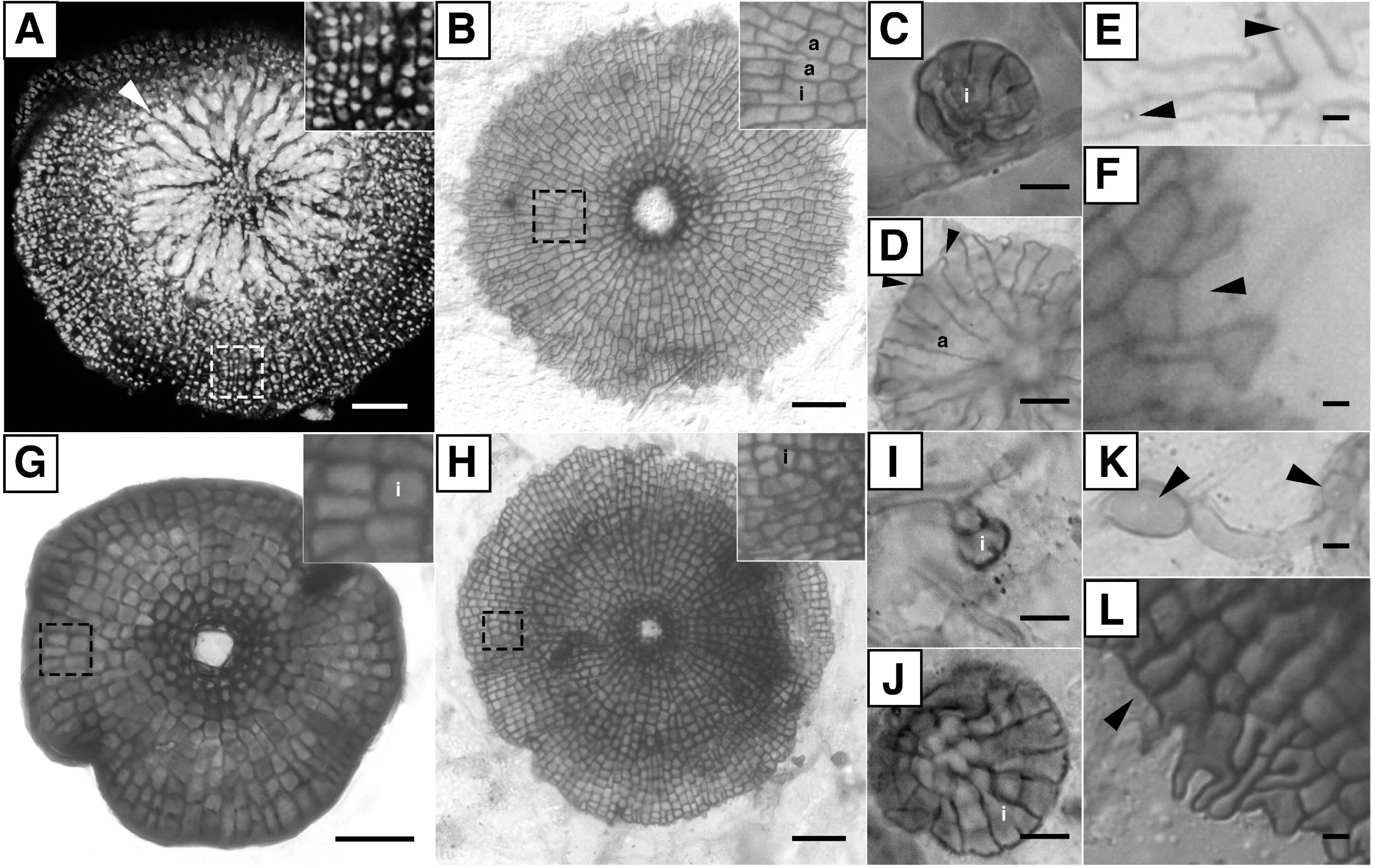
Morphological characters of Microthyriales in face view. (A) Calcofluor white staining of *Microthyrium* sp. reveals radially arranged asci (arrowhead) in a hymenium below the scutellum. The inset shows uneven fluorescence of scutellum hyphae due to dark pigments. (B, G, H) Mature scutella are radiate and insets show details of anisotomous (a) and isotomous (i) hyphal branching. Dehiscence is with an ostiole that is lined by small, darkly pigmented cells. (C, D, I, J) Early development of scutella. The appressed margins of young scutella are entire. (I) The scutellum is initiated at the tip of a lateral generator hypha. (C, D, J) New hyphal tips arise from the generator hypha with closely spaced, isotomous, dichotomous branches. (D, J) The young scutella overgrow and conceal the generator hyphae. (E, K) Intercalary appressoria in a regular (E) or swollen (K) hyphal segment, showing melanized rings (arrowheads). (F, L) Margins of mature scutella are considered to be entire if hyphal tips are blunt, at least in part (arrowheads) even if narrow portions of the tip hyphae extend further into an irregular, meandering fringe as in (B, L). Scales, (A–C) 20 µm; (D) 10 µm; (E–I) 5 µm. (A) *Microthyrium* sp., 44 focal planes (f.p.); (B–F), *Microthyrium macrosporum* CBS 143810, with 4, 3, 1, 1 and 1 focal planes (f.p.), respectively; (G), *Lichenopeltella pinophylla*, CBS 143816, 14 f.p.; (H–L), *M*. *ilicinum* CBS 143808, with 4, 1, 3 and 2 f.p. respectively.

#### Locule circumference shape

In most Ascomycota, spores are formed in a cavity called a locule (Kirk et al., 2008). In Dothideomycetes, the circumference of the locule within the ascoma or conidioma is commonly round but may be elongate. Elongate locules are scattered across the phylogeny. Thyriothecial species with elongate locules are inferred to be closely related to species with round locules (Appendix S13), contrary to predictions from the current classification (Wu et al., 2011b). For example, ascoma shape, which was used to separate the elongate *Lembosia* from circular *Asterina*, fails to predict relationships. Instead, species of both genera appear in both Asterotexiales and in Asterinales (Fig. 1; Appendix S6). In tree topology tests, constraining either *Asterina* or *Lembosia* to be monophyletic is rejected with strong support (Appendix S3).

#### Radiate development

Scutella of thyriothecia are either made up of radiating hyphae that can be traced most of the distance from their distal tips to their central point of origin (Figs. 1–5), or of hyphae that are not radiate and cannot be traced back to their origin (Figs. 1 and 5). Radiating patterns of thyriothecia are restricted to Dothideomycetes (Fig. 1; Appendices S4 and S14). They are reconstructed to have originated convergently from ancestors with non-radiating patterns. Radiate scutella appear to have arisen independently in Asterinales and in Muyocopronales (Appendix S14). One or multiple additional transitions to radiate thyriothecia may have occurred among Asterotexiales, Aulographaceae, and in the clade including Microthyriales and Zeloasperisporiales (Table 3; Appendix S14). Non-radiating thyriothecial patterns also appear to have originated convergently, being found in distantly related species in Capnodiales in Dothideomycetes, and in Micropeltidaceae in Ostropomycetidae (Fig. 1; Appendices S4 and S14).

**Figure 5.**
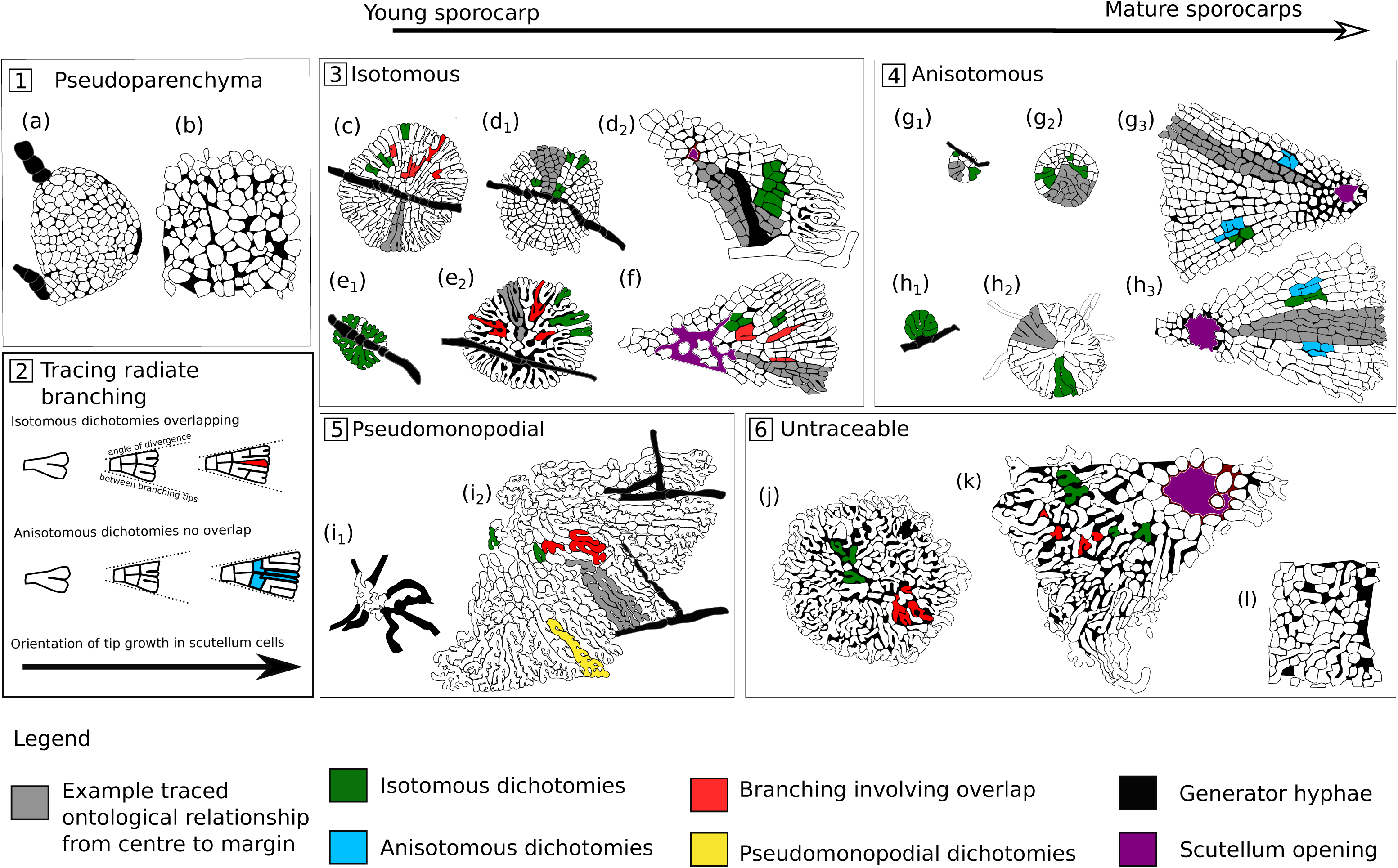
Scutellum initiation and interpretation of branching. (A) Ancestral Dothideomycetes had pseudoparenchymatous walls (a–b). (B) Radiate scutella in thyriothecia grow by successive dichotomies followed by cross-wall formation. Isotomous dichotomies produce daughter hyphae of similar width. Hyphae occasionally overlap so that one file of cells seems to vanish (top); anisotomous dichotomies produce daughter hyphae that differ in width and rarely overlap (bottom). (C) Asterotexiales/Asterinales pattern. Scutellum hyphae show isotomous dichotomies, and some hyphae overlap in some specimens (c, e) but not in all (d_1-2_) specimens. Overlap can be missing in young specimens (e_1_). Initiation of the scutellum often begins with a single intercalary generator hypha. The generator hypha persists above the scutellum at maturity (c, d_1-2_, e_1-2_). (D) Microthyriales pattern. Young scutella are initiated at the tip of a short lateral generator hypha (g_1_, h_1_). The generator hypha gives rise to scutellum hyphae that eventually overgrow it as they branch with isotomous dichotomies (g_1-2_, h_1-2_). The generator hypha is not visible in the mature scutellum (g_2-3_, h_2-3_). In mature scutella, hyphae show anisotomous and isotomous dichotomies and hyphae do not overlap (g_3_, h_3_). (E) Aulographaceae pattern. Scutella with pseudomonopodial branching (i_2_). The scutellum is usually initiated by coordinated growth of multiple adjacent, intercalary generator hyphae (i_1_). (F) *Stomiopeltis* (UBC F33041) pattern. Scutellum hyphae are generally radiate but they overlap and are irregular (j–l) and they cannot be traced to generator hypha or from the center of the scutellum to the margin. Sources of drawings: (a) *Fumiglobus pieridicola* UBC-F33205; (b) *Pleospora herbarum* UBC-F4461; (c) *Batistinula gallesiae* VIC 42514; (d) Asterotexiaceae sp. UBC F33036; (e) *Asterina chrysophylli* VIC 42823; (f) Asterotexiaceae sp. CBS 143813; (g) *Microthyrium ilicinum* CBS 143808; (h) from *M. macrosporum* CBS 143810, (i) *Aulographum* sp. CBS 143545, (j) *Stomiopeltis* sp. UBC F33041, (k) *Stomiopeltis* sp. CBS 143811 and (l) *Peltaster cerophilus* redrawn from Fig. 2M in (Medjedović et al., 2014).

#### Phylogenetic distribution of hyphal branching patterns among thyriothecial scutella

Likelihood consistently reconstructs pseudoparenchymatous sporocarp walls of polyhedral cells as the ancestral state at the base of Dothideomycetes and Lecanoromycetes, (Figs. 1 and 5, Table 3; Appendix S15). However, corresponding BayesTraits reconstructions are equivocal (Appendices S6, S7 and S15). Among thyriothecial fungi, branching patterns of the hyphae forming the scutellum wall are usually distinctive compared with the untraceable hyphae in the more widespread pseudoparenchymatous ascomatal walls of most Dothideomycetes.

Scutella with isotomous overlapping branching are reconstructed as possibly ancestral at the base of Clade A in Asterotexiales (proportional likelihood 68%, BayesTraits 52%) and for Asterinales after the divergence of *Parmularia styracis* (proportional likelihood 100%, BayesTraits 56%), with increasing support for reconstructions with this state near the tips of the tree (Table 3; Appendices S6, S7 and S15). Isotomous branching patterns in thyriothecia expand the scutellum circumference mainly by duplicating files of cells through apical, nearly equal, isotomous dichotomies, resulting first in a “Y” shaped cell with a basal septum, and then in three cells, after additional septa delimit the two arms of the “Y” as separate cells (Figs. 2A, 2B, 2H and 5B). In some cases, one of the two branches overlaps the other, growing downward and disappearing below its neighbors (Figs. 2D, 2I and 5C). Septa are sometimes aligned around part of the circumference of the scutellum, resulting in partial concentric rings of septa (cf. Fig. 3.8 “d” in Hofmann 2009). Pigmentation in older parts of the scutellum contrasts with more translucent tips at the margin of scutellum hyphae (Figs. 2H, 2J and 2K).

Scutella with pseudomonopodial branching are restricted to Aulographaceae. The common ancestor of Aulographaceae is reconstructed with pseudomonopodial branching with 99% proportional likelihood and 28% posterior probability (Table 3; Appendices S6, S7 and S15). In this growth form, the tips of dominant hyphae grow at the circumference of the expanding scutellum, sequentially producing short lateral branches (Figs. 3B inset, 3D and 5E) that overlap with neighboring hyphae. Hyphae curve as they grow, and septa only form in the oldest hyphal segments. This results in a scutellum of irregular, multi-lobed cells (Fig. 5E).

Distal tips at the appressed margin are rounded, and uniform in diameter (Figs. 3B, 3D and 3H). Within Aulographaceae, the construction of scutella is similar in *Lembosina* (Fig. 3B) and *Aulographum* (Fig. 3D; Appendix S4).

Scutellum hyphae in some thyriothecial species in Venturiales and Capnodiales in Dothideomycetes, and in all Micropeltidaceae in Lecanoromycetes are “untraceable” (Figs.1, 3A, and 3C). Scutella may show some evidence of radiate construction, but details of hyphal branching are obscured by dark pigmentation, or curved or meandering scutellum hyphae that overlap and obscure one another, resulting in a “*textura intricata”* or “*textura epidermoidea”* (Kirk et al., 2008; Figs. 3A, 3C and 5F).

In Microthyriales, scutellum hyphae usually branch dichotomously without overlap. While their branching is usually isotomous, anisotomous branching also occurs, producing hyphal tips of unequal width (Figs. 4B inset, 4D, 5B and 5D). A septum sometimes forms at the base of only one of the two new hyphal tips, resulting in a roughly ’L’ shaped proximal cell (Figs. 4B and 5D; cf. Fig. 15E Wu et al. 2011). The anisotomous expansion is reconstructed in the common ancestor of Zeloasperisporiales with 81% proportional likelihood (Fig. 1) but only 21% posterior probability from BayesTraits (Appendices S6 and S7). Septa are angular (Figs. 4B, 4G, 4H and 5D) and constrictions at septa are infrequent, resulting in trapezoidal to nearly rectangular cells with sharp angles. Partially aligned septa in adjacent hyphae sometimes result in a pattern of incomplete concentric rings. Early stages of thyriothecial development may be isotomous in Microthyriaceae (Figs. 4C, 4I, 4J and 5D) making the distinction between anisotomy and isotomy most visible in fully grown structures (Fig. 5D).

#### Scutellum margins

Thyriothecia are unusual in that the margins of each scutellum are pressed to the surface of a host--a leaf or a lichen--and the characters of the hyphae at the scutellum margin differ across clades. Other kinds of sporocarps lack a direct homolog to the scutellum margin, because their outer walls curve up from the substrate. In Asterotexiales, Asterinales, Aulographaceae, in species of *Stomiopeltis*, and in Muyocopronales, the scutellum margins often look crenulate or finely scalloped, because the dome-shaped hyphal tips bulge out around the circumference (Figs. 2H, 2J, 3C and 3D; Appendix S16). In some *Stomiopeltis*-like species (Fig. 3A), tips of hyphae are free; we interpret this as a variant of a crenulate margin (Table 2). In other species, for example in *Prillieuxina baccharidincola* (Fig. 2K), the scutellum hyphae anastomose with other superficial hyphae on the leaf surface, so that the margin of the scutellum is poorly defined and continuous with the surface mycelium. In Microthyriales and Zeloasperisporiales, scutellum margins are often entire, the scutellum hyphae ending in truncate tips aligned around the circumference (Figs. 4D, 4F and 4L). In *Microthyrium*, the hyphal tips of scutella that have finished radial expansion will narrow and continue to grow, forming a short fringe at the scutellum periphery (Figs. 4B and 4L). Fringe hyphae appear to be more tightly appressed to the host surface than central portions of scutella.

#### Lateral appressoria

Appressoria are widely distributed across thyriothecium-forming Dothideomycetes and vary in their morphology and position. Conspicuous, pigmented, lateral appressoria are only produced by Asterinales and Asterotexiales (Figs. 2P–2S; Appendices S4 and S17). The appressoria are usually unicellular (Figs. 2P and 2R) but in some species they may be two-celled (Figs. 2Q and 2S) (Hofmann, 2009, Figs. 3.8; 3.14; 3.21; 3.24; 3.31; 3.56). Lateral appressoria may be swollen, lobed or minimally differentiated, and they branch from surface hyphae. We coded thyriothecial taxa of Asterinales and Asterotexiales that lack appressoria or produce them below the scutellum rather than on surface hyphae as absent for this character (Table 1).

Other kinds of appressoria appear in other thyriothecial clades. All Aulographaceae examined have intercalary appressoria on swollen cells with a conspicuous melanized ring (Fig. 3G), and strains isolated in pure culture form an appressorium regularly in each segment of somatic hyphae. Species of *Stomiopeltis* (Venturiales) form infrequent, intercalary appressoria (Figs. 3E and 3F). The hyphae bearing the appressoria may be swollen (Figs. 3E and 3G).

Inconspicuous, hyaline, intercalary appressoria are produced by some Microthyriales (Figs. 4E and 4K). BayesTraits reconstructs lateral appressoria as present in the most recent shared ancestor of Asterotexiales, and in the most recent shared ancestor of Asterinales, while likelihood reconstructs them as ancestral only for lineages that have diverged more recently from one another (Table 3; Appendix S17).

#### Initiation of thyriothecia

Development of thyriothecia begins with distinctive, lineage-specific patterns of coordinated hyphal growth and septation in the species where observation was possible. We observed different developmental stages of 13 taxa (Table 1) and analyzed development for 14 additional taxa from published illustrations (Appendix S4). Otherwise, data on development is scarce and ancestral state reconstructions are equivocal where data are missing (Appendices S4, S6, S7 and S18).

In Asterinales and Asterotexiales a single “generator hypha” (Hofmann, 2009) initiates the thyriothecia by forming multiple, closely spaced intercalary septa (Figs. 2A–2C, 2H–2J and 5C). Each of the delimited cells then gives rise to transverse, adjacent, dichotomizing hyphal tips that together form the scutellum. The generator hypha remains above the dorsal surface of the mature scutella and is visible even in mature thyriothecia (Figs. 2I–2J and 5C). Ascomata in one species of Asterotexiales, *Rhagadolobiopsis thelypteridis*, are initiated directly over host stomata, as shown by Guatimosim et al. (2014) and in Fig. 2G here; in *Asterotexis cucurbitacearum,* they are initiated directly from ascospores (Figs. 2E and 2F) (shown previously by Guerrero et al., 2011; Guatimosim et al., 2015).

Of all the Aulographaceae observed, three species of *Lembosina* and one species of *Aulographum* initiate their ascomata with coordinated development from multiple generator hyphae. Adjacent intercalary portions of the multiple generator hyphae develop abutting, transverse branches that contribute to forming a single scutellum (Figs. 3H and 5E). At a later stage, radially arranged hyphae form pseudomonopodial branching units that cover the upper surface of the hyphal aggregate (Fig. 3D).

In contrast to the widespread intercalary origin among Asterinales and Asterotexiales, all Microthyriales examined initiate their ascomata at the tip of a generator hypha (terminal), located on a short lateral branch of a surface hypha (Figs. 4C, 4I and 5D). The generator hypha does not become highly septate, but instead its tip gives rise to a succession of dichotomously branched, closely appressed hyphal tips that develop into a scutellum. The scutellum in Microthyriales overgrows the generator hypha; the generator hypha can no longer be seen in older scutella (Figs. 4B, 4G and 4H).

#### Case studies using thyriothecial fossils

Morphological characters of thyriothecial scutella and mycelia offer a range of variation, allowing phylogenetically informed interpretations of published fossils. We looked for the oldest geological occurrence of thyriothecia, and also selected two other thyriothecial fossils that have a reasonably complete set of morphological characters (Fig. 6; Appendix S4). Each phylogenetic analysis that included a fossil yielded more than one equally parsimonious tree, given the morphological dataset and the rDNA likelihood constraint tree. All trees from the morphological datasets were the same length, 232 steps, whether or not one of the fossils was included.

**Figure 6.**
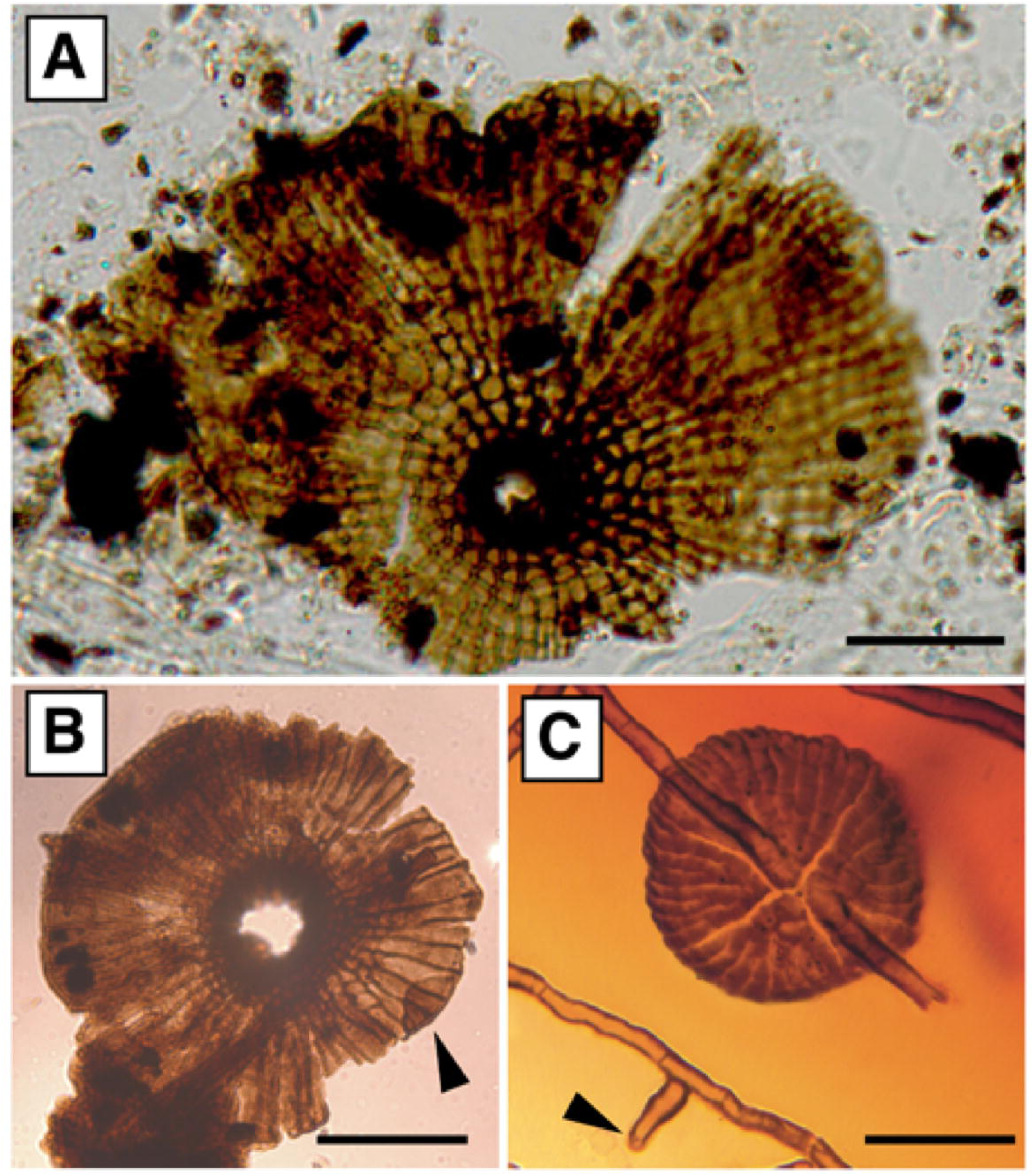
Reproduction of fossil illustrations used in our case studies. (A) Triassic dispersed “fungal thalli,” courtesy of Mishra et al. (2018), reproduced with the permission of Springer Nature; B.S.I.P. Slide No.15524, J36-4. (B) Early Eocene dispersed *Trichothyrites setifer* with lower wall (arrowhead) seen through the translucent scutellum, reproduced with permission from Monga et al. (2015); B.S.I.P. slide no. 15297, U41/3. (C) Middle Eocene *Asterina eocenica* on cuticle of *Chrysobalanus* sp. described in Dilcher (1965), illustration courtesy Steven Manchester, reproduced from Taylor et al. (2015), with permission from Elsevier; Florida Museum of Natural history, l.f. 87. Scales, 20µm.

*Triassic dispersed scutellum–*The oldest evidence of a radiate fungal scutellum is a fossil from the Early Triassic of India (Induan, ∼251 Ma) referred to as a “fungal thallus” but not identified to species or genus. The fossil was from a palynological sample that was rich in land plant cuticles and spores (Mishra et al. 2018, Fig. 8j). It shares five characters with extant thyriothecial taxa (Appendix S4). Like extant radial thyriothecia, it is made up of hypha-like filaments that are closely appressed along their sides. The filaments are regularly septate into trapezoidal cells that form partial, concentric rings. Some of the filaments dichotomize. A central circular opening appears to be an ostiole surrounded by a ring of very dark cells. Where the scutellum margin is in focus, it appears to be crenulate (Fig. 6A; Appendix S4).

We found 28 equally parsimonious positions for this fossil relative to the constraint tree. Based on its five coded characters, the fossil could cluster with thyriothecial taxa in Asterinales, Asterotexiales, or Microthyriales. Missing character state data from some extant taxa increased the number of possible, equally parsimonious positions for the fossil to include positions among the asexual taxa of Zeloasperisporiales. A strict consensus of the 28 trees resulted in a polytomy of 45 lineages, all stemming from the most recent ancestor of all Dothideomycetes (Fig. 1; Appendix S19).

#### Microthyriaceous fossil

*Trichothyrites setifer* (Cookson) Saxena & Misra, a fossil from the early Eocene (Ypresian, 56–47.8 Ma) sediments of India is illustrated by Monga et al. (2015) (Fig. 6B). While this specimen was from dispersed material, rather than in situ on a leaf surface, it shares seven characters with extant *Lichenopeltella pinophylla* and six characters with other thyriothecial Microthyriales (Appendix S4). It has a circular, multicellular thallus. The radiate scutellum has an entire margin and shows the presence of a lower wall (Fig. 6B, arrowhead). The scutellum branching pattern is mostly isotomous, but also shows anisotomous dichotomies. Its central ostiole is surrounded by papillate cells. Papillate ostioles are uncommon but occur in *Chaetothyriothecium elegans* and *L. pinophylla*. However, *C. elegans* lacks a differentiated lower wall.

We found 16 equally parsimonious trees resolving *T. setifer* in our phylogeny of living fungi. The strict consensus position of this fossil suggests it shares affinities with the clade containing Microthyriales and Zeloasperisporiales (Fig. 1; Appendix S20). Among the 16 reconstructions, *T. setifer* is alternatively placed in Microthyriales; as sister to Microthyriales (10/16); as sister to asexual taxa in Zeloasperisporiales (that are missing data for characters of the sporocarp) (3/16); as the sister to the perithecioid Natipusillales (2/16); or as the sister to all these lineages (Fig. 1; Appendix S20).

#### Asterina fossil

The early Eocene (56-48 Ma) *Asterina eocenica* was found on leaves of *Chrysobalanus* L. (Fagaceae Dumort.) and consists of circular, multicellular thalli and mycelia that are preserved in different states of development (Appendix S4), allowing comparison with extant Asterinales and Asterotexiales (Dilcher, 1965). The fossil thyriothecium lacks a lower wall, and dehiscence was by means of irregular slits. The pattern of branching of the scutellum is isotomous, with overlapping hyphae, and the margin is crenulate, as in living *Asterina.* The sporocarp was initiated with an intercalary generator hypha (Fig. 6C). The mycelium bears lateral appressoria.

A strict consensus of the 23 equally parsimonious trees places *A. eocenica* in a polytomy at the base of Dothideomycetes (Appendix S21). *Asterina eocenica* shares all 11 morphological characters with three extant taxa in Asterinales and with nine in Asterotexiales (Fig. 6C; Appendix S4). Consistent with the shared characters, 18 of the individual phylogenies show *A. eocenica* within or as sister to Asterotexiales (Appendix S22), and five show it in, or as sister to Asterinales (Appendix S23).

## DISCUSSION

Results from our molecular phylogeny of 320 species including 59 species of thyriothecial taxa combined with critical analysis of 11 morphological characters of extant Dothideomycetes offer grounds for cautious optimism for incorporating thyriothecial fossil data into the broader picture of fungal evolution. Ancestral state reconstructions show that convergent evolution of thyriothecia in Dothideomycetes likely began among fungi that produced pseudoparenchymatous-walled sporocarps on surfaces of plant cuticles. Our reconstructions suggest that radiate scutella arose at least three times within epiphyllous Dothideomycetes. As radiate thyriothecial forms evolved, patterns of hyphal branching and septation diversified, giving rise to distinctive scutella. As the following fossil case studies show, a phylogenetic approach takes into account morphological variation within orders and convergence among orders. We apply the approach to fossils but the same methods could be used to evaluate possible relationships among extant taxa that are difficult to sample using molecular techniques.

### Case studies: interpreting characters of fossils

*A dispersed scutellum as evidence of Triassic Dothideomycetes*—Among our sampling of extant taxa, radiate scutella are restricted to Dothideomycetes. Further, ancestral state reconstructions show radiate thyriothecia as a derived character state that arose convergently, but only among epiphyllous Dothideomycetes. Consistent with results from the parsimony analysis, the combination of an ostiole and a scutellum with dichotomous branching that is found in the fossil is ancestral for Asterinales, Microthyriales, and possibly for Asterotexiales. The fossil could represent a member of any of these groups.

Although it cannot be classified more precisely, parsimony analysis shows that the scutellum- like “fungal thallus” of Mishra et al. provides evidence of Dothideomycetes. The fossil from the Early Triassic (∼251 Ma) (2018, Fig. 8j) likely represents the oldest evidence of thyriothecia.

### Fossil evidence of Microthyriales

Monga (2015, Pl.2 Fig.18) illustrates the fossil *Trichothyrites setifer*. Like the “fungal thallus” discussed above, *T. setifer* is from a sample rich in land plant material, but its substrate is unknown. *Trichothyrites* specimens (synonym of *Notothyrites*, see Kalgutkar and Jansonius, 2000), interpreted as members of Trichothyriaceae have frequently been reported in the fossil record throughout the Tertiary (Kalgutkar and Jansonius, 2000). Although classified in Microthyriaceae, extant *Lichenopeltella* shares multiple characters with Trichothyriaceae Theiss., including a scutellum with isotomous and anisotomous dichotomies; a scutellum margin that is entire, a papillate ostiole, and a differentiated lower wall that can be seen through the translucent scutellum (Spooner and Kirk, 1990). Thyriothecia with a papillate ostiole and a lower wall are characteristic of Trichothyriaceae. Anisotomous dichotomies were not present in *Lichenopeltella pinophylla*, the only extant taxon we sampled with this morphology, but they do occur in the genus and are illustrated for *Lichenopeltella arctomiae* Pérez-Ort. & T. Sprib. (Fig. 1 in Pérez-Ortega and Spribille, 2009).

Although *T. setifer* is more similar to *Lichenopeltella* than any other taxa in our dataset, some of its shared characters including ostiolate thyriothecia are reconstructed as ancestral in the larger clade encompassing Zeloasperisporiales, Natipusillales and Microthyriales. Three of six taxa in Zeloasperisporiales are only known from asexual states and are missing data for the sporocarp. A differentiated lower wall also occurs in non-thyriothecial ascomata of Natipusillales. As a result of plesiomorphic characters and missing data, equally most parsimonious placements of *T. setifer* were: in Microthyriales; as the sister lineage to Microthyriales, Natipusillales or Zeloasperisporiales; and as sister to the clade including all of these taxa.

At present, *T. setifer* (Monga et al., 2015), from the early Eocene (56–47.8 Ma), represents the oldest convincing evidence of the clade including Microthyriales and Venturiales. Investigating Paleocene deposits may very well yield further and older evidence of the divergence of Trichothyriaceae. For more formal testing, the molecular phylogenetic positions of extant taxa of *Actinopeltis* Höhn., *Lichenopeltella*, *Trichothyrium* Speg., and *Trichothyrina* (Petr.) Petr. should be established using molecular data, and their morphology analyzed for comparison with the fossils.

Asterina*-like fossils could represent either Asterinales or Asterotexiales*—Contrary to our expectations, combining close zmorphological observations and the molecular phylogeny revealed no clade-specific morphological differences between Asterinales and Asterotexiales.

Guatimosim et al. (2015) showed that Asterinales and Asterotexiales both include taxa with scutellum dehiscence by irregular cracks, and superficial hyphae bearing lateral appressoria. The orders share crenulate scutellum margins. *Lembosia* species (characterized by elongate thyriothecial locules) occur nested among both Asterinales and Asterotexiales species with round locules.

Developmental and anatomical characters for Asterinales and Asterotexiales were sometimes illustrated in great detail in early works (Arnaud, 1918; Doidge, 1919; Viégas, 1944). More recently, Hofmann’s (2009) thesis contains elegant drawings from diverse species, documenting the common intercalary origin of thyriothecia, starting with septation in a generator hypha. At the time of her thesis, the distinction between Asterinales and Asterotexiales had yet to be discovered and Hofmann lacked sequence data to link her observations to clades. Our analysis of species of Asterinales and Asterotexiales shows that development and scutellum branching as Hofmann described are common to both orders.

Liu et al. (2017) interpreted Asterotexiales as “Asterinales sensu stricto” and suggested that the Asterinales, represented by a clade of six species and their LSU rDNA sequences, all from Brazil, should be considered “Asterinales sensu lato”. However, the six Brazilian Asterinales species that were sequenced by Guatimosim et al. (2015) include *Asterina melastomatis* (from the epitype specimen of the Asterinales), which means that the ordinal name applies to their clade. If a single sequence had been involved, or if the disputed Brazilian Asterinales were from a clade of fungi commonly amplified from leaf surfaces in other studies, contamination by non-target DNAs might seem a possible explanation for the presence of morphologically similar species in phylogenetically divergent groups. However, the Brazilian sequences give congruent results for species in five genera of thyriothecium forming taxa: *Batistinula*, *Prillieuxina*, *Parmularia*, *Asterina*, and *Lembosia*. In a survey of *Eucalyptus* L’Hér. spp. Crous et al. (2019) found additional thyriothecial species of *Thyrinula* Petr. & Syd. to be closely related to asexual species of *Blastacervulus.* Our analysis shows these two genera to be in Asterinales, and future detailed analysis of their scutellum morphology will provide data useful in reconstructing ancestral character states of Asterinales. Other thyriothecial species from leaves of *Eucalyptus* spp. lack sequence data but are diverse in their scutellum morphology (Swart, 1986) and some of these may also represent Asterinales. In the absence of any contradictory evidence, the sequences from Guatimosim et al. (2015) and from Crous et al. (2019) must be assumed to come from their target fungi.

It is possible that more molecular data may resolve Asterinales and Asterotexiales as sister groups, consistent with their morphological similarities. Tree topology tests did not rule out a sister relationship between the two orders. Although the orders consistently appear unrelated in published phylogenies, their separation never receives strong bootstrap support or high posterior probabilities (Guatimosim et al., 2015; Firmino, 2016; Liu et al., 2017 and Fig. 1). With cultures now available representing Asterotexiales (Asterotexiaceae sp.2 CBS 143813) and Asterinales (*Blastacervulus robbenensis* CBS 12478) it should soon be possible to expand available sequences beyond the ∼1 kb of 28S ribosomal DNA per taxon that currently limits phylogenetic resolution. If Asterinales and Asterotexiales are not sister groups, their thyriothecial characters represent surprising convergence. If Asterinales and Asterotexiales are sister groups, a phylogenetic analysis of their morphological characters, including dehiscence type, scutellum branching and pattern of initiation of sporocarp, should likely allow unambiguous assignment of fossils to their clade.

*Asterina eocenica* from the Middle Eocene (48.6-37.2 Ma) of western Tennessee (Dilcher, 1965; Elsik and Dilcher, 1974) is the oldest fossil that combines a convincing Asterinales/Asterotexiales suite of characters. The exquisitely preserved scutella of *A. eocenica* show clear isotomous dichotomies with overlap (Figs. 64 and 65 in Dilcher, 1965). This places the fossil with clades of thyriothecia in which hyphae branch dichotomously at their apices to produce radial scutella (Farr, 1969; Reynolds and Gilbert, 2005; Hofmann and Piepenbring, 2011; Guatimosim et al., 2015). Dichotomous branching is infrequent among fungi, where branching is usually initiated some micrometers behind the hyphal apex. *Asterina eocenica* has nine out of 11 characters shared by Asterinales and Asterotexiales. The development of thyriothecia of *A. eocenica* is typical of Asterinales and Asterotexiales, with initiation by closely spaced septa forming in a generator hypha. The generator hypha persists on the dorsal side of the scutellum (Figs. 59, 60, 63, 64 and 65 in Dilcher, 1965). Like Asterinales and Asterotexiales, *A. eocenica* has slit-like dehiscence (Figs. 63 and 65 in Dilcher 1965), an undifferentiated lower wall (Figs. 61, 63, 65 and 66 in Dilcher, 1965), and some elongate, one-celled tapering lateral hyphal projections described as appressoria (Figs. 57 and 59 in Dilcher, 1965). In their otherwise detailed and useful review of fossils, Samarakoon et al. (2019) assign *A. eocenica* to crown group Asterinales (=Asterotexiales) and with a minimum age as Paleocene, 54 Ma. The Paleocene age appears to be an error because it is inconsistent with an analysis by Elsik and Dilcher (1974), which assigned a more recent Middle Eocene age to the fossil’s source material from the Lawrence Clay Pit. None of the characters in *A. eocenica* justify positioning it in the crown group of either order as opposed to a stem relationship of either order, and as the parsimony analysis shows, it could equally well represent Asterinales and Asterotexiales. A strong implication of the close similarity of Asterinales and Asterotexiales is that even a beautiful, complete fossil like *A. eocenica* cannot be assigned to one order versus the other.

## CONCLUSION

Much cryptic diversity in Dothideomycetes consists of minute leaf dwelling fungi (Arx and Müller, 1954; Müller and von Arx, 1962; von Arx and Müller, 1975). As our analysis shows, epiphyllous fungi together with lichenized, lichenicolous and aquatic fungi are all relevant to the evolution of thyriothecia, but they have been studied by different mycologists who sequenced different loci for phylogenetic analysis (Dhanasekaran et al., 2006; Miadlikowska et al., 2006; Etayo, 2010). The sequencing of the 5’ end of the LSU was common across most studies, but choices of other loci varied. We anticipate that sequencing of more Dothideomycetes genomes, including some from the new isolates from this study from cultures available through the Westerdijk Institute, will contribute to improved resolution of relationships among extant taxa.

Our molecular phylogeny of thyriothecial taxa and their closest relatives leads to testable predictions about the sequence of early events in their evolutionary history. A remaining limitation of this study is that we could include only a small fraction of living fly-speck species. The fidelity of association of characters with extant lineages should be tested across a broader range of taxa using morphological and molecular phylogenetic analysis. Undoubtedly, further study will help reveal convergence and variation, and perhaps additional distinctive synapomorphies. Our analyses suggest that analyzing and coding scutellum characters can link fossils on leaf cuticle to phylogenetic lineages, offering rewarding insights into fungal evolution through geological time. However limited fossilized characters may be, documenting equivalent anatomical features in extant taxa helps communicate the significance of fossil morphologies, increasing their value for paleomycology.

## ACKNOWLEDGEMENTS

This research was enabled in part by support provided by WestGrid (www.westgrid.ca), Compute Canada Calcul Canada (www.computecanada.ca) and CIPRES Science Gateway (www.phylo.org). The authors also wish to express their gratitude to UBC Bioimaging facilities for use of their fluorescence microscope, UBC InterLibrary Loan service for providing crucial literature, Robert Lücking for gift to UBC Herbarium of lichenized taxa in genera *Microtheliopsis* and *Asterothyrium*, OTS and Conagebio, and more specifically Francisco Campos Rivera and Bernal Mattarita Carranza for allowing collection of tropical thyriothecium forming fungi in the research station of La Selva in Costa Rica. Funding for this research was provided by a Natural Sciences and Engineering Research Council of Canada Discovery Grant RGPIN-2016–03746 to MLB.

## SUPPORTING INFORMATION

Additional Supporting Information may be found online in the supporting information section at the end of the article

**Appendix S1**. Isolates included in this study; names in bold indicate newly contributed collections and voucher/strain numbers beginning with ’CBS’ and followed by ’*’ represent cultures newly deposited in the Westerdijk Institute. When LSU and SSU were derived from two different collections, bold indicates the voucher/strain name that is used in phylogenies.

Accessions labeled ’JGI’ refer to sequences extracted from JGI genomes project, for which no GenBank accession exists.

**Appendix S2**. JGI sequence data retrieved from MycoCosm portal.

**Appendix S3**. Results from CONSEL for six topologies tested against most likely tree.

**Appendix S4**. Matrix of characters used in ancestral character state reconstructions and in parsimony analyses of fossils.

**Appendix S5**. Consensus tree from Mr. Bayes analysis with four runs of eight chains each, running 160 million generations and sampling every 5000 trees, from which we discarded 50% of the samples as burn-in. Posterior probabilities mapped on branches, clades are labeled and ordered as in Fig. 1.

**Appendix S6**. Maximum likelihood phylogeny labeled with codes for nodes reconstructed in ancestral character state analyses. Taxa forming thyriothecia are indicated with pink diamonds. Most likely tree obtained out of 4000 independent ML searches of a 4552 bp alignment of LSU and SSU rDNA data for 320 taxa, with bootstrap support > 70% in red and a posterior probability > 0.95.

**Appendix S7**. Detailed output of ancestral state reconstruction for each of 208 nodes as proportional likelihoods (white rows) or posterior probabilities (grey rows). Node labels in column 1 are designated in Fig. 1. Nodes labeled in Column 2 are designated in the phylogeny in Appendix S6.

**Appendix S8–S18**. Ancestral character state reconstructions for 11 morphological characters, given the most likely DNA sequence topology. Reconstructions from MK1 analysis are shown above branches as proportional likelihoods in pie charts with red outlines. Reconstructions from BayesTraits are shown below branches as posterior probabilities in pie charts with black outlines. In each tree, names in grey indicate taxa that could not be coded for the character, with ancestral states that could not be reconstructed. Bootstrap support >0.7 is reported in red above branches and posterior probabilities >0.95 are in black below branches.

**Appendix S8.** Ancestral character state reconstructions for substrate (Sub).

**Appendix S9.** Ancestral character states reconstruction (Lic), as lichenized or non-lichenized.

**Appendix S10.** Ancestral character state reconstruction for sporocarp type (Spo).

**Appendix S11.** Ancestral character state reconstruction for differentiated or undifferentiated lower wall of sporocarp (Low).

**Appendix S12.** Ancestral character state reconstruction for sporocarp dehiscence (Deh).

**Appendix S13.** Ancestral character state reconstruction for locule circumference (Loc).

**Appendix S14.** Ancestral character state reconstruction for presence or absence of a radiate sporocarp (Rad).

**Appendix S15.** Ancestral character state reconstruction for scutellum branching (Bra).

**Appendix S16.** Ancestral character state reconstruction for margin of sporocarp (Mar).

**Appendix S17.** Ancestral character state reconstruction for the presence or absence of lateral appressoria on superficial mycelium (App).

**Appendix S18.** Ancestral character state reconstruction for Sporocarp initiation (Ini): terminal on generator hypha, intercalary on generator hypha, from spore, from host stomata or from multiple generator hyphae.

**Appendix S19.** Consensus of 28 equally parsimonious phylogenetic positions of the Triassic “fungal thallus” from India described in Mishra et al. (2018).

**Appendix S20.** Consensus of 16 equally parsimonious phylogenetic positions of *Trichothyrites setifer* from the Eocene of India described in Monga et al. (2015).

**Appendix S21.** Consensus of 23 equally parsimonious phylogenetic positions of *Asterina eocenica* from the Eocene of USA, described in Dilcher (1965).

**Appendix S22.** Consensus of the 18 equally parsimonious phylogenetic trees placing *Asterina eocenica* with taxa in Asterotexiales.

**Appendix S23.** Consensus of the 5 equally parsimonious phylogenetic trees placing *Asterina eocenica* with taxa in Asterinales.

## LITERATURE CITED

Arnaud, G. 1917. Sur la famille des microthyriacées. Compte Rendu de l’Académie des Sciences Paris 164: 574–577.

Arnaud, G. 1918. Les Asterinées. Annales de l’École Nationale d’Agriculture de Montpellier 16: 1–288.

Arnaud, G. 1925. Les Asterinées. IVe partie. Annales de l’École National d’Agriculture de Montpellier, série 10 5: 643–722.

Arx, J. A. v., and E. Müller. 1954. Die Gattungen der amerosporen Pyrenomyceten. Beiträge zur Kryptogamenflora der Schweiz 11: 1–434.

Batista, A. C. 1959. Monografia dos fungos Micropeltaceae. Publicações do Instituto de Micologia da Universidade do Recife 56: 1–519.

Batista, A. C., J. L. Bezerra, T. T. Barros, and F. B. Leal. 1969. Sobre um novo gênero de Microthyriaceae da Nova Caledônia. Instituto de Micologia, Universidade do Recife 637: 1–11.

Bercovici, A., A. Hadley, and U. Villanueva-Amadoz. 2009. Improving depth of field resolution for palynological photomicrography. Palaeontologia Electronica 12: 1–12.

Clements, F. E., and C. L. Shear. 1931. The genera of fungi, second ed. New York.

Crous, P. W., M. J. Wingfield, R. Cheewangkoon, A. J. Carnegie, T. I. Burgess, B. A. Summerell, J. Edwards, et al. 2019. Foliar pathogens of eucalypts. Studies in Mycology 94: 125– 298.

Darriba, D., G. L. Taboada, R. Doallo, and D. Posada. 2012. jModelTest 2: more models, new heuristics and parallel computing. Nature Methods 9: 772–772.

Dhanasekaran, V., R. Jeewon, and K. D. Hyde. 2006. Molecular taxonomy, origins and evolution of freshwater ascomycetes. Fungal Diversity 23: 351–390.

Dilcher, D. L. 1965. Epiphyllous fungi from Eocene deposits in western Tennessee, U.S.A. Palaeontographica Abteilung B 116: 1–54.

Doidge, E. M. 1919. South African Microthyriaceae. Transactions of the Royal Society of South Africa 8: 235–282.

Doidge, E. M. 1942. A revision of the South African Microthyriaceae. Bothalia 4: 273–420.

Ducomet, V. 1907. Recherches sur le développement de quelques champignons parasites à thalle subcuticulaire. Thèse doctorale, Faculté des sciences de Paris, Rennes, France.

Ellis, J. P. 1976. British *Microthyrium* species and similar fungi. Transactions of the British Mycological Society 67: 381–394.

Elsik, W. C. 1978. Classification and geologic history of the microthyriaceous fungi. IVth International Palynological Conference (1976–77), Lucknow, India.

Elsik, W. C., and D. L. Dilcher. 1974. Palynology and age of clays exposed in Lawrence clay pits, Henry County, Tennessee. Palaeontographica Abteilung B: 65–87.

Ertz, D., and P. Diederich. 2015. Dismantling Melaspileaceae: a first phylogenetic study of *Buelliella*, *Hemigrapha*, *Karschia*, *Labrocarpon* and *Melaspilea*. Fungal Diversity 71: 141–164.

Etayo, J. 2010. Lichenicolous fungi from the western Pyrenees. V. Three new ascomycetes. Opuscula Philolichenum 8: 131–139.

Farr, M. L. 1969. Some “black mildew”, “sooty mold”, and “fly-speck” fungi and their hyperparasites from Dominica. Canadian Journal of Botany 47: 369–381.

Felsenstein, J. 1985. Confidence limits on phylogenies: an approach using the bootstrap. Evolution 39: 783–791.

Firmino, A. L. 2016. Taxonomy, phylogeny and biogeography of Asterinales on native forest species from Atlantic Forest and Cerrado. Tese doutourado, Universidade Federal de Viçosa, Viçosa, Minas Gerais, Brazil.

Gäumann, E. A. 1952. The fungi. A description of their morphological features and evolutionary development. Hafner Publishing Company, New York.

Grigoriev, I. V., R. Nikitin, S. Haridas, A. Kuo, R. Ohm, R. Otillar, R. Riley, et al. 2013. MycoCosm portal: gearing up for 1000 fungal genomes. Nucleic acids research 42: D699–D704.

Guatimosim, E., H. J. Pinto, R. W. Barreto, and J. Prado. 2014. Rhagadolobiopsis, a new genus of Parmulariaceae from Brazil with a description of the ontogeny of its ascomata. Mycologia 106: 276–281.

Guatimosim, E., A. Firmino, J. Bezerra, O. Pereira, R. Barreto, and P. Crous. 2015. Towards a phylogenetic reappraisal of Parmulariaceae and Asterinaceae (Dothideomycetes). Persoonia 35: 230–241.

Guerrero, Y., T. A. Hofmann, C. Williams, M. Thines, and M. Piepenbring. 2011. *Asterotexis cucurbitacearum*, a poorly known pathogen of Cucurbitaceae new to Costa Rica, Grenada and Panama. Mycology 2: 87–90.

Hernández-Restrepo, M., J. D. P. Bezerra, Y. P. Tan, N. P. Wiederhold, P. W. Crous, J. Guarro, and J. Gené. 2019. Re-evaluation of *Mycoleptodiscus* species and morphologically similar fungi. Persoonia 42: 205–227.

Hofmann, T., R. Kirschner, and M. Piepenbring. 2010. Phylogenetic relationships and new records of Asterinaceae (Dothideomycetes) from Panama. Fungal Diversity 43: 39–53.

Hofmann, T. A. 2009. Plant parasitic Asterinaceae and Microthyriaceae from the Neotropics (Panama). Ph.D., Johann Wolfgang Goethe-University, Frankfurt, Germany.

Hofmann, T. A., and M. Piepenbring. 2006. New records and host plants of fly-speck fungi from Panama. Fungal Diversity 22: 55–70.

Hofmann, T. A., and M. Piepenbring. 2011. Biodiversity of *Asterina* species on neotropical host plants: New species and records from Panama. Mycologia 103: 1284–1301.

Holm, K., and L. Holm. 1991. Ascomycetes on *Myrica gale* in Sweden. Nordic Journal of Botany 11: 675–687.

Hongsanan, S., and D. K. Hyde. 2017. Phylogenetic placement of Micropeltidaceae. Mycosphere 8: 1930–1942.

Hongsanan, S., Y.-M. Li, J.-K. Liu, T. Hofmann, M. Piepenbring, J. D. Bhat, S. Boonmee, et al. 2014. Revision of genera in Asterinales. Fungal Diversity 68: 1–68.

Hosagoudar, V., and J. Kapoor. 1985. New technique of mounting meliolaceous fungi. Indian Phytopathology 38: 548–549.

Hughes, S. J. 1976. Sooty moulds. Mycologia 68: 693–820.

James, T. Y., F. Kauff, C. L. Schoch, P. B. Matheny, V. Hofstetter, C. J. Cox, G. Celio, et al. 2006. Reconstructing the early evolution of Fungi using a six-gene phylogeny. Nature 443: 818– 822.

Kalgutkar, R. M., and J. Jansonius. 2000. Synopsis of fossil fungal spores, mycelia and fructifications. AASP Contributions Series, 351. American Association of Stratigraphic Palynologists Foundation.

Katoh, K., and D. M. Standley. 2013. MAFFT Multiple sequence alignment software Version 7: Improvements in performance and usability. Molecular Biology and Evolution 30: 772–780.

Kirk, P. M., P. F. Cannon, D. W. Minter, and J. A. Stalpers. 2008. Ainsworth & Bisby’s Dictionary of the Fungi, 10th ed. CABI Europe, UK.

Kumar, M. 1996. Palynostratigraphy and paleoecology of Early Eocene palynoflora of Rajpardi lignite, Bharuch District, Gujarat. Palaeobotanist 43: 110–121.

Lee, S. B., and J. W. Taylor. 1990. Isolation of DNA from fungal mycelia and single spores. In M. A. Innis, D. H. Gelfand, J. J. Sninsky, and T. J. White [eds.], PCR Protocols: A Guide to Methods and Applications, 282–287. Academic Press, San Diego.

Lewis, P. O. 2001. A likelihood approach to estimating phylogeny from discrete morphological character data. Systematic Biology 50: 913–925.

Mapook, A., K. Hyde, S. Hongsanan, C. Phukhamsakda, J. Li, and S. Boonmee. 2016. Palawaniaceae fam. nov., a new family (Dothideomycetes, Ascomycota) to accommodate Palawania species and their evolutionary time estimates. Mycosphere 7: 1732–1745.

Medjedovi, A., J. Frank, H.-J. Schroers, B. Oertel, and J. C. Batzer. 2014. Peltaster cerophilus is a new species of the apple sooty blotch complex from Europe. Mycologia 106: 525–536.

Miadlikowska, J., F. Kauff, V. Hofstetter, E. Fraker, M. Grube, J. Hafellner, V. Reeb, et al. 2006. New insights into classification and evolution of the Lecanoromycetes (Pezizomycotina, Ascomycota) from phylogenetic analyses of three ribosomal RNA- and two protein-coding genes. Mycologia 98: 1088–1103.

Miller, M. A., W. Pfeiffer, and T. Schwartz. 2010. Creating the CIPRES Science Gateway for inference of large phylogenetic trees. Gateway Computing Environments Workshop (GCE), New Orleans, LA: 1–8.

Mishra, S., N. Aggarwal, and N. Jha. 2018. Palaeoenvironmental change across the Permian- Triassic boundary inferred from palynomorph assemblages (Godavari Graben, south India). Palaeobiodiversity and Palaeoenvironments 98: 177–204.

Mitchell, A. 2015. Collecting in collections: a PCR strategy and primer set for DNA barcoding of decades-old dried museum specimens. Molecular Ecology Resources 15: 1102–1111.

Monga, P., M. Kumar, V. Prasad, and Y. Joshi. 2015. Palynostratigraphy, palynofacies and depositional environment of a lignite-bearing succession at Surkha Mine, Cambay Basin, north- western India. Acta Palaeobotanica 55: 183–207.

Müller, E., and J. A. von Arx. 1962. Die Gattungen der didymosporen Pyrenomyceten. Beiträge zur Kryptogamenflora der Schweiz 11: 1–922.

Pagel, M., and A. Meade. 2007. BayesTraits v.2 Computer program and documentation available at http://www.evolution.rdg.ac.uk/BayesTraits.html.

Parbery, D., and R. Emmett. 1977. Hypotheses regarding appressoria, spores, survival and phylogeny in parasitic fungi. Revue de Mycologie 41: 429–447.

Phipps, C. J., and W. C. Rember. 2004. Epiphyllous fungi from the Miocene of Clarkia, Idaho: reproductive structures. Review of Palaeobotany and Palynology 129: 67–79.

Reynolds, D. R., and G. S. Gilbert. 2005. Epifoliar fungi from Queensland, Australia. Australian Systematic Botany 18: 265–289.

Ronquist, F., M. Teslenko, P. Van Der Mark, D. L. Ayres, A. Darling, S. Höhna, B. Larget, et al. 2012. MrBayes 3.2: efficient Bayesian phylogenetic inference and model choice across a large model space. Systematic Biology 61: 539–542.

Shimodaira, H. 2002. An approximately unbiased test of phylogenetic tree selection. Systematic Biology 51: 492–508.

Shimodaira, H., and M. Hasegawa. 2001. CONSEL: for assessing the confidence of phylogenetic tree selection. Bioinformatics 17: 1246–1247.

Spooner, B. M., and P. M. Kirk. 1990. Observations on some genera of Trichothyriaceae. Mycological Research 94: 223–230.

Stamatakis, A. 2014. RAxML version 8: a tool for phylogenetic analysis and post-analysis of large phylogenies. Bioinformatics 30: 1312–1313.

Stevens, F. L., and M. H. Ryan. 1939. The Microthyriaceae. The University of Illinois press, Urbana.

Swart, H. J. 1986. Australian leaf-inhabiting fungi XXII. Microthyrium-like fungi on Eucalyptus. Transactions of the British Mycological Society 87: 81–91.

Taylor, T. N., M. Krings, and E. L. Taylor. 2015. Fossil Fungi, 1st ed. Elsevier/Academic Press Inc., San Diego.

Viégas, A. P. 1944. Algun fungos do Brasil II Ascomycetos. Bragantia 4: 1–392.

von Arx, J. A., and E. Müller. 1975. A re-evalutation of bitunicate ascomycetes with keys to families and genera. Studies in Mycology 9: 1–159.

Warren, D. L., A. J. Geneva, and R. Lanfear. 2017. RWTY (R We There Yet): An R package for examining convergence of bayesian phylogenetic analyses. Molecular Biology and Evolution 34: 1016–1020.

Wijayawardene, N., P. Crous, P. Kirk, D. Hawksworth, S. Boonmee, U. Braun, D.-Q. Dai, et al. 2014. Naming and outline of Dothideomycetes–2014 including proposals for the protection or suppression of generic names. Fungal Diversity 69: 1–55.

Wu, H., K. D. Hyde, and H. Chen. 2011a. Studies on Microthyriaceae: placement of *Actinomyxa*, *Asteritea*, *Cirsosina*, *Polystomellina* and *Stegothyrium*. *Cryptogamie*, Mycologie 32: 3–12.

Wu, H., Q. Tian, W.-J. Li, and K. Hyde. 2014. A reappraisal of Microthyriaceae. Phytotaxa 176: 201–212.

Wu, H.-X., C. L. Schoch, S. Boonmee, A. H. Bahkali, P. Chomnunti, and K. D. Hyde. 2011b. A reappraisal of Microthyriaceae. Fungal Diversity 51: 189–248.

