## Supplemental Data 1 for "Character evolution of modern fly-speck fungi and implications for interpreting thyriothecial fossils"

### Appendix S1. Isolates included in this study

| **Order** | **Taxa** | **Voucher/Strain** | **GenBank accession – SSU** | **GenBank accession - LSU** |
| --- | --- | --- | --- | --- |
| Abrothallales Pérez-Ort. & Suija | *Abrothallus acetabuli* Diederich | SPO308 | KF816215.1 | KF816232.1 |
| Abrothallales | *Abrothallus buellianus* De Not. | SPO303 | KF816217.1 | KF816234.1 |
| Abrothallales | *Abrothallus cladoniae* R. Sant. & D. Hawksw. | AB53 | KF816220.1 | KF816228.1 |
| Abrothallales | *Abrothallus parmeliarum* (Sommerf.) Arnold | AB36 | KF816222.1 | KF816229.1 |
| Abrothallales | *Abrothallus parmotrematis* Diederich | AB1 | KF816225.1 | KF816231.1 |
| Abrothallales | *Abrothallus secedens* Wedin & R. Sant. | SPO305 | KF816216.1 | KF816236.1 |
| Abrothallales | *Abrothallus suecicus* (Kirschst.) Nordin | AB56 | - | KF816226.1 |
| Abrothallales | *Abrothallus usneae* Rabenh. | AB20 | KF816223.1 | - |
| Abrothallales | *Lichenoconium lecanorae* (Jaap) D. Hawksw. | strain JL382-10 | - | HQ174263.1 |
| Abrothallales | *Abrothallus hypotrachynae* Etayo & Diederich | SPO302 | KF816218.1 | KF816233.1 |
| Acrospermales Minter, Peredo & A.T. Watson | *Acrospermum compressum* Tode | M151 | EU940012.1 | EU940084.1 |
| Acrospermales | *Acrospermum graminum* Lib. | M152 | EU940013.1 | EU940085.1 |
| Acrospermales | *Acrospermum adeanum* Höhn. | M133 | EU940031.1 | EU940104.1 |
| Arthoniales Henssen ex D. Hawksw. & O.E. Erikss. | *Arthonia dispersa* (Schrad.) Nyl. | UPSC 2583 | AY571381.1 | AY571381.1 |
| Arthoniales | *Chrysothrix candelaris* (L.) J.R. Laundon | UPS Frisch 11/Se45 | - | KF707640.1 |
| Arthoniales | *Combea mollusca* (Ach.) Nyl. | Tehler 7725 | AY571380.1 | AY571382.1 |
| Arthoniales | *Dendrographa leucophaea* (Tuck.) Darb. | Ornduff 10070 Duke | AF279381.1 | AF279382.1 |
| Arthoniales | *Lecanographa amylacea* (Ehrh. ex Pers.) Egea & Torrente | UPS Thor 26176 | - | KF707639.1 |
| Arthoniales | *Lichinella iodopulchra* (Couderc ex Croz.) P.P. Moreno & Egea | AFTOL-ID 896 | - | DQ782916.1 |
| Arthoniales | *Melarthonis piceae* Frisch & G. Thor | UPS Thor 25995 | - | KJ851080.1 |
| Arthoniales | *Opegrapha dolomitica* (Arnold) Clauzade & Cl. Roux ex Torrente & Egea | AFTOL-ID 993 | DQ883706.1 | - |
| Arthoniales | *Reichlingia zwackhii* (Sandst.) Frisch & G. Thor | UPS Thor 11/3 | - | KF707637.1 |
| Arthoniales | *Roccella fuciformis* (L.) DC. | AFTOL-ID 126 | AY584678.1 | AY584654.1 |
| Arthoniales | *Roccellographa cretacea* J. Steiner | AFTOL-ID 93 | DQ883705.1 | DQ883696.1 |
| Arthoniales | *Schismatomma decolorans* (Erichsen) Clauzade & Vězda | DUKE 0047570 Ertz 5003 (BR) | NG_013155.1 | NG_027622.1 |
| Arthoniales | *Simonyella variegata* J. Steiner | AFTOL-ID 80 | AY584669.1 | - |
| Arthoniales | *Alyxoria varia* (Pers.) Ertz & Tehler | - | AF138853.1 | **EU704103.1** |
| Asterinales M.E. Barr ex D. Hawksw. & O.E. Erikss. | *Asterina chrysophylli* Henn. | VIC 42823 | - | KP143738.1 |
| Asterinales | *Asterina melastomatis* Lév. | VIC 42822 | - | MK251541 |
| Asterinales | *Batistinula gallesiae* Arx | VIC 42514 | - | KM111255.1 |
| Asterinales | *Blastacervulus eucalypti* H.J. Swart | CBS 124759 | - | GQ303302.1 |
| Asterinales | *Blastacervulus eucalyptorum* Crous | CPC 29450 | - | KY173484.1 |
| Asterinales | *Blastacervulus robbenensis* (Crous et al.) Crous | CBS 124780 | - | HM628777.1 |
| Asterinales | *Lembosia abaxialis* Firmino & R.W. Barreto | VIC 42825 | - | KP143737.1 |
| Asterinales | *Parmularia styracis* Lév. | VIC 42587 | - | KP143730.1 |
| Asterinales | *Prillieuxina baccharidincola* (Rehm) Petr. | VIC 42817 | - | KP143735.1 |
| Asterinales | *Thyrinula eucalypti* (Cooke & Massee) H.J. Swart | CPC 12986 | - | HM535600.1 |
| Asterotexiales Firmino, O.L. Pereira & Crous | *Asterina* (Lév.) sp. | MFLU13-0619 | - | KM386978.1 |
| Asterotexiales | *Asterina cestricola* (R.W. Ryan) Hosag. & T.K. Abraham | TH 591 | GU586209.1 | GU586215.1 |
| Asterotexiales | *Asterina cynometrae* Hongsanan & K.D. Hyde | MFLU 13-0373 | - | KX845436.1 |
| Asterotexiales | *Asterina fuchsiae* Syd. | TH 590 | GU586210.1 | GU586216.1 |
| Asterotexiales | *Asterina phenacis* Syd. | TH 589 | GU586211.1 | GU586217.1 |
| Asterotexiales | *Asterina siphocampyli* Syd. | M-0141060 (PMA) | - | HQ701140.1 |
| Asterotexiales | *Asterina weinmanniae* Syd. | TH592 | GU586212.1 | GU586218.1 |
| Asterotexiales | *Asterina zanthoxyli* W. Yamam. | TH 561 | GU586213.1 | GU586219.1 |
| Asterotexiales | **Asterotexiaceae [Firmino, O.L. Pereira & Crous] sp. 1** | UBC-F33036 | - | MG844156 |
| Asterotexiales | **Asterotexiaceae sp. 2** | CBS 143813 * | - | MG844162 |
| Asterotexiales | *Asterotexis cucurbitacearum* | PMA M-0141224 | - | HQ610510.1 |
| Asterotexiales | *Asterotexis cucurbitacearum* (Rehm) Arx | VIC 42814 | - | KP143734.1 |
| Asterotexiales | *Buelliella minimula* (Tuck.) Fink | Lendemer 42273 (NY) | - | KX244961.1 |
| Asterotexiales | *Buelliella physciicola* Poelt & Hafellner | Ertz 19173 (BR) | - | KP456148.1 |
| Asterotexiales | *Buelliella poetschii* Hafellner | Ertz 18116 (BR) | - | KP456150.1 |
| Asterotexiales | *Discopycnothyrium palmae* Hongsanan & K.D. Hyde | MFLU13-0485 | - | KM386979.1 |
| Asterotexiales | *Hemigrapha atlantica* Diederich & Wedin | Ertz 14014 (BR) | - | KP456151.1 |
| Asterotexiales | *Inocyclus angularis* Guatim. & R.W. Barreto | VIC 39747 | - | KP143731.1 |
| Asterotexiales | *Karschia cezannei* Ertz & Diederich | Ertz 19186(BR) | - | KP456154.1 |
| Asterotexiales | *Karschia talcophila* (Ach.) Körb. | Diederich 16749 | - | KP456155.1 |
| Asterotexiales | *Labrocarpon canariense* (D. Hawksw.) Etayo & Pérez-Ort. | Ertz 16907 (BR) | - | KP456158.1 |
| Asterotexiales | *Lembosia albersii* Henn. | MFLU13-0377 | - | KM386982.1 |
| Asterotexiales | *Lembosia xyliae* X.Y. Zeng, T., C. Wen & K.D. Hyde | MFLU14-0004 | - | KT283685.1 |
| Asterotexiales | *Mahanteshomyces* Hosag. & C.K. Biju sp. | TH 588 | GU586214.1 | GU586220.1 |
| Asterotexiales | *Melaspilea lekae* Brackel & Kalb | Ertz 17325 (BR) | - | KP456162.1 |
| Asterotexiales | *Melaspileopsis* cf. *diplasiospora* (Nyl.) Ertz & Diederich | Ertz 16625 | - | KP456166.1 |
| Asterotexiales | *Morenoina calamicola* S. Konta & K.D. Hyde |  | KY511427 | KY511424 |
| Asterotexiales | *Mycosphaerella pneumatophorae* Kohlm. | AFTOL-ID 762 | - | FJ176856.1 |
| Asterotexiales | *Stictographa lentiginosa* (Lyell ex Leight.) Mudd | van den Boom 47621 | - | KP456171.1 |
| Asterotexiales | *Taeniolella hawksworthiana* Heuchert, Ertz & Common | Common 9199B (BR) | - | KX244970.1 |
| Asterotexiales | *Taeniolella punctata* M.S. Christ. & D. Hawksw. | Ertz 17390 (BR) | - | KX244972.1 |
| Asterotexiales | *Taeniolella pyrenulae* Heuchert & Diederich | Diederich 17075 | - | KX244976.1 |
| Asterotexiales | *Taeniolella* S. Hughes sp. | Ertz 11026 (BR) | - | KX244979.1 |
| Asterotexiales | *Taeniolella toruloides* Heuchert & Diederich | Diederich 17048 | - | KX244978.1 |
| Asterotexiales | *Rhagadolobiopsis thelypteridis* Guatim. & R.W. Barreto | EG 156 | - | KC171177.1 |
| Aulographaceae Luttr. ex P.M. Kirk, P.F. Cannon & J.C. David | *Aulographum hederae* | CBS 113979 | JGI | JGI |
| Aulographaceae | ***Aulographum* Lib. sp.** | CBS 143545 ***** | MG844146 | MG844158 |
| Aulographaceae | ***Lembosina aulographoides* E. Bommer, M. Rousseau & Sacc.) Theiss.** | CBS 143809 * | MG844145 | MG844157 |
| Aulographaceae | ***Lembosina* Theiss. sp.** | CBS 143815 * | - | MG844165 |
| Aulographaceae | ***Lembosina* sp.** | CBS 144007 * | - | MG844164 |
| Aulographaceae | *Aulographum hederae* Lib. | MFLUCC13-0001 | - | KM386981.1 |
| Botryosphaeriales C.L. Schoch, Crous & Shoemaker | *Botryosphaeria dothidea* (Moug.) Ces. & De Not. | CBS 115476 | DQ677998.1 | NG_027577.1 |
| Botryosphaeriales | *Kellermania anomala* (Cooke) Höhn. | CBS 132218 | KF766259.1 | NG_042700.1 |
| Botryosphaeriales | *Kellermania dasylirionicola* Minnis & A.H. Kenn. | CBS 131720 | KF766262.1 | NG_042703.1 |
| Botryosphaeriales | *Neofusicoccum ribis* (Slippers, Crous & M.J. Wingf.) Crous, Slippers & A.J.L. Phillips | AFTOL-ID 1232 | DQ678000.1 | DQ678053.1 |
| Botryosphaeriales | *Kellermania yuccifoliorum* A.W. Ramaley | CBS 131726 | KF766271.1 | NG_042713.1 |
| Caliciales Bessey | *Physcia aipolia* (Ehrh. ex Humb.) Fürnr. | AFTOL-ID 84 | DQ782876.1 | DQ782904.1 |
| Capnodiales Woron. | *Acidomyces richmondensis* B.J. Baker, M.A. Lutz, S.C. Dawson, P.L. Bond & Banfield | C2 | JGI | JGI |
| Capnodiales | *Arthrocatena tenebrio* Egidi & Selbmann | CCFEE 5413 | GU250342.1 | GU250385.1 |
| Capnodiales | *Aulographina pinorum* (Desm.) Arx & E. Müll. | CBS 174.90 | GU296138.1 | GU301802.1 |
| Capnodiales | *Capnobotryella renispora* Sugiy. | CBS 214.90 | AF006723.1 | GU214398.1 |
| Capnodiales | *Capnodium citri* Berk. & Desm. | CBS 451.66 | GU296177.1 | AY004337.1 |
| Capnodiales | *Capnodium coffeae* Pat. | CBS 147.52 | DQ247808.1 | GU214400.1 |
| Capnodiales | *Catenulostroma chromoblastomycosum* Crous & U. Braun | CBS 597.97 | GU214516.1 | EU019251.2 |
| Capnodiales | *Cercospora zebrina* Pass. | (SSU) STE-U 3955, (LSU) **CBS 118790** | AY251104.2 | JQ739815.1 |
| Capnodiales | *Cladosporium bruhnei* Linder | CPC 5101 | AY251096.2 | GU214408.1 |
| Capnodiales | *Comminutispora agavacearum* A.W. Ramaley | CBS 619.95 | Y18699.1 | EU981286.1 |
| Capnodiales | *Conidiocarpus caucasicus* Woron. | GUMH937 | KC833051.1 | KC833050.1 |
| Capnodiales | *Davidiella tassiana* (De Not.) Crous & U. Braun | DAOM 196248 | JN939022.1 | JN938886.1 |
| Capnodiales | *Elasticomyces elasticus* Zucconi & Selbmann | CCFEE 5320 | GU250333.1 | GU250376.1 |
| Capnodiales | *Fumiglobus pieridicola* T. Bose | UBC F23788 | KC833053.1 | KC833052.1 |
| Capnodiales | *Houjia yanglingensis* G.Y. Sun & Crous | YHJN13 | - | GQ433631.1 |
| Capnodiales | *Johansonia chapadensis* Crous, R.W. Barreto, Alfenas & R.F. Alfenas | CBS H-20484 | - | HQ423450.1 |
| Capnodiales | *Leptoxyphium fumago* (Woron.) R.C. Srivast. | CBS 123.26 | GU214535.1 | GU301831.1 |
| Capnodiales | *Microcyclospora pomicola* J. Frank, B. Oertel, Schroers & Crous | CPC 16173 | GU570559.1 | GU570551.1 |
| Capnodiales | *Mycosphaerella latebrosa* (Cooke) J. Schröt. | CBS 687.94 | GU214546.1 | GU214444.1 |
| Capnodiales | *Passalora fulva* (Cooke) U. Braun & Crous | (SSU) STE-U 3688, (LSU) **CBS 119.46** | AY251109.2 | DQ008163.2 |
| Capnodiales | *Peltaster fructicola* Eric M. Johnson, T.B. Sutton & Hodges | strain 11157 | KF550926.1 | JN573665.1 |
| Capnodiales | *Penidiella columbiana* Crous & U. Braun | CBS 486.80 | GU214565.1 | EU019274.2 |
| Capnodiales | *Phaeophleospora atkinsonii* (Syd.) Pennycook & McKenzie | CBS 124565 | JN938701.1 | GU214462.1 |
| Capnodiales | *Phaeotheca fissurella* Sigler, Tsuneda & J.W. Carmich. | CBS 520.89 | Y18697.1 | GU117900.1 |
| Capnodiales | *Phragmocapnias asiaticus* Chomnunti & K.D. Hyde | MFLUCC10-0062 18S | JN832597.1 | JN832612.1 |
| Capnodiales | *Piedraia hortae* Fonseca & Leão | CBS 480.64 | AY016349.1 | AY016366.1 |
| Capnodiales | *Racodium rupestre* Pers. | L346 | EU048575.1 | EU048583.1 |
| Capnodiales | *Rasutoria tsugae* (Dearn.) M.E. Barr | EF1147 | EF114730.1 | EF114705.1 |
| Capnodiales | *Readeriella mirabilis* Syd. & P. Syd. | CBS 116293 | EU754110.2 | EU019291.2 |
| Capnodiales | *Recurvomyces mirabilis* Selbmann & de Hoog | CCFEE 5264 | GU250329.1 | GU250372.1 |
| Capnodiales | *Schizothyrium pomi* (Mont. & Fr.) Arx | CBS 228.57 | - | KF902007.1 |
| Capnodiales | *Scorias spongiosa* (Schwein.) Fr. | AFTOL-ID 1594 | DQ678024.1 | DQ678075.1 |
| Capnodiales | *Sphaerulina polyspora* F.A. Wolf | (SSU) STE-U 4301, (LSU) **CBS 354.29** | AY251095.1 | GU214501.1 |
| Capnodiales | *Stomiopeltis versicolor* (Desm.) Arx | GA3 23C2b | - | FJ147163.1 |
| Capnodiales | *Teratosphaeria stellenboschiana* (Crous) Crous | CPC 10886 | GU214583.1 | EU019295.2 |
| Capnodiales | *Uwebraunia communis* (Crous & Mansilla) Crous | CBS 110747 | NG_016521.1 | GQ852589.1 |
| Capnodiales | *Xenomeris juniperi* (Dearn.) M.E. Barr & E. Müll. | isolate="xejucf" | EF114734.1 | EF114709.1 |
| Capnodiales | *Zasmidium anthuriicola* (U. Braun & C.F. Hill) Crous & U. Braun | CBS 118742 | GU214595.1 | GQ852732.1 |
| Capnodiales | *Mycosphaerella walkeri* R.F. Park & Keane | CPC 11252 | GU214593.1 | GU214500.1 |
| Chaetothyriales M.E. Barr | *Capronia pilosella* (P. Karst.) E. Müll., Petrini, P.J. Fisher, Samuels & Rossman | AFTOL-ID 657 | DQ823106.1 | DQ823099.1 |
| Chaetothyriales | *Ceramothyrium carniolicum* (Rehm) Petr. | CBS 175.95 | AF346418.1 | AY004339.1 |
| Chaetothyriales | *Ceramothyrium podocarpi* Crous | CPC 19826 | - | KC005795.1 |
| Chaetothyriales | *Chaetothyrium agathis* Hongsanan & K.D. Hyde | MFLUCC 12 C0113 | - | KP744480.1 |
| Chaetothyriales | *Chaetothyrium brischoficola* Chomnunti & K.D. Hyde | MFLU(CC)10-0012 | - | HQ895836.1 |
| Chaetothyriales | *Exophiala dermatitidis* (Kano) de Hoog | AFTOL-ID 668 | DQ823107.1 | DQ823100.1 |
| Chaetothyriales | *Exophiala pisciphila* McGinnis & Ajello | AFTOL-ID 669 | DQ823108.1 | DQ823101.1 |
| Chaetothyriales | *Phaeosaccardinula dendrocalami* Chomnunti & K.D. Hyde | IFRDCC 2663 | - | KF667246.1 |
| Chaetothyriales | *Phaeosaccardinula ficus* Chomnunti & K.D. Hyde | MFLU(CC)10-0009 | - | HQ895837.1 |
| Chaetothyriales | *Sarcinomyces petricola* Wollenz. & de Hoog | CBS 101157 | FJ358318.1 | FJ358249.1 |
| Chaetothyriales | *Ceramothyrium linnaeae* (Dearn.) S. Hughes | UPSC 2646 | AF022715.1 | - |
| Dothideales Lindau | *Aureobasidium subglaciale* (Zalar, de Hoog & Gunde-Cim.) Zalar, de Hoog & Gunde-Cim. | EXF-2481 | JGI | JGI |
| Dothideales | *Dothidea insculpta* Wallr. | CBS 189.58 | DQ247810.1 | NG_027643.1 |
| Dothideales | *Stylodothis puccinioides* (DC.) Arx & E. Müll. | CBS 193.58 | NG_013130.1 | NG_027594.1 |
| Dothideales | *Sydowia polyspora* (Bref. & Tavel) E. Müll. | AFTOL-ID 178 | AY544718.1 | AY544675.1 |
| Dothideales | *Dothidea sambuci* (Pers.) Fr. | AFTOL-ID 274 | AY544739.1 | AY544681.1 |
| Dyfrolomycetales K.L. Pang, K.D. Hyde & E.B.G. Jones | *Dyfrolomyces rhizophorae* (K.D. Hyde) K.D. Hyde, K.L. Pang, Alias, Suetrong & E.B.G. Jones | JK 5349A | GU479766.1 | GU479799.1 |
| Dyfrolomycetales | *Dyfrolomyces tiomanensis* K.L. Pang, Alias, K.D. Hyde, Suetrong & E.B.G. Jones | NTOU3636 | KC692155.1 | KC692156.1 |
| Erysiphales E. Warming | *Erysiphe mori* (I. Miyake) U. Braun & S. Takam. | MUMHS77 | AB033484.2 | AB022418.1 |
| Eurotiales G.W. Martin ex Benny & Kimbr. | *Aspergillus fumigatus* Fresen. | (SSU) **JCM1738** (LSU) HP044 | AB008401.1 | KT323254.1 |
| Eurotiales | *Aspergillus niger* Tiegh. | ATCC 1015 | JGI | JGI |
| Eurotiales | *Monascus purpureus* Went | AFTOL-ID 426 | DQ782881.1 | DQ782908.1 |
| Eurotiales | *Emericella nidulans* (Eidam) G. Winter | ATCC 10074 | U77377.1 | KC146369.1 |
| Helotiales Nannf. ex Korf & Lizoň | *Cudoniella clavus* Alb. & Schwein.) Dennis | AFTOL-ID 166 | DQ470992.1 | DQ470944.1 |
| Helotiales | *Mollisia cinerea* (Batsch) P. Karst. | AFTOL-ID 76 | DQ470990.1 | DQ470942.1 |
| Helotiales | *Monilinia laxa* (Aderh. & Ruhland) Honey | CBS 122031 | AY544714.1 | NG_027608.1 |
| Hysteriales Lindau | *Hysterobrevium constrictum* (N. Amano) E. Boehm & C.L. Schoch | SMH 5211.1 | GU397361.1 | GQ221905.2 |
| Hysteriales | *Psiloglonium araucanum* (Speg.) E. Boehm, Marinc. & C.L. Schoch | CBS 112412 | FJ161133.2 | FJ161172.2 |
| Hysteriales | *Psiloglonium clavisporum* (Seaver) E. Boehm, C.L. Schoch & Spatafora | GKM L172A | GU323192.1 | GU323204.1 |
| Hysteriales | *Hysterobrevium smilacis* (Schwein.) E. Boehm & C.L. Schoch | CBS 114601 | FJ161135.2 | FJ161174.2 |
| Lecanorales Nannf. | *Cladonia caroliniana* | AFTOL-ID 3 | AY584664.1 | AY584640.1 |
| Lecanorales | *Lecanora hybocarpa* (Tuck.) Brodo | AFTOL-ID 639 | DQ782883.1 | DQ782910.1 |
| Licheno-stigmatales Ertz, Diederich & Lawrey | *Cf. Arthoniales sp.* | GU250325.1 | GU250325.1 | - |
| Licheno-stigmatales | *Dothideomycetes sp.* | TRN 213 | GU324008.1 | - |
| Licheno-stigmatales | *Lichenostigma maureri* Hafellner | Diederich 17326 | - | KF176953.1 |
| Licheno-stigmatales | *Phaeococcomycetaceae* McGinnis & Schell sp. | TRN 529 | GU324016.1 | GU323987.1 |
| Licheno-stigmatales | *Phaeococcomycetaceae* sp. | TRN 456 | GU324015.1 | GU323986.1 |
| Licheno-stigmatales | *Phaeococcomycetaceae* sp. | TRN 452 | GU324014.1 | GU323985.1 |
| Licheno-stigmatales | ***Seuratia millardetii* (Racib.) Meeker** | UBC-F33043 | MG844150 | MG844163 |
| Licheno-stigmatales | *Teratosphaeriaceae* Crous & U. Braun 2007 sp. | D007 09 | GU250359.1 | GU250402.1 |
| Microthyriales G. Arnaud | *Chaetothyriothecium elegans* Hongsanan & K.D. Hyde | CPC 21375 | - | KF268420.1 |
| Microthyriales | *Heliocephala gracilis* (R.F. Castañeda) R.F. Castañeda & Unter. | MUCL 41200 | HQ333479.1 | HQ333479.1 |
| Microthyriales | ***Lichenopeltella pinophylla* (Höhn.) P.M. Kirk & Minter** | UBC-F33032 | - | VUL.316 |
| Microthyriales | ***Microthyrium illicinum* De Not.** | CBS 143808 * | MG844144 | MG844151 |
| Microthyriales | ***Microthyrium macrosporum* (Sacc.) Höhn.** | CBS 143810 * | MG844147 | MG844159 |
| Microthyriales | *Microthyrium microscopicum* Desm. | CBS 115976 | GU296175.1 | GU301846.1 |
| Microthyriales | *Stomiopeltis betulae* J.P. Ellis | CBS 114420 | GU214701.1 | GU214701.1 |
| Microthyriales | *Tumidispora shoreae* Hongsanan & K.D. Hyde | MFLUCC 12-0409 | - | KT314074.1 |
| Microthyriales | *Heliocephala zimbabweensis* Decock, V. Robert & Masuka | MUCL 40019 | HQ333481.1 | HQ333481.1 |
| Monoblastiales Lücking, M.P. Nelsen & K.D. Hyde | *Acrocordia subglobosa* (Vězda) Poelt & Vězda | HTL940 | JN887373.1 | JN887392.1 |
| Monoblastiales | *Anisomeridium phaeospermum* R.C. Harris 1995 | MPN539 | JN887374.1 | JN887394.1 |
| Monoblastiales | *Funbolia dimorpha* Crous & Seifert | CPC 14170 | - | JF951156.1 |
| Monoblastiales | *Heleiosa barbatula* Kohlm., Volkm.-Kohlm. & O.E. Erikss. | JK 5548I | GU479753.1 | GU479787.1 |
| Monoblastiales | *Megalotremis verrucosa* (Makhija & Patw.) Aptroot | MPN104 | JN887383.1 | GU327718.1 |
| Monoblastiales | *Musaespora kalbii* Lücking & Sérus. | MPN243 | JN887391.1 | JN887406.1 |
| Monoblastiales | *Anisomeridium ubianum* (Vain.) R.C. Harris | MPN94 | JN887379.1 | GU327709.1 |
| Muyocopronales Mapook, Boonmee & K.D. Hyde | *Arxiella dolichandrae* Crous | CBS 138853 | - | KP004477.1 |
| Muyocopronales | *Muyocopron castanopsis* Mapook, Boonmee & K.D. Hyde | MFLUCC 14-1108 | KU726968.1 | KU726965.1 |
| Muyocopronales | *Muyocopron dipterocarpi* | MFLUCC 14-110 | KU726969.1 | KU726966.1 |
| Muyocopronales | *Muyocopron garethjonesii* Tibpromma, Karun. & K.D. Hyde | MFLU 16-2664 | KY070275.1 | KY070274.1 |
| Muyocopronales | *Muyocopron lithocarpi* Mapook, Boonmee & K.D. Hyde | MFLUCC 14-1106 | KU726970.1 | KU726967.1 |
| Muyocopronales | *Mycoleptodiscus indicus* (V.P. Sahni) B. Sutton | UAMH 8520 | - | GU980695.1 |
| Muyocopronales | *Paramycoleptodiscus albiziae* Crous & M.J. Wingf. | CPC 27552 | - | KX228330.1 |
| Muyocopronales | *Muyocopron* Speg. *sp.* | MFLU (CC) 10-0041 | JQ036226.1 | JQ036230.1 |
| Myriangiales | *Elsinoe veneta* (Burkh.) Jenkins | AFTOL-ID 1853 | DQ767651.1 | DQ767658.1 |
| Myriangiales Starbäck | *Endosporium aviarium* Tsuneda | UAMH 10530 | EU304349.1 | EU304351.1 |
| Myriangiales | *Endosporium populi-tremuloides* Tsuneda | UAMH 10529 | EU304346.1 | EU304348.1 |
| Myriangiales | *Myriangium hispanicum* J.B. Martínez | CBS 247.33 | GU296180.1 | GU301854.1 |
| Myriangiales | *Myriangium duriaei* Mont. & Berk. | CBS 260.36 | JGI | JGI |
| Mytilinidiales E. Boehm, C.L. Schoch & Spatafora | *Cenococcum geophilum* Fr. | JGI 1.58 v2.0 | JGI | JGI |
| Mytilinidiales | *Lepidopterella palustris* Shearer & J.L. Crane | CBS 459.81 | JGI | JGI |
| Mytilinidiales | *Lophium elegans* H. Zogg | EB 0366 | GU323184.1 | GU323210.1 |
| Mytilinidiales | *Lophium mytilinum* (Pers.) Fr. | AFTOL-ID 1609 | DQ678030.1 | DQ678081.1 |
| Mytilinidiales | *Mytilinidion mytilinellum* (Fr.) H. Zogg | CBS 303.34 | FJ161144.2 | FJ161184.2 |
| Mytilinidiales | *Mytilinidion scolecosporum* M.L. Lohman | CBS 305.34 | FJ161146.2 | FJ161186.2 |
| Natipusillales Raja, Shearer, A.N. Mill. & K.D. Hyde | *Natipusilla bellaspora* Raja, Shearer & A.N. Mill. | PE91 1a | JX474868.1 | JX474863.1 |
| Natipusillales | *Natipusilla decorospora* A. Ferrer, A.N. Mill. & Shearer | AF236 1A | HM196376.1 | HM196369.1 |
| Natipusillales | *Natipusilla naponensis* A. Ferrer, A.N. Mill. & Shearer | AF217 1A | HM196378.1 | HM196371.1 |
| Natipusillales | *Natipusilla limonensis* A. Ferrer, A.N. Mill. & Shearer | AF286 1A | HM196377.1 | HM196370.1 |
| Onygenales Cif. ex Benny & Kimbr. | *Spiromastix warcupii* Kuehn & G.F. Orr | AFTOL-ID 430 | DQ782882.1 | DQ782909.1 |
| Ostropales Nannf. | *Absconditella sphagnorum* Vězda & Poelt | T. Laukka 52 (TUR) | EU940022.1 | EU940095.1 |
| Ostropales | *Acarosporina microspora* (R.W. Davidson & R.C. Lorenz) Sherwood | AFTOL-ID 78 | AY584667.1 | AY584643.1 |
| Ostropales | *Cryptodiscus gloeocapsa* (Nitschke ex Arnold) Baloch, Gilenstam & Wedin | TSB 30770 | AF465456.1 | AF465440.1 |
| Ostropales | *Diploschistes cinereocaesius* (Sw.) Vain. | DUKE 0047509 | DQ883790.1 | DQ883790.1 |
| Ostropales | *Diploschistes ocellatus* (Fr.) Norman | Spain 995 | NG_013126.1 | NG_027624.1 |
| Ostropales | *Diploschistes rampoddensis* (Nyl.) Zahlbr. | - | AF274111.1 | AF274094.1 |
| Ostropales | *Diploschistes thunbergianus* (Ach.) Lumbsch & Vězda | Eldridge 3800 (F) | AF274112.1 | AF274095.1 |
| Ostropales | *Dyplolabia afzelii* (Ach.) A. Massal. | Luecking 26509a | - | HQ639628.1 |
| Ostropales | *Fissurina insidiosa* C. Knight & Mitt. | AFTOL-ID 1662 | DQ973022.1 | DQ973045.1 |
| Ostropales | *Fissurina marginata* Staiger | DNA3418 | - | JX421493.1 |
| Ostropales | *Gyalecta hypoleuca* (Ach.) Zahlbr. | TSB 20801 | AF465460.1 | AF465453.1 |
| Ostropales | *Gyalecta ulmi* (Sw.) Zahlbr. | Scheidegger 30.05.1998 (Duke) | AF465464.1 | AF465463.1 |
| Ostropales | ***Micropeltis* [Mont.] sp.** | UBC-F33034 | - | MG844154 |
| Ostropales | *Micropeltis zingiberaceicola* Henn. | IFRDCC 2264 | JQ036222.1 | JQ036227.1 |
| Ostropales | *Odontotrema phacidioides* Nyl. | Palice 11440 | - | HM244770.1 |
| Ostropales | *Porina farinosa* C. Knight | Lucking Pan-02 (F) | - | KJ449332.1 |
| Ostropales | *Porina guentheri* (Flot.) Zahlbr. | Lutzoni 97.10.09-10(Duke) | AF279404.1 | AF279405.1 |
| Ostropales | ***Scolecopeltidium* F. [Stevens & Manter] sp. 2** | UBC-F33035 | - | MG844155 |
| Ostropales | ***Scolecopeltidium* sp*.* 1** | UBC-F33033 | - | MG844153 |
| Ostropales | *Sphaeropezia arctoalpina* (Döbbeler & Poelt) Baloch, Gilenstam & Wedin | Baloch SW057 (S) | - | HM244760.1 |
| Ostropales | *Stictis radiata* (L.) Pers. | - | U20610.1 | **AF356663.1** |
| Ostropales | *Trapelia placodioides* Coppins & P. James | - | AF119500.2 | AF274103 |
| Ostropales | *Diploschistes muscorum* (Scop.) R. Sant. | Palice 2805 (HB Palice) | - | AY300836.1 |
| Patellariales D. Hawksw. & O.E. Erikss. | *Glyphium elatum* (Grev.) H. Zogg | EB 0342 | KM220935.1 | KM220938.1 |
| Patellariales | *Hysteropatella clavispora* (Peck) Höhn. | CBS 247.34 | DQ678006.1 | AY541493.1 |
| Patellariales | *Patellaria atrata* (Hedw.) Fr. | CBS 101060 | JGI | JGI |
| Patellariales | *Hysteropatella elliptica* (Fr.) Rehm | CBS 935.97, AFTOL-ID 1790 | EF495114.1 | DQ767657.1 |
| Pertusariales M. Choisy ex D. Hawksw. & O.E. Erikss. | *Dibaeis baeomyces* (L. f.) Rambold & Hertel | (SSU) taxon 83478, (LSU) **Lutzoni 93.08.20** | AF113712.1 | KJ462342.1 |
| Pertusariales | *Pertusaria dactylina* (Ach.) Nyl. | AFTOL-ID 224 | DQ782880.1 | DQ782907.1 |
| Phaeotrichales Ariyaw., Jian K. Liu & K.D. Hyde | *Trichodelitschia bisporula* (P. Crouan & H. Crouan) Munk ex N. Lundq. | CBS 262.69 | JGI | JGI |
| Phaeotrichales | *Phaeotrichum benjaminii* Malloch & Cain | CBS 541.72 | AY016348.1 | AY004340.1 |
| Pleosporales Luttr. ex M.E. Barr | *Aigialus grandis* Kohlm. & S. Schatz | JK 5244A | NG_016503.1 | GU301793.1 |
| Pleosporales | *Alternaria alternata* (Fr.) Keissl. | SRC1lrK2f | HM216191.1 | HM216200.1 |
| Pleosporales | *Arthopyrenia salicis* A. Massal. | CBS 368.94 | AY538333.1 | AY538339.1 |
| Pleosporales | *Ascocratera manglicola* | JK 5262C | GU296136.1 | GU301799.1 |
| Pleosporales | *Astrosphaeriella stellata* (Pat.) Sacc. | MFLUCC10-0095 | JN846741.1 | JN846720.1 |
| Pleosporales | *Astrosphaeriella vesuvius* (Berk. & Broome) D. Hawksw. & Boise | LF-157 | KP814137.1 | KP814136.1 |
| Pleosporales | *Bipolaris maydis* (Y. Nisik. & C. Miyake) Shoemaker | AFTOL-ID 54 | AY544727.1 | AY544645.1 |
| Pleosporales | *Chaetasbolisia erysiphoides* (Griffon & Maubl.) Griffon & Maubl. | CBS 148.94 | EU754041.1 | EU754140.1 |
| Pleosporales | *Cheirosporium triseriale* L. Cai & K.D. Hyde | HMAS 180703 | - | EU413954.1 |
| Pleosporales | *Dictyocheirospora rotunda* M.J. D'souza, Bhat & K.D. Hyde | DLUCC 0577 | - | KY320517.1 |
| Pleosporales | *Dictyosporium elegans* Corda | NBRC 32502 | DQ018079.1 | DQ018100.1 |
| Pleosporales | *Halojulella avicenniae* | JK 5326A | GU479756.1 | GU479790.1 |
| Pleosporales | *Halojulella avicenniae* (Borse) Suetrong, K.D. Hyde & E.B.G. Jones | BCC 18422 | GU371831.1 | GU371823.1 |
| Pleosporales | *Jalapriya pulchra* M.J. D'souza, Hong Y. Su, Z.L. Luo & K.D. Hyde | SB-2016c isolate LQXM47 | KU179110.1 | KU179109.1 |
| Pleosporales | *Leptosphaeria heterospora* (De Not.) Niessl | CBS 644.86 | AY016354.1 | AY016369.1 |
| Pleosporales | *Pleospora herbarum* (Pers.) Rabenh. | CBS 191.86 | DQ247812.1 | JX681120.1 |
| Pleosporales | *Pseudocoleophoma polygonicola* Kaz. Tanaka & K. Hiray. | KT 731 | AB797256.1 | AB807546.1 |
| Pleosporales | *Pseudodictyosporium wauense* Matsush. | NBRC 30078 | DQ018083.1 | DQ018105.1 |
| Pleosporales | *Pyrenophora phaeocomes* (Rebent.) Fr. | AFTOL-ID 283 | DQ499595.1 | NG_027575.1 |
| Pyrenulales Fink ex D. Hawksw. & O.E. Erikss. | *Pyrenula pseudobufonia* (Rehm) R.C. Harris | Reeb VR (DUKE) | AY641001.1 | AY640962.1 |
| Pyrenulales | *Pyrgillus javanicus* (Mont. & Bosch) Nyl. | AFTOL-ID 342 | DQ823110.1 | DQ823103.1 |
| Rhytismatales M.E. Barr ex Minter | *Coccomyces dentatus* (J.C. Schmidt & Kunze) Sacc. | AFTOL-ID 147 | AY544701.1 | AY544657.1 |
| Strigulales Lücking, M.P. Nelsen & K.D. Hyde | *Flavobathelium epiphyllum* Lücking, Aptroot & G. Thor | MPN67 | JN887382.1 | GU327717.1 |
| Strigulales | *Phyllobathelium anomalum* Lücking | MPN242 | JN887386.1 | GU327722.1 |
| Strigulales | *Strigula jamesii* (Swinscow) R.C. Harris | MPN548 | JN887388.1 | JN887404.1 |
| Strigulales | *Strigula nemathora* Mont. | MPN72 | JN887389.1 | JN887405.1 |
| Strigulales | *Strigula schizospora* R. Sant. | MPN73 | JN887390.1 | - |
| Strigulales | *Taeniolella exilis* (P. Karst.) S. Hughes | CBS122902 | - | KX244968.1 |
| Strigulales | *Phyllobathelium firmum* (Stirt.) Vězda | MPN545 | JN887387.1 | JN887401.1 |
| Trypetheliales Lücking, Aptroot & Sipman | *Astrothelium megaspermum* (Mont.) Aptroot & Lücking | AFTOL-ID 2094 | GU561841.1 | FJ267702.1 |
| Trypetheliales | *Mycomicrothelia hemisphaerica* (Müll. Arg.) D. Hawksw. | MPN102 | JN887384.1 | GU327719.1 |
| Trypetheliales | *Mycomicrothelia miculiformis* (Nyl. ex Müll. Arg.) D. Hawksw. | MPN101B | JN887385.1 | GU327720.1 |
| Trypetheliales | *Trypethelium eluteriae* Spreng. | CBS 132375 | JGI | JGI |
| Trypetheliales | *Trypethelium nitidiusculum* (Nyl.) R.C. Harris | AFTOL-ID 2099 | GU561842.1 | GU561856.1 |
| Tubeufiales Boonmee & K.D. Hyde | *Helicomyces roseus* Link | AFTOL-ID 1613 | DQ678032.1 | DQ678083.1 |
| Tubeufiales | *Tubeufia cerea* (Berk. & M.A. Curtis) Höhn. | AFTOL-ID 1316 | DQ471034.1 | DQ470982.1 |
| Tubeufiales | *Tubeufia helicomyces* Höhn. | AFTOL-ID 1580 | DQ767649.1 | DQ767654.1 |
| Tubeufiales | *Wiesneriomyces conjunctosporus* Kuthub. & Nawawi | BCC18525 | KJ425436.1 | KJ425450.1 |
| Tubeufiales | *Tubeufia paludosa* (P. Crouan & H. Crouan) Rossman | CBS 120503 | GU296203.1 | GU301877.1 |
| Umbilicariales J.C. Wei & Q.M. Zhou | *Umbilicaria mammulata* (Ach.) Tuck. | (SSU)Wei 96042, 1996, (LSU)**AFTOL-ID 645** | AY648114.1 | DQ782912.1 |
| Venturiales Y. Zhang ter, C.L. Schoch & K.D. Hyde | *Apiosporina collinsii* (Schwein.) Höhn. | CBS 118973 | GU296135.1 | GU301798.1 |
| Venturiales | *Aulographina pinorum* (Desm.) Arx & E. Müll. | CBS 655.86 | - | KF902102.1 |
| Venturiales | *Dibotryon morbosum* (Schwein.) Theiss. & Syd. | Oregon-non-cultivated ‘dimosp’ | EF114718.1 | EF114694.1 |
| Venturiales | *Fusicladium oleagineum* (Castagne) Ritschel & U. Braun | CBS 113427 | KF766251.1 | KF766331.1 |
| Venturiales | *Gibbera conferta* (Fr.) Petr. | CBS 191.53 | GU296150.1 | GU301814.1 |
| Venturiales | *Metacoleroa dickiei* (Berk. & Broome) Petr. | Oregon-non-cultivated | EF114719.1 | EF114695.1 |
| Venturiales | *Protoventuria barriae* Carris & A.P. Poole | ATCC 90285 | EF114728.1 | JQ036232.1 |
| Venturiales | ***Stomiopeltis* [Theiss.] sp.** | CBS 143811 * | MG844148 | MG844160 |
| Venturiales | ***Stomiopeltis* sp.** | UBC-F33041 | MG844149 | MG844161 |
| Venturiales | *Tothia fuscella* (Sacc.) Bat. | CBS 130266 | JGI | JGI |
| Venturiales | *Tyrannosorus pinicola* (Petrini & P.J. Fisher) Unter. & Malloch | AFTOL-ID 1235 | DQ471025.1 | DQ470974.1 |
| Venturiales | *Venturia inaequalis* (Cooke) G. Winter | CBS 594.70 | KF156093.1 | GU301879.1 |
| Venturiales | *Venturia pyrina* Aderh. | ICMP 11032 | JGI | JGI |
| Venturiales | *Veronaeopsis simplex* (Papendorf) Arzanlou & Crous | CBS 588.66 | KF156095.1 | EU041877.1 |
| Venturiales | *Sympoventuria capensis* Crous & Seifert | CBS 120136 | KF156094.1 | KF156104.1 |
| Verrucariales Mattick ex D. Hawksw. & O.E. Erikss. | *Endocarpon pallidulum* (Nyl.) Nyl. | AFTOL-ID 661 | DQ823097.1 | DQ823104.1 |
| Verrucariales | *Staurothele frustulenta* Vain. | AFTOL-ID 697 | DQ823105.1 | DQ823098.1 |
| Zeloasperisporiales Hongsanan & K.D. Hyde | *Zeloasperisporium cliviae* Crous | CPC 25145 | - | KR476781.1 |
| Zeloasperisporiales | *Zeloasperisporium eucalyptorum* Cheew. & Crous | CBS 124809 | - | GQ303329.1 |
| Zeloasperisporiales | *Zeloasperisporium ficusicola* Hongsanan & K.D. Hyde | MFLUCC 15-0222 | KT387736.1 | KT387735.1 |
| Zeloasperisporiales | *Zeloasperisporium searsiae* Crous & A.R. Wood | CPC 25880 | - | KT950866.1 |
| Zeloasperisporiales | *Zeloasperisporium siamense* (Boonmee, H.X. Wu & K.D. Hyde) Hongsanan & K.D. Hyde | IFRDCC 2194 | JQ036223.1 | JQ036228.1 |
| Zeloasperisporiales | *Zeloasperisporium wrightiae* Hongsanan & K.D. Hyde | MFLUCC 15-0215 | KT387746.1 | KT387742.1 |
| Incertae sedis | *Arthrographis kalrae* (R.P. Tewari & Macph.) Sigler & J.W. Carmich. | IFM 52423 (YRL) | - | AB116544.1 |
| Incertae sedis | *Caryosporella rhizophorae* Kohlm. | JK 5386C | GU479750.1 | GU479784.1 |
| Incertae sedis | *Collophora africana* Damm & Crous | CBS 120872 | GQ154630.1 | GQ154609.1 |
| Incertae sedis | *Coniosporium apollinis* Sterfl. | CBS 352.97 | GU250916.1 | GU250895.1 |
| Incertae sedis | *Coniosporium uncinatum* De Leo, Urzì & de Hoog | CBS 100212 | GU250922.1 | GU250902.1 |
| Incertae sedis | *Cryomyces antarcticus* Selbmann, de Hoog, Mazzaglia, Friedmann & Onofri | CCFEE 536 | GU250321.1 | GU250365.1 |
| Incertae sedis | *Cryomyces minteri* Selbmann, de Hoog, Mazzaglia, Friedmann & Onofri | CCFEE 5187 | KC315858.1 | GU250369.1 |
| Incertae sedis | *Dothideomycetes* [O.E. Erikss. & Winka] sp. | CCFEE5460 | GU250349.1 | GU250391.1 |
| Incertae sedis | *Dothideomycetes* sp. | CCFEE5416 | GU250344.1 | GU250387.1 |
| Incertae sedis | *Encephalographa elisae* A. Massal. | EB 0347 | GU397358.1 | GU397343.1 |
| Incertae sedis | *Eremomyces bilateralis* Malloch & Cain | CBS 781.70 | JGI | JGI |
| Incertae sedis | *Farlowiella carmichaeliana* (Berk.) Sacc. | CBS 164.76 | GU296129.1 | GU301791.1 |
| Incertae sedis | *Hysterographium fraxini* (Pers.) De Not. | CBS 109.43 | FJ161132.2 | FJ161171.2 |
| Incertae sedis | *Lichenothelia calcarea* Henssen | L1324 | KC015082.1 | KC015062.1 |
| Incertae sedis | *Lichenothelia convexa* Henssen | L1609 | KC015086.1 | KC015071.1 |
| Incertae sedis | *Lineolata rhizophorae* (Kohlm. & E. Kohlm.) Kohlm. & Volkm.-Kohlm. | CBS 641.66 | GU479758.1 | GU479792.1 |
| Incertae sedis | *Minutisphaera fimbriatispora* Shearer, A.N. Mill. & A. Ferrer | G155.1 | JX474865.1 | JX474859.1 |
| Incertae sedis | *Minutisphaera japonica* Kaz. Tanaka, Raja & Shearer | KTC 2738 | AB733434.1 | NG_042338.1 |
| Incertae sedis | *Pseudeurotium hygrophilum* (Sogonov, W. Gams, Summerb. & Schroers) Minnis & D.L. Lindner | isolate 229 | JQ780655.1 | JQ780654.1 |
| Incertae sedis | *Rhexothecium globosum* Samson & Mouch. | CBS 955.73 | - | HG004544.1 |
| Incertae sedis | *Saxomyces alpinus* Zucconi & Selbmann | CCFEE 5466 | GU250350.1 | GU250392.1 |
| Incertae sedis | *Saxomyces penninicus* Zucconi & Onofri | CCFEE 5495 | KC315875.1 | KC315864.1 |
| Incertae sedis | *Geastrumia polystigmatis* Bat. & M.L. Farr | strain NC4 1.8F1a | - | FJ147177.1 |

Notes: Isolates included in this study; names in bold indicate newly contributed collections and voucher/strain numbers beginning with 'CBS' and followed by '*' represent cultures newly deposited in the Westerdijk Institute. When LSU and SSU were derived from two different collections, bold indicates the voucher/strain name that is used in phylogenies. Accessions labeled 'JGI' refer to sequences extracted from JGI genomes project, for which no GenBank accession exists.

### Appendix S2 JGI sequence data retrieved from MycoCosm portal

| **Taxon/strain** | **Blast target** | **SSU Sequence length (bp)** | **SSU assembly** | **LSU Sequence length (bp)** | **LSU Assembly** | **Publication** | **Genome project PI** |
| --- | --- | --- | --- | --- | --- | --- | --- |
| *Acidomyces richmondensis* C2 | Assembly | 1729 | 8 scaffolds, 2 bp conflicts | 2995 | 6 scaffolds, 1 bp conflicts | Mosier et al. (2016)^1^ | Singer, Steven - Berkeley Lab, USA |
| *Aspergillus niger* ATCC 1015 | Assembly | 1738 | Single scaffold | 3424 | Single scaffold | Andersen et al. (2011)^2^ | Baker, Scott - Environmental Molecular Sciences Laboratory, USA |
| *Aulographum hederae* CBS 113979 | Assembly | 1791 | Single scaffold with SSU and LSU | 2934 | Single scaffold with SSU and LSU | permission granted | Spatafora, Joseph - Oregon State University, USA |
| *Aureobasidium subglacialis* EXF-2481 | EST | 1731 | 5 clusters | 3092 | 3 clusters | Gostincar et al. (2014)^3^ | Gunde-Cimerman, Nina - University of Ljubljana, Department of Biology, Slovenia |
| *Cenococcum geophilum* strain 1.58 | Assembly + EST | 1737 | Single scaffold for both regions | 3610 | 1 scaffold + 3 clusters | Peter et al. (2016)^4^ | Martin, Francis - INRA Nancy, France |
| *Eremomyces bilateralis* CBS 781.70 |  | 1729 | Single cluster | 2875 | Single cluster | permission granted | Crous, Pedro and Binder, Manfred - CBS-KNAW, Netherlands |
| *Lepidopterella palustris* CBS 459.81 | EST | 1728 | 2 clusters | 2951 | 2 clusters | Peter et al. (2016) ^4^ | Spatafora, Joseph |
| *Myriangium duriaei* CBS 260.36 | Genbank + Assembly | 1703 | genbank sequence | 2969 | 1 scaffold | permission granted | Spatafora, Joseph |
| *Patellaria atrata* CBS 101060 | Assembly + EST | 1477 | single scaffold from assembly | 3011 | Contig of 4 ESTs | permission granted | Spatafora, Joseph |
| *Tothia fuscella* CBS 130266 | Assembly | 1738 | Single scaffold, big insert | 1125 | Single scaffold | permission granted | Binder, Manfred |
| *Trichodelitschia bisporula* CBS 262.69 |  | 1731 | 5 clusters, 1bp conflict | 3015 | 3 clusters | permission granted | Binder, Manfred |
| *Trypethelium eluteriae* MPN111 | EST | 1122 | Single cluster | 3292 | 2 clusters | McDonald et al. (2013)^5^ | Binder, Manfred |
| *Venturia pirina* ICMP 11032 | EST | 1725 | Single scaffold | 3158 | Single scaffold | Cooke et al. (2014)^6^ | Deng, Cecilia - The New Zealand Institute for Plant & Food Research Limited, New Zealand |

### Appendix S3. Results from CONSEL for six topologies tested against most likely tree

| **Test Statistic** | **Approximately Unbiased test** | **Kishino Hasegawa Test** | **Bootstrap probability** | **Constraint** |
| --- | --- | --- | --- | --- |
| 55.9 | 0.086 | 0.06 | 0.029 | Asterinales and Asterotexiales monophyletic |
| 102.3 | 0.011 | 0.006 | 0.002 | Microthyriales + Zeloasperisporiales monophyletic |
| 161 | 3.00E-04 | 0 | 2.00E-05 | *Lembosia* monophyletic |
| 1242.2 | 4.00E-06 | 0 | 1.00E-05 | Radiate thyriothecia monophyletic |
| 419.2 | 2.00E-80 | 0 | 4.00E-21 | *Asterina* spp. monophyletic |

### Appendix S4. Matrix of characters used in ancestral character state reconstructions and in parsimony analyses of fossils.

| **Order** | **Taxon** | **Sub** | **Lic** | **Spo** | **Low** | **Deh** | **Loc** | **Rad** | **Bra** | **Mar** | **App** | **Ini** | **Ref** | **Illustration in reference** |
| --- | --- | --- | --- | --- | --- | --- | --- | --- | --- | --- | --- | --- | --- | --- |
| Fossil, Early Triassic of India (Induan, ~251 Ma) | "Fungal thallus" | ? | ? | 2 | ? | 0 | 0 | 1 | 1? 2? | ? | ? | ? | 1 | Fig. 8j |
| Fossil, early Eocene of India (Ypresian, 56–47.8 Ma) | Trichothyrites setifer | ? | ? | 2 | 1 | 0 | 0 | 1 | 2 | 2 | ? | ? | 2 | Pl.2 Fig, 18 |
| Fossil, early Eocene of USA (Ypresian, 56–47.8 Ma) | Asterina eocenica | 1 | 0 | 2 | 0 | 1 | 0 | 1 | 1 | 1 | 1 | 1 | 3 | Pl. 7, Fig. 56; Pl. 8, Figs. 57—68 |
| Abrothallales | *Abrothallus acetabuli* SPO308 | 2 | 0 | 0 | 1 | 4 | 0 | 0 | 0 | 0 | 0 | ? | 7 | all *Abrothallus* in ^7^ coded similarly |
| Abrothallales | *Abrothallus buellianus* SPO303 | 2 | 0 | 0 | 1 | 4 | 0 | 0 | 0 | 0 | 0 | ? | 7 | " |
| Abrothallales | *Abrothallus cladoniae* AB53 | 2 | 0 | 0 | 1 | 4 | 0 | 0 | 0 | 0 | 0 | ? | 7 | " |
| Abrothallales | *Abrothallus hypotrachynae* SPO302 | 2 | 0 | 0 | 1 | 4 | 0 | 0 | 0 | 0 | 0 | ? | 7 | " |
| Abrothallales | *Abrothallus parmeliarum* AB36 | 2 | 0 | 0 | 1 | 4 | 0 | 0 | 0 | 0 | 0 | ? | 7 | " |
| Abrothallales | *Abrothallus parmotrematis* AB1 | 2 | 0 | 0 | 1 | 4 | 0 | 0 | 0 | 0 | 0 | ? | 8 | " |
| Abrothallales | *Abrothallus secedens* SPO305 | 2 | 0 | 0 | 1 | 4 | 0 | 0 | 0 | 0 | 0 | ? | 7 | " |
| Abrothallales | *Abrothallus suecicus* AB56 | 2 | 0 | 0 | 1 | 4 | 0 | 0 | 0 | 0 | 0 | ? | 7 | " |
| Abrothallales | *Abrothallus usneae* AB20 | 2 | 0 | 0 | 1 | 4 | 0 | 0 | 0 | 0 | 0 | ? | 7 | " |
| Ostropales | *Absconditella sphagnorum* EU940095.1 | 4 | 0 | 0 | 1 | 4 | 0 | 0 | 0 | 0 | 0 | ? | 9 | Fig. 3b |
| Ostropales | *Acarosporina microspora* AFTOL-ID 78 | 3 | 0 | 0 | 1 | 4 | 0 | 0 | 0 | 0 | 0 | ? | 10 | p. 36 |
| Capnodiales | *Acidomyces richmondensis* JGI | ? | 0 | ? | ? | ? | ? | 0 | ? | ? | 0 | ? | 11 | Fig. 8 |
| Monoblastiales | *Acrocordia subglobosa* HTL940 | 0 [^*^](http://lichenportal.org/imglib/lichens/misc/201605/index_1463618021_web.jpg) | 1 | 1 | 1 | 0 | 0 | 0 | 0 | 0 | 0 | ? |  | Based on *A. cavata* morphology UBC L47305 |
| Acrospermales | *Acrospermum adeanum* M133 | 1 | 0 | 1 | 1 | 0 | 0 | 0 | 0 | 0 | 0 | ? | 9 | Fig. 3a |
| Acrospermales | *Acrospermum compressum* M151 | 1 | 0 | 1 | 1 | 0 | 0 | 0 | 0 | 0 | 0 | ? | 12 | Fig. 6–12 |
| Acrospermales | *Acrospermum graminum* M152 | 1 | 0 | 1 | 1 | 0 | 0 | 0 | 0 | 0 | 0 | ? |  | [link](http://www.ascofrance.com/recolte/2470/dothideomycetes-incertae-sedis-acrospermaceae-acrospermum-graminum) |
| Pleosporales | *Aigialus grandis* JK 5244A | 3 | 0 | 1 | 1 | 0 | 0 | 0 | 0 | 0 | 0 | ? | 13 | Fig. 1–3 |
| Pleosporales | *Alternaria alternata* SRC1lrK2f | 1 | 0 | 1 | 1 | ? | 0 | 0 | ? | 0 | 0 | ? | 14 | Based on genus description |
| Arthoniales | *Alyxoria varia* EU704103.1 | 3 | 1 | 0 | 1 | 2 | 1 | 0 | 0 | 0 | 0 | ? | 15 16 | Fig. 551 |
| Monoblastiales | *Anisomeridium phaeospermum* MPN539 | 3 | 1 | 1 | 1 | 0 | 0 | 0 | 0 | 0 | 0 | ? | 17 | Figs. 2c; 3b |
| Monoblastiales | *Anisomeridium ubianum* MPN94 | 3 | 1 | 1 | 1 | 0 | 0 | 0 | 0 | 0 | 0 | ? | 17 | derived from *A. phaeospermum* |
| Venturiales | *Apiosporina collinsii* CBS 118973 | 1 | 0 | 1 | 1 | 3 | 0 | 0 | 0 | 0 | 0 | ? | 18 | Pl. XVI |
| Arthoniales | *Arthonia dispersa* UPSC 2583 | 3 | 1 | 0 | 0 | 2 | 1 | 0 | 0 | 0 | 0 | ? | 15 19 | from genus description |
| Pleosporales | *Arthopyrenia salicis* CBS 368.94 | 3 | 0 | 1 | 1 | 0 | 0 | 0 | 0 | 0 | 0 | ? |  | [link](http://www.irishlichens.ie/pages-lichen/l-458.html) |
| Capnodiales | *Arthrocatena tenebrio* CCFEE 5413 | 0 | 0 | ? | ? | ? | ? | 0 | ? | ? | 0 | ? | 20 | Figs. 15d­–15g |
| Incertae sedis | *Arthrographis kalrae* IFM 52423(YRL) | ? | 0 | ? | ? | ? | ? | 0 | ? | ? | 0 | ? | 21 | Fig. 3 |
| Muyocopronales | *Arxiella dolichandrae* CBS 138853 | 1 | 0 | ? | ? | ? | ? | 0 | ? | ? | 0 | ? | 22 | p. 226 |
| Pleosporales | *Ascocratera manglicola* JK 5262C | 3 | 0 | 1 | 1 | 0 | 0 | 0 | 0 | 0 | 0 | ? | 23 | Figs. 1–7 |
| Eurotiales | *Aspergillus fumigatus* JCM1738 | ? | 0 | 3 | 1 | 3 | 0 | 0 | 0 | 0 | 0 | ? |  | [link](https://mycology.adelaide.edu.au/descriptions/hyphomycetes/aspergillus/) |
| Eurotiales | *Aspergillus niger* ATCC 1015 | ? | 0 | 3 | 1 | 3 | 0 | 0 | 0 | 0 | 0 | ? |  | [link](https://mycology.adelaide.edu.au/descriptions/hyphomycetes/aspergillus/) |
| Asterotexiales | *Asterina cestricola* TH 591 | 1 | 0 | 2 | 0 | 1 | 0 | 1 | 1 | 1 | 1 | 1 | 24 | Fig. 3.1 |
| Asterinales | *Asterina chrysophylli* VIC 42823 | 1 | 0 | 2 | 0 | 1 | 0 | 1 | 1 | 1 | 1 | 1 | 25 | Fig. 4 |
| Asterotexiales | *Asterina cynometrae* MFLU 13-0373 | 1 | 0 | 2 | 0 | 1 | 0 | 1 | 1 | 1 | 1 | 1 | 26 | Fig. 2 |
| Asterotexiales | *Asterina fuchsiae* TH 590 | 1 | 0 | 2 | 0 | 1 | 0 | 1 | 1 | 1 | 1 | 1 | 27 | Fig. 6 |
| Asterinales | *Asterina melastomatis* VIC 42822 | 1 | 0 | 2 | 0 | 1 | 0 | 1 | 1 | 1 | 1 | 1 | 25 | Fig. 3 |
| Asterotexiales | *Asterina phenacis* TH 589 | 1 | 0 | 2 | 0 | 1 | 0 | 1 | 1 | 1 | 1 | 1 | 27 | Fig. 8 |
| Asterotexiales | *Asterina siphocampyli* M 0141060 PMA | 1 | 0 | 2 | 0 | 1 | 0 | 1 | 1 | 1 | 1 | 1 | 28 | Fig. 7 |
| Asterotexiales | *Asterina* sp. MFLU13-0619 | 1 | 0 | 2 | 0 | 1 | 0 | 1 | 1 | 1 | 1 | 1 | 29 | Fig. 5 |
| Asterotexiales | *Asterina weinmanniae* TH592 | 1 | 0 | 2 | 0 | 1 | 0 | 1 | 1 | 1 | 1 | 1 | 24 | Fig. 3 |
| Asterotexiales | *Asterina zanthoxyli* TH 561 | 1 | 0 | 2 | 0 | 1 | 0 | 1 | 1 | 1 | 1 | 1 | 24 | Fig. 4 |
| Asterotexiales | Asterotexiaceae sp.1 UBC-F33036 | 1 | 0 | 2 | 0 | 0 | 0 | 1 | 1 | 1 | 0 | 1 |  | Personal collection |
| Asterotexiales | Asterotexiaceae sp.2 CBS 143813 | 1 | 0 | 2 | 0 | 1 | 0 | 1 | ? | 1 | 0 | 0 |  | Personal collection |
| Asterotexiales | *Asterotexis cucurbitacearum* PMA M-0141224 | 1 | 0 | 2 | 0 | 1 | 1 | 1 | ? | 1 | 0 | 2 | 30 | Figs. 1–2 |
| Asterotexiales | *Asterotexis cucurbitacearum* VIC 42814 | 1 | 0 | 2 | 0 | 1 | 1 | 1 | ? | 1 | 0 | 2 | 25 | Fig. 8 |
| Pleosporales | *Astrosphaeriella stellata* MFLUCC10-0095 | 1 | 0 | 1 | 1 | 0 | 0 | 0 | 0 | 0 | 0 | ? | 31 | Fig. 2 |
| Pleosporales | *Astrosphaeriella vesuvius* LF-157 | 3 | 0 | 1 | 1 | 0 | 0 | 0 | 0 | 0 | 0 | ? | 31 | Derived from *A. stellata* |
| Trypetheliales | *Astrothelium megaspermum* AFTOL-ID 2094 | 3 | 1 | 1 | 1 | 0 | 0 | 0 | 0 | 0 | 0 | ? | 32 | Fig.1O |
| Trypetheliales | *Astrothelium nitidiusculum* AFTOL-ID 2099 | 3 | 1 | 1 | 1 | 0 | 0 | 0 | 0 | 0 | 0 | ? | 33 | Figs.1; 3A; 4D; 4B; 6A |
| Capnodiales | *Aulographina pinorum* CBS 174.90 | 1 | 0 | ? | ? | ? | ? | ? | ? | ? | 0 | ? | 34 | *Abb. 1* |
| Venturiales | *Aulographina pinorum* CBS 655.86 | ? | 0 | ? | ? | ? | ? | ? | ? | ? | 0 | ? | 34 | *Abb. 1* |
| Aulographaceae | *Aulographum hederae* CBS 113979 | 1 | 0 | 2 | 0 | 1 | 1 | 1 | 3 | 1 | 0 | 4 | 35 | Fig. 23 |
| Aulographaceae | *Aulographum hederae* MFLUCC13-0001 | 1 | 0 | 2 | 0 | 1 | 1 | 1 | 3 | 1 | 0 | 4 | 35 | Fig. 23 |
| Aulographaceae | *Aulographum* sp. CBS 143545 | 1 | 0 | 2 | 0 | 1 | 1 | 1 | 3 | 1 | 0 | 4 |  | Personal collection |
| Capnodiales | *Aureobasidium subglaciale* EXF-2481 | ? | 0 | ? | ? | ? | ? | 0 | ? | ? | 0 | ? |  | [link](https://mycology.adelaide.edu.au/descriptions/hyphomycetes/aureobasidium/) |
| Asterinales | *Batistinula gallesiae* B VIC 42514 | 1 | 0 | 2 | 0 | 1 | 0 | 1 | 1 | 1 | 1 | 1 | 25 | Fig. 5 |
| Pleosporales | *Bipolaris maydis* AFTOL-ID 54 | ? | 0 | 1 | 1 | ? | 0 | 0 | ? | 0 | 0 | ? | 36 | Fig. 22 |
| Asterinales | *Blastacervulus eucalypti* CBS 124759 | 1 | 0 | ? | ? | ? | ? | 0 | ? | ? | 0 | ? | 37 | Fig. 4 |
| Asterinales | *Blastacervulus eucalyptorum* CPC 29450 | 1 | 0 | ? | ? | ? | ? | 0 | ? | ? | 0 | ? | 38 | p. 292 |
| Asterinales | *Blasttacervulus robbenensis* CBS 124780 | 1 | 0 | ? | ? | ? | ? | 0 | ? | ? | 0 | ? | 39 | Fig. 3 |
| Botryosphaeriales | *Botryosphaeria dothidea* CBS 115476 | 3 | 0 | 1 | 1 | 0 | 0 | 0 | 0 | 0 | 0 | ? | 40 | Figs. 1–7 |
| Asterotexiales | *Buelliella minimula* Lendemer 42273(NY) | 2 | 0 | 0 | 1 | 4 | 0 | 0 | 0 | 0 | 0 | ? | 41 | Figs. 4a; 5a |
| Asterotexiales | *Buelliella physciicola* Ertz 19173(BR) | 2 | 0 | 0 | 1 | 4 | 0 | 0 | 0 | 0 | 0 | ? | 41 | Figs. 4b; 5c |
| Asterotexiales | *Buelliella poetschii* Ertz 18116(BR) | 2 | 0 | 0 | 1 | 4 | 0 | 0 | 0 | 0 | 0 | ? | 41 | Figs. 4c; 5d |
| Capnodiales | *Capnobotryella renispora* CBS 214.90 | 1 | 0 | 1 | ? | 0 | ? | 0 | ? | ? | 0 | ? | 42 43 | Fig. 7.5–7.8, and Pl. 170A in ^43^ |
| Capnodiales | *Capnodium citri* CBS 451.66 | 1 | 0 | 1 | 1 | 0 | ? | 0 | ? | 0 | 0 | ? | 44 | Fig. 9 |
| Capnodiales | *Capnodium coffeae* CBS 147.52 | 1 | 0 | 1 | 1 | 0 | 0 | 0 | 0 | 0 | 0 | ? | 44 | Following genus description |
| Chaetothyriales | *Capronia pilosella* AFTOL-ID 657 | 3 | 0 | 1 | 1 | 0 | 0 | 0 | 0 | 0 | 0 | ? | 45 | Figs. 14–19 |
| Incertae sedis | *Caryosporella rhizophorae* JK 5386C | 3 | 0 | 1 | 1 | **0** | 0 | 0 | 0 | 0 | 0 | ? | 46 | Figs. 1–5 |
| Capnodiales | *Catenulostroma chromoblastomycosum* CBS 597.97 | ? | 0 | ? | ? | ? | ? | 0 | ? | ? | 0 | ? | 47 | Fig. 6 |
| Mytilinidiales | *Cenococcum geophilum* JGI 1.58 v2.0 | 4 | 0 | ? | ? | ? | ? | 0 | ? | ? | 0 | ? | 48 | NA |
| Chaetothyriales | *Ceramothyrium carniolicum* CBS 175.95 | 1 | 0 | ? | ? | ? | 0 | ? | ? | ? | 0 | ? | 49 | Derived from *C. thailandicum*, Fig. 3 |
| Chaetothyriales | *Ceramothyrium linnaeae* UPSC 2646 | 1 | 0 | 1 | 1 | ? | 0 | 0 | 4 | 3 | 0 | ? | 50 | Fig. 7 |
| Chaetothyriales | *Ceramothyrium podocarpi* CPC 19826 | 1 | 0 | ? | ? | ? | ? | ? | ? | ? | 0 | ? | 49 | Derived from *C. thailandicum*, Fig. 3 |
| Capnodiales | *Cercospora zebrina* CBS 118790 | ? | 0 | 1 | 1 | 0 | 0 | 0 | ? | 0 | 0 | ? | 51 |  |
| Lichenostigmatales | Cf. Arthoniales sp. CCFEE 5176 | 0 | 0 | ? | ? | ? | ? | 0 | ? | ? | 0 | ? | 52 | none |
| Venturiales | cf. *Stomiopeltis* sp. 1 CBS 143811 | 1 | 0 | 2 | 0 | 0 | 0 | 1 | 4 | 1 | 0 | ? |  | Personal collection |
| Venturiales | cf. *Stomiopeltis* sp. 2 UBC-F33041 | 1 | 0 | 2 | 0 | 0 | 0 | 1 | 4 | 1 | 0 | ? |  | Personal collection |
| Capnodiales | *Chaetasbolisia erysiphoides* CBS 148.94 | 0 | 0 | ? | ? | 0 | ? | 0 | ? | ? | 0 | ? | 53 | Figs. 1–2 |
| Microthyriales | *Chaetothyriothecium elegans* CPC 21375 | 1 | 0 | 2 | ? | 0 | ? | 1 | ? | ? | 0 | ? | 54 | Fig. 2 |
| Chaetothyriales | *Chaetothyrium agathis* MFLUCC 12 C0113 | 1 | 0 | 1 | 1 | 3 | 0 | 0 | 0 | 0 | 0 | ? | 55 | Fig. 132 |
| Chaetothyriales | *Chaetothyrium brischoficola* MFLUCC 10-0012 | 1 | 0 | 1 | 1 | 3 | 0 | 0 | 0 | 0 | 0 | ? | 49 | Derived from *C. thailandicum*, Fig. 3 |
| Pleosporales | *Cheirosporium triseriale* HMAS 180703 | 3 | 0 | ? | ? | ? | ? | 0 | ? | ? | 0 | ? | 56 | Fig. 3 |
| Arthoniales | *Chrysothrix candelaris* KF707640.1 | 0 | 1 | 0 | 1 | 4 | 0 | 0 | 0 | 0 | 0 | ? |  | [link](http://lichenportal.org/portal/taxa/index.php?taxon=52529) |
| Lecanorales | *Cladonia caroliniana* AFTOL-ID 3 | 0 | 1 | 0 | 1 | 4 | 0 | 0 | 0 | 0 | 0 | ? | 15 | Fig. 220 |
| Capnodiales | *Cladosporium bruhnei* CPC 5101 | ? | 0 | ? | ? | ? | ? | 0 | ? | ? | 0 | ? | 57 | Fig. 9 |
| Rhytismatales | *Coccomyces dentatus* AFTOL-ID 147 | 1 | 0 | 0 | 1 | 4 | 0 | 0 | 0 | 0 | 0 | ? | 58 | Fig. 19 |
| Incertae sedis | *Collophora africana* CBS 120872 | 3 | 0 | ? | ? | ? | ? | 0 | ? | ? | 0 | ? | 59 | Fig. 7 |
| Arthoniales | *Combea mollusca* Tehler 7725 | 0 | 1 | 0 | 1 | 4 | 0 | 0 | 0 | 0 | 0 | ? |  | [link](http://lichenportal.org/portal/taxa/index.php?taxon=127290) |
| Capnodiales | *Comminutispora agavacearum* CBS 619.95 | 1 | 0 | ? | 1 | ? | ? | 0 | ? | 0 | 0 | ? | 60 | Figs. 1–3 |
| Capnodiales | *Conidiocarpus caucasicus* GUMH937 | ? | 0 | 1 | 1 | 0 | ? | 0 | ? | 0 | 0 | ? | 61 | Fig. 10 |
| Incertae sedis | *Coniosporium apollinis* CBS 352.97 | 0 | 0 | ? | ? | ? | ? | 0 | ? | ? | 0 | ? | 62 | Fig. 5 |
| Incertae sedis | *Coniosporium uncinatum* CBS 100212 | 0 | 0 | ? | ? | ? | ? | 0 | ? | ? | 0 | ? | 63 | Pl. 2 |
| Incertae sedis | *Cryomyces antarcticus* CCFEE 536 | 0 | 0 | ? | ? | ? | ? | 0 | ? | ? | 0 | ? | 52 | Fig. 9 |
| Incertae sedis | *Cryomyces minteri* CCFEE 5187 | 0 | 0 | ? | ? | ? | ? | 0 | ? | ? | 0 | ? | 52 | Fig. 10 |
| Ostropales | *Cryptodiscus gloeocapsa* TSB 30770 | 3 | 1 | 0 | 1 | 4 | 0 | 0 | 0 | 0 | 0 | ? | 64 | Fig. 3b |
| Helotiales | *Cudoniella clavus* AFTOL-ID 166 | 3 | 0 | 0 | 1 | 4 | 0 | 0 | 0 | 0 | 0 | ? | 65 | Pl. 36  Fig. 347 |
| Capnodiales | *Davidiella tassiana* DAOM 196248 | 1 | 0 | 1 | 1 | 0 | 0 | 0 | 0 | 0 | 0 | ? | 57 | Fig. 18 |
| Arthoniales | *Dendrographa leucophaea* Ornduff 10070 Duke | 0 | 1 | 0 | 1 | 4 | 0 | 0 | 0 | 0 | 0 | ? | 66 | Figs. 2–3;  9–14 |
| Pertusariales | *Dibaeis baeomyces* Lutzoni 93.08.20(Duke) | 0 | 1 | 0 | 1 | 4 | 0 | 0 | 0 | 0 | 0 | ? |  | [link](http://lichenportal.org/portal/taxa/index.php?taxon=Dibaeis%20baeomyces) |
| Venturiales | *Dibotryon morbosum* EF114694.1 | 1 | 0 | 1 | 1 | 0 | 0 | 0 | 0 | 0 | 0 | ? | 67 | Fig. 5 |
| Pleosporales | *Dictyocheirospora rotunda* MFLUCC 0577 | 3 | 0 | ? | ? | ? | ? | 0 | 0 | ? | 0 | ? | 56 | Fig. 4 |
| Pleosporales | *Dictyosporium elegans* NBRC 32502 | 3 | 0 | ? | ? | ? | ? | 0 | 0 | ? | 0 | ? | 56 | Fig. 10 |
| Ostropales | *Diploschistes cinereocaesius* DUKE 0047509 | 0 | 1 | 0 | ? | 4 | 0 | 0 | 0 | ? | 0 | ? | 68 | derived from genus description |
| Ostropales | *Diploschistes muscorum* Palice 2805 (HB Palice) | 0 | 1 | 0 | ? | 4 | 0 | 0 | 0 | ? | 0 | ? | 68 | derived from genus description |
| Ostropales | *Diploschistes ocellatus* Spain 995 | 0 | 1 | 0 | 1 | 4 | 0 | 0 | 0 | 0 | 0 | ? | 68 | derived from genus description |
| Ostropales | *Diploschistes rampoddensis* AF274094.1 | 0 | 1 | 0 | ? | 4 | 0 | 0 | 0 | ? | 0 | ? | 68 | derived from genus description |
| Ostropales | *Diploschistes thunbergianus* Eldridge 3800(F) | 0 | 1 | 0 | ? | 4 | 0 | 0 | 0 | ? | 0 | ? | 68 | derived from genus description |
| Asterotexiales | *Discopycnothyrium palmae* MFLU13-0485 | 1 | 0 | 2 | ? | 0 | 0 | 1 | ? | ? | 0 | 1 | 69 | Fig. 1 |
| Dothideales | *Dothidea insculpta* CBS 189.58 | 1 | 0 | 1 | 1 | 0 | 0 | 0 | 0 | 0 | 0 | ? | 70 | Fig. 3 |
| Dothideales | *Dothidea sambuci* AFTOL-ID 274 | 1 | 0 | 1 | 1 | 0 | 0 | 0 | 0 | 0 | 0 | ? | 70 | Fig. 2 |
| Incertae sedis | Dothideomycetes sp. CCFEE5416 | 0 | 0 | ? | ? | ? | ? | 0 | ? | ? | 0 | ? | 71 | none |
| Lichenostigmatales | Dothideomycetes sp. CCFEE5460 | 0 | 0 | ? | ? | ? | ? | 0 | ? | ? | 0 | ? | 52 | none |
| Incertae sedis | Dothideomycetes sp. TRN 213 | 0 | 0 | ? | ? | ? | ? | 0 | ? | ? | 0 | ? | 72 | none |
| Dyfrolomycetales | *Dyfrolomyces rhizophorae* JK 5349A | 3 | 0 | 1 | 1 | 0 | 0 | 0 | 0 | 0 | 0 | ? | 73 | Figs. 17–20 |
| Dyfrolomycetales | *Dyfrolomyces tiomanensis* NTOU3636 | 3 | 0 | 1 | 1 | 0 | 0 | 0 | 0 | 0 | 0 | ? | 74 | Figs. 2–4 |
| Ostropales | *Dyplolabia afzelii* Luecking 26509a | 3 | 1 | 0 | 1 | 4 | 1 | 0 | 0 | 0 | 0 | ? | 75,76 | Fig. 2a in ^75^, [link](http://www.seaveyfieldguides.com/Lichens/d_lichen/dyplolabia_afzelii_thin_section.htm) |
| Capnodiales | *Elasticomyces elasticus* CCFEE 5320 | 2 | 0 | ? | ? | ? | ? | 0 | ? | ? | 0 | ? | 71 | Fig. 7 |
| Myriangiales | *Elsinoe veneta* AFTOL-ID 1853 | 1 | 0 | 1 | 1 | 0 | 0 | 0 | ? | 0 | 0 | ? | 77 | p. 129 |
| Eurotiales | *Emericella nidulans* ATCC 10074 | ? | 0 | 3 | 1 | 3 | 0 | 0 | ? | 0 | 0 | ? | 78 | p. 6 |
| Incertae sedis | *Encephalographa elisae* EB 0347 | 0 | 1 | 0 | ? | 2 | 1 | 0 | ? | ? | 0 | ? | 41 | Fig. 3n |
| Verrucariales | *Endocarpon pallidulum*  AFTOL-ID 661 | 0 | 1 | 0 | 1 | 4 | 0 | 0 | 0 | 0 | 0 | ? | 79 | Fig. 3 right |
| Myriangiales | *Endosporium aviarium*  UAMH 10530 | ? | 0 | ? | ? | ? | ? | 0 | ? | ? | 0 | ? | 80 | Figs. 25–37 |
| Myriangiales | *Endosporium populi-tremuloides* UAMH 10529 | ? | 0 | ? | ? | ? | ? | 0 | ? | ? | 0 | ? | 80 | Figs. 1–24 |
| Incertae sedis | *Eremomyces bilateralis* CBS 781.70 | ? | 0 | 3 | 1 | 3 | 0 | 0 | 0 | 0 | 0 | ? | 81 | Fig. 8 |
| Erysiphales | *Erysiphe mori* MUMHS77 | 1 | 0 | 3 | 1 | 3 | 0 | 0 | 0 | 0 | 0 | ? | 82 | treated as similar to Erisyphe sect. Uncinula |
| Chaetothyriales | *Exophiala dermatitidis* AFTOL-ID 668 | ? | 0 | 1 | 1 | 0 | ? | 0 | ? | 0 | 0 | ? | 83 | Fig. 3 |
| Chaetothyriales | *Exophiala pisciphila* AFTOL-ID 669 | ? | 0 | 1 | 1 | 0 | ? | 0 | ? | 0 | 0 | ? | 83 | Fig. 4 |
| Incertae sedis | *Farlowiella carmichaeliana* CBS 164.76 | 3 | 0 | 0 | 1 | 2 | 0 | 0 | 0 | 0 | 0 | ? | 65 | Pl. 11  Fig. 102 |
| Ostropales | *Fissurina insidiosa* AFTOL-ID 1662 | 3 | 1 | 0 | 1 | 2 | 1 | 0 | 0 | 0 | 0 | ? | 84 | Figs. 6A; 8B |
| Ostropales | *Fissurina marginata* DNA3418 | 3 | 1 | 0 | 1 | 2 | 1 | 0 | 0 | 0 | 0 | ? | 84 | Coded similarly to *F. insidiosa* |
| Strigulales | *Flavobathelium epiphyllum* MPN67 | 1 | 1 | 1 | 1 | **0** | 0 | 0 | 0 | 0 | 0 | ? | 85 | Figs. 1–2 |
| Capnodiales | *Fumiglobus pieridicola* UBC F23788 | 1 | 0 | 1 | 1 | 0 | ? | 0 | ? | 0 | 0 | ? | 86 | Figs. 2–3 |
| Monoblastiales | *Funbolia dimorpha* CPC 14170 | 3 | 0 | ? | ? | ? | ? | 0 | ? | ? | 0 | ? | 87 | p.114 |
| Venturiales | *Fusicladium oleaginum* CBS 113427 | 1 | 0 | ? | ? | ? | ? | 0 | ? | ? | 0 | ? | 88 | Fig. 35 |
| Incertae sedis | *Geastrumia polystigmatis* FJ147177.1 | 1 | 0 | ? | 0 | 1 | ? | 0 | ? | 0 | 0 | ? | 89 | Figs. 1–2 |
| Venturiales | *Gibbera conferta* CBS 191.53 | 1 | 0 | 1 | 1 | 3 | 0 | 0 | 0 | 0 | 0 | ? | 90 | Fig. 3 |
| Patellariales | *Glyphium elatum* EB 0342 | 1 | 0 | 0 | 1 | 2 | 0 | 0 | 0 | 0 | 0 | ? | 91 | Figs. 1;7 |
| Ostropales | *Gyalecta hypoleuca* TSB 20801 | 0 | 1 | 0 | 1 | 4 | 0 | 0 | 0 | 0 | 0 | ? | 92 | coded as *G. ulmi* |
| Ostropales | *Gyalecta ulmi* AF465463.1 | 3 | 1 | 0 | 1 | 4 | 0 | 0 | 0 | 0 | 0 | ? | 92 | Figs. 1A–2A |
| Pleosporales | *Halojulella avicenniae* BCC 18422 | 3 | 0 | 1 | 1 | 0 | 0 | 0 | 0 | 0 | 0 | ? | 93 | Figs. 1–11 |
| Pleosporales | *Halojulella avicenniae* JK 5326A | 3 | 0 | 1 | 1 | 0 | 0 | 0 | 0 | 0 | 0 | ? | 93 | Figs. 1–11 |
| Monoblastiales | *Heleiosa barbatula* JK 5548I | 1 | 0 | 1 | 1 | 0 | 0 | 0 | 0 | 0 | 0 | ? | 94 | Figs. 1–12 |
| Tubeufiales | *Helicomyces roseus* AFTOL-ID 1613 | 3 | 0 | 1 | 1 | 0 | 0 | 0 | 0 | 0 | 0 | ? | 95 | Fig. 1H |
| Microthyriales | *Heliocephala gracilis* MUCL 41200 | 3 | 0 | ? | ? | ? | ? | 0 | ? | ? | 0 | ? | 96 | Figs. 10–11 |
| Microthyriales | *Heliocephala zimbabweensis* MUCL 40019 | 3 | 0 | ? | ? | ? | ? | 0 | ? | ? | 0 | ? | 97 | Fig. 1 |
| Asterotexiales | *Hemigrapha atlantica* Ertz 14014 (BR) | 2 | 0 | 2 | 0 | 1 | 1 | 1 | ? | 2 | 0 | ? | 98 | Fig. 5 |
| Capnodiales | *Houjia yanglingensis* YHJN13 | 1 | 0 | ? | ? | ? | ? | 0 | ? | ? | 0 | ? | 99 | Fig. 5 |
| Hysteriales | *Hysterobrevium constrictum* SMH 5211.1 | 1 | 0 | 0 | 1 | 2 | 1 | 0 | 0 | 0 | 0 | ? | 100 | Figs. 5A–5E |
| Hysteriales | *Hysterobrevium smilacis* CBS 114601 | 1 | 0 | 0 | 1 | 2 | 1 | 0 | 0 | 0 | 0 | ? | 100 | Figs. 5F–5I |
| Incertae sedis | *Hysterographium fraxini* CBS 109.43 | 1 | 0 | 0 | 1 | 2 | 1 | 0 | 0 | 0 | 0 | ? | 65 | Fig. 610 |
| Patellariales | *Hysteropatella clavispora* CBS 247.34 | 3 | 0 | 0 | 1 | 2 | 1 | 0 | 0 | 0 | 0 | ? | 101 | Pl. 35 |
| Patellariales | *Hysteropatella* *elliptica* CBS 935.97 | 1 | 0 | 0 | 1 | 2 | 1 | 0 | 0 | 0 | 0 | ? | 102 | As *H.prostii* Fig. 7 |
| Asterotexiales | Inocyclus angularis VIC 39747 | 1 | 0 | 2 | 1 | 2 | 1 | 1 | ? | 1 | 0 | ? | 103 | Figs. 1–2 |
| Pleosporales | *Jalapriya pulchra* AF465463.1 | 3 | 0 | ? | ? | ? | ? | 0 | ? | ? | 0 | ? | 56 | Fig. 14 |
| Capnodiales | *Johansonia chapadiensis* CBS H-20484 | 1 | 0 | 0 | 1 | 4 | 0 | 0 | 0 | 0 | 0 | ? | 104 | Fig. 2 |
| Asterotexiales | *Karschia cezannei* Ertz 19186 (BR) | 3 | 0 | 0 | 1 | 4 | 0 | 0 | 0 | 0 | 0 | ? | 41 | Figs. 4h; 5i |
| Asterotexiales | *Karschia talcophila* Diederich 16749 | 2 | 0 | 0 | 1 | 4 | 0 | 0 | 0 | 0 | 0 | ? | 41 | Figs. 4g; 5n |
| Botryosphaeriales | *Kellermania anomala* CBS 132218 | 3 | 0 | 1 | 1 | 0 | ? | 0 | ? | 0 | 0 | ? | 105 | Based on *K. yuccigena*, Fig. 7 |
| Botryosphaeriales | *Kellermania dasylirionicola* CBS 131720 | 3 | 0 | 1 | 1 | 0 | ? | 0 | ? | 0 | 0 | ? | 105 | Based on *K. yuccigena*, Fig. 7 |
| Botryosphaeriales | *Kellermania yuccifoliorum* CBS 131726 | 3 | 0 | 1 | 1 | 0 | ? | 0 | ? | 0 | 0 | ? | 105 | Based on *K. yuccigena*, Fig. 7 |
| Asterotexiales | *Labrocarpon canariense* Ertz 16907(BR) | 2 | 0 | 0 | ? | 2 | 0 | 0 | 0 | ? | 0 | ? | 41 | Fig. 4i |
| Arthoniales | *Lecanographa amylacea* UPS Thor 26176 | 3 | 1 | 0 | 1 | 4 | 1 | 0 | 0 | 0 | 0 | ? |  | [link](http://www.lichens.lastdragon.org/Lecanographa_amylacea.html) |
| Lecanorales | *Lecanora hybocarpa* AFTOL-ID 639 | 3 | 1 | 0 | 1 | 4 | 0 | 0 | 0 | 0 | 0 | ? |  | [link](http://lichenportal.org/portal/taxa/index.php?taxon=53819) |
| Asterinales | *Lembosia abaxialis* VIC 42825 | 1 | 0 | 2 | 0 | 1 | 1 | 1 | 1 | 1 | 1 | 1 | 25 | Fig. 6 |
| Asterotexiales | *Lembosia albersii* MFLU13-0377 | 1 | 0 | 2 | 0 | 1 | 1 | 1 | 1 | 1 | 1 | 1 | 29 | Fig. 18 |
| Asterotexiales | *Lembosia xyliae* MFLU14-0004 | 1 | 0 | 2 | 0 | 1 | 1 | 1 | 1 | 1 | 1 | 1 | 106 | Fig. 7 |
| Aulographaceae | *Lembosina aulographoides* CBS 143809 | 1 | 0 | 2 | 0 | 1 | 1 | 1 | 3 | 1 | 0 | 4 |  | Personal collection |
| Aulographaceae | *Lembosina* sp. 1 CBS 144007 | 1 | 0 | 2 | 0 | 1 | 1 | 1 | 3 | 1 | 0 | 4 |  | personal collection |
| Aulographaceae | *Lembosina* sp. 2 CBS 143815 | 1 | 0 | 2 | 0 | 1 | 1 | 1 | 3 | 1 | 0 | 4 |  | personal collection |
| Mytilinidiales | *Lepidopterella* *palustris* CBS 459.81 | 3 | 0 | 3 | 1 | 3 | 0 | 0 | 0 | 0 | 0 | ? | 107 | Figs. 71–80 |
| Pleosporales | *Leptosphaeria* *heterospora* CBS 644.86 | 1 | 0 | 1 | 1 | 0 | 0 | 0 | 0 | 0 | 0 | ? | 108 | Figs. 17–23 |
| Capnodiales | *Leptoxyphium fumago* CBS 123.26 | 1 | 0 | 1 | 1 | 0 | 0 | 0 | 0 | 0 | 0 | ? | 109 | Figs. 4–7 |
| Abrothallales | *Lichenoconium lecanorae* JL382-10 | 2 | 0 | 1 | 1 | 0 | ? | 0 | ? | 0 | 0 | ? | 110 | Figs. 5Q–5S |
| Microthyriales | *Lichenopeltella pinophylla* UBC-F33032 | 1 | 0 | 2 | 1 | 0 | 0 | 1 | ? | 2 | 0 | 0 |  | personal collection |
| Lichenostigmatales | *Lichenostigma maureri* Diederich 17326 | 0 | 0 | 0 | 1 | 4 | ? | 0 | 0 | 0 | 0 | ? |  | [link](http://lichenportal.org/portal/taxa/index.php?taxon=52691) |
| Incertae sedis | *Lichenothelia calcarea* L1324 | 0 | 0 | 0 | 1 | 4 | ? | 0 | 0 | 0 | 0 | ? | 111 | Fig. 1 |
| Incertae sedis | *Lichenothelia convexa* L1609 | 0 | 0 | 0 | 1 | 4 | ? | 0 | 0 | 0 | 0 | ? | 111 | Fig. 1 |
| Arthoniales | *Lichinella iodopulchra* AFTOL-ID 896 | 0 | 1 | 0 | 1 | 4 | 0 | 0 | 0 | 0 | 0 | ? |  | [link](http://lichenportal.org/portal/taxa/index.php?taxon=124667) |
| Incertae sedis | *Lineolata rhizophorae* CBS 641.66 | 3 | 0 | 1 | 1 | 0 | 0 | 0 | 0 | 0 | 0 | ? | 112 | Fig. 48 |
| Mytilinidiales | *Lophium elegans* EB 0366 | 1 | 0 | 0 | 1 | 2 | ? | 0 | 0 | 0 | 0 | ? | 113 | Fig. 1 |
| Mytilinidiales | *Lophium mytilinum* AFTOL-ID 1609 | 1 | 0 | 0 | 1 | 2 | ? | 0 | 0 | 0 | 0 | ? | 114 | Fig. 1R; 1X |
| Asterotexiales | *Mahanteshomyces* sp. TH 588 | 1 | 0 | 2 | 0 | 1 | 0 | 1 | 1 | 1 | 1 | 1 | 115 | Fig. 3.80 |
| Monoblastiales | *Megalotremis verrucosa* MPN104 | 0 | 1 | 1 | ? | 0 | 0 | 0 | 0 | ? | 0 | ? | 116 | Based on *M. laterale* and *M. cauliflora*, Figs. 80h–i |
| Arthoniales | *Melarthonis piceae* UPS Thor25995 | 3 | 1 | 0 | 1 | 4 | ? | 0 | 0 | 0 | 0 | ? | 117 | Figs. 4D; 4G |
| Asterotexiales | *Melaspilea lekae* Ertz 17325(BR) | 2 | 0 | 0 | 1 | 4 | 0 | 0 | 0 | 0 | 0 | ? | 41 | Fig. 4r |
| Asterotexiales | *Melaspileopsis* cf. *diplasiospora* Ertz 16625 | 3 | 0 | 0 | 0 | 2 | 1 | 0 | 0 | 0 | 0 | ? | 41 | Figs. 4o; 5s |
| Venturiales | *Metacoleroa dickiei* EF114695.1 | 1 | 0 | 1 | ? | 0 | 0 | 0 | 0 | ? | 0 | ? | 90 | Fig. 8 |
| Capnodiales | *Microcyclospora pomicola* CPC 16173 | 1 | 0 | ? | ? | ? | ? | 0 | ? | ? | 0 | ? | 118 | Fig. 6 |
| Ostropales | *Micropeltis* sp. UBC-F33034 | 1 | 0 | 2 | 0 | 0 | 0 | 0 | 4 | 3 | 0 | ? |  | Personal collection |
| Ostropales | *Micropeltis zingiberacicola* IFRDCC 2264 | 1 | 0 | 2 | 0 | 0 | 0 | 0 | 4 | 3 | 0 | ? | 35 | Fig. 11 |
| Microthyriales | *Microthyrium illicinum* CBS 143808 | 1 | 0 | 2 | 0 | 0 | 0 | 1 | 2 | 2 | 0 | 0 |  | Personal collection |
| Microthyriales | *Microthyrium macrosporum* CBS 143810 | 1 | 0 | 2 | 0 | 0 | 0 | 1 | 2 | 2 | 0 | 0 |  | Personal collection |
| Microthyriales | *Microthyrium microscopicum* CBS 115976 | 1 | 0 | 2 | 0 | 0 | 0 | 1 | 2 | 2 | 0 | ? | 35 | Fig. 3 |
| Incertae sedis | *Minutisphaera fimbriatispora* G155.1 | 3 | 0 | 1 | 1 | **0** | 0 | 0 | 0 | 0 | 0 | ? | 119 | Figs. 22–27 |
| Incertae sedis | *Minutisphaera japonica* KTC 2738 | 3 | 0 | 1 | 1 | **0** | 0 | 0 | 0 | 0 | 0 | ? | 119 | Figs. 3–21 |
| Helotiales | *Mollisia cinerea* AFTOL-ID 76 | 4 | 0 | 0 | 1 | 4 | 0 | 0 | 0 | 0 | 0 | ? | 65 | Pl. 4 Fig. 30 |
| Eurotiales | *Monascus purpureus* AFTOL-ID 426 | ? | 0 | 3 | 1 | 3 | 0 | 0 | 0 | 0 | 0 | ? |  | yeast |
| Helotiales | *Monilinia laxa* CBS 122031 | 4 | 0 | 0 | 1 | 4 | 0 | 0 | 0 | 0 | 0 | ? | 65 | based on *M. cinerea* |
| Asterotexiales | *Morenoina calamicola* MFLUCC 14-1162 | 1 | 0 | 2 | 0 | 1 | 1 | 1 | 1 | 1 | 0 | 1 | 120 | Fig. 2 |
| Monoblastiales | *Musaespora kalbii* MPN243 | 1 | 1 | 1 | 1 | 0 | 0 | 0 | 0 | 0 | 0 | ? | 121 | Figs. 1–2 |
| Muyocopronales | *Muyocopron castanopsis* MFLUCC 14-1108 | 1 | 0 | 2 | 0 | 0 | 0 | 1 | ? | 1 | 0 | ? | 122 | Fig. 2 |
| Muyocopronales | *Muyocopron dipterocarpi* MFLUCC 14-110 | 1 | 0 | 2 | 0 | 0 | 0 | 1 | ? | 1 | 0 | ? | 122 | Fig. 3 |
| Muyocopronales | *Muyocopron garethjonesii* MFLU 16-2664 | 1 | 0 | 2 | 0 | 0 | 0 | 1 | ? | 1 | 0 | ? | 123 | Fig. 2 |
| Muyocopronales | *Muyocopron lithocarpi* MFLUCC 10-0041 | 1 | 0 | 2 | 0 | 0 | 0 | 1 | ? | 1 | 0 | ? | 122 | Fig. 4 |
| Muyocopronales | *Muyocopron lithocarpi* MFLUCC 14-1106 | 1 | 0 | 2 | 0 | 0 | 0 | 1 | ? | 1 | 0 | ? | 122 | Fig. 4 |
| Muyocopronales | *Mycoleptodiscus indicus* UAMH 8520 | 1 | 0 | ? | ? | ? | ? | 1 | ? | ? | 0 | ? | 124 | Fig. 3 |
| Trypetheliales | *Mycomicrothelia hemisphaerica* MPN102 | 3 | 1 | 1 | 1 | 0 | 0 | 0 | 0 | 0 | 0 | ? | 125 | Based on *M.* modesta Fig. 3B |
| Trypetheliales | *Mycomicrothelia* *miculiformis* MPN101B | 3 | 1 | 1 | 1 | 0 | 0 | 0 | 0 | 0 | 0 | ? | 125 | Based on *M. modesta*  Fig. 3B |
| Capnodiales | *Mycosphaerella latebrosa* CBS 687.94 | 1 | 0 | 1 | 1 | 0 | 0 | 0 | 0 | 0 | 0 | ? | 126 | none |
| Asterotexiales | *Mycosphaerella pneumatophorae* AFTOL-ID 762 | 3 | 0 | 1 | 1 | 1 | 0 | 0 | 0 | 0 | 0 | ? | 127 | Fig. 87a |
| Capnodiales | *Mycosphaerella walkeri* CPC 11252 | 1 | 0 | 1 | 1 | 0 | 0 | 0 | 0 | 0 | 0 | ? | 128 | Based on *M. sumatrensis*, Fig. 25 |
| Myriangiales | *Myriangium duriaei* CBS 260.36 | 3 | 0 | 1 | 1 | 0 | 0 | 0 | 0 | 0 | 0 | ? | 116 | Fig. 86 |
| Myriangiales | *Myriangium hispanicum* CBS 247.33 | 3 | 0 | 1 | 1 | 0 | 0 | 0 | 0 | 0 | 0 | ? | 116 | Based on *M. duriaei* |
| Mytilinidiales | *Mytilinidion mytilinellum* CBS 303.34 | 1 | 0 | 0 | 1 | 2 | 0 | 0 | 0 | 0 | 0 | ? | 116 | Fig. 87 |
| Mytilinidiales | *Mytilinidion scolecosporum* CBS 305.34 | 3 | 0 | 0 | 1 | 2 | 0 | 0 | 0 | 0 | 0 | ? | 129 | Pl. 17A |
| Natipusillales | *Natipusilla bellaspora* PE91 1a | 3 | 0 | 3 | 1 | 3 | 0 | 0 | 0 | 0 | 0 | ? | 130 | Figs. 1–12 |
| Natipusillales | *Natipusilla decorospora* AF236 1A | 3 | 0 | 3 | 1 | 3 | 0 | 0 | 0 | 0 | 0 | ? | 131 | Figs. 11–16 |
| Natipusillales | *Natipusilla limonensis* AF286 1A | 3 | 0 | 3 | 1 | 3 | 0 | 0 | 0 | 0 | 0 | ? | 131 | Figs. 17–23 |
| Natipusillales | *Natipusilla* *naponensis* AF217 1A | 3 | 0 | 3 | 1 | 3 | 0 | 0 | 0 | 0 | 0 | ? | 131 | Figs. 24–30 |
| Botryosphaeriales | *Neofusicoccum ribis* AFTOL-ID 1232 | 3 | 0 | 1 | 1 | 0 | 0 | 0 | 0 | 0 | 0 | ? | 132 | Figs. 2-3 |
| Ostropales | *Odontotrema phacidioides* Palice 11440 | 3 | 1 | 0 | 1 | 4 | ? | 0 | 0 | 0 | 0 | ? | 133 | Fig. 3F |
| Arthoniales | *Opegrapha dolomitica* AFTOL-ID 993 | 0 | 1 | 0 | 1 | 2 | 1 | 0 | 0 | 0 | 0 | ? | 15 | Based on *O. varia* Fig. 551 |
| Muyocopronales | *Paramycoleptodiscus albizziae* CPC 27552 | 1 | 0 | ? | ? | ? | ? | 1 | ? | ? | 0 | ? | 134 | p. 370 |
| Asterinales | *Parmularia styracis* VIC 42587 | 1 | 0 | 2 | 1 | 1 | 1 | 1 | ? | 1 | 0 | ? | 25 | Fig.2 |
| Capnodiales | *Passalora fulva* CBS 119.46 | 1 | 0 | ? | ? | ? | ? | 0 | ? | ? | 0 | ? | 135 | Fig.6A |
| Patellariales | *Patellaria atrata* CBS 101060 | 3 | 0 | 0 | 1 | 2 | 0 | 0 | 0 | 0 | 0 | ? | 65 | Pl.5 Fig. 45 |
| Capnodiales | *Peltaster fructicola* JN573665.1 | 1 | 0 | 2 | 0 | 1 | 0 | 0 | ? | ? | 0 | ? | 136 | Figs. 5–8 |
| Capnodiales | *Penidiella columbiana* CBS 486.80 | 1 | 0 | ? | ? | ? | ? | 0 | ? | ? | 0 | ? | 47 | Fig. 8 |
| Pertusariales | *Pertusaria dactylina* AFTOL-ID 224 | 0 | 1 | 0 | 1 | 4 | 0 | 0 | 0 | 0 | 0 | ? | 15 | Fig. 629 |
| Lichenostigmatales | Phaeococcomycetaceae sp. TRN 452 | 0 | 0 | ? | ? | ? | ? | 0 | ? | ? | 0 | ? | 72 | none |
| Lichenostigmatales | Phaeococcomycetaceae sp. TRN 456 | 0 | 0 | ? | ? | ? | ? | 0 | ? | ? | 0 | ? | 72 | none |
| Lichenostigmatales | Phaeococcomycetaceae sp. TRN 529 | 0 | 0 | ? | ? | ? | ? | 0 | ? | ? | 0 | ? | 72 | none |
| Capnodiales | *Phaeophleospora atkinsonii* CBS 124565 | 1 | 0 | 1 | ? | 0 | ? | 0 | ? | ? | 0 | ? | 137 | none |
| Chaetothyriales | *Phaeosaccardinula dendrocalami* IFRDCC 2663 | 1 | 0 | 1 | 1 | 0 | 0 | 0 | 0 | 0 | 0 | ? | 138 | Fig. 3 |
| Chaetothyriales | *Phaeosaccardinula ficus* MFLUCC 10-0009 | 1 | 0 | 1 | 1 | 0 | 0 | 0 | 0 | 0 | 0 | ? | 49 | Fig. 3 |
| Capnodiales | *Phaeotheca fissurella* CBS 520.89 | ? | 0 | ? | ? | ? | ? | 0 | ? | ? | 0 | ? | 139 | Figs. 1–5 |
| Phaeotrichales | *Phaeotrichum benjaminii* CBS 541.72 | ? | 0 | 3 | 1 | 3 | 0 | 0 | 0 | 0 | 0 | ? | 140 | Based on *P. hystricinum*, Figs. 1–11 |
| Capnodiales | *Phragmocapnias asiaticus* MFLUCC10-0062 | 1 | 0 | 1 | 1 | 0 | 0 | 0 | 0 | 0 | 0 | ? | 44 | Fig. 5 |
| Strigulales | *Phyllobathelium anomalum* MPN242 | 1 | 1 | 1 | 1 | 0 | 0 | 0 | 0 | 0 | 0 | ? | 125 | Based on *P. firmum*  Fig. 3M |
| Strigulales | *Phyllobathelium firmum* MPN545 | 1 | 1 | 1 | 1 | 0 | 0 | 0 | 0 | 0 | 0 | ? | 125 | Fig. 3M |
| Caliciales | *Physcia aipolia* AFTOL-ID 84 | 3 | 1 | 0 | 1 | 4 | 0 | 0 | 0 | 0 | 0 | ? | 15 | Fig. 659 |
| Capnodiales | *Piedraia hortae* CBS 480.64 | ? | 0 | 3 | 1 | 3 | 0 | 0 | 0 | 0 | 0 | ? | 141 | Figs. 1–2 |
| Pleosporales | *Pleospora herbarum* CBS 191.86 | 1 | 0 | 1 | 1 | 0 | 0 | 0 | 0 | 0 | 0 | ? | 142 | Fig. 1 |
| Ostropales | *Porina farinosa* MPN35 | 1 | 1 | 1 | 1 | 0 | 0 | 0 | 0 | 0 | 0 | ? | 143 | Fig. 4 |
| Ostropales | *Porina guentheri* AF279405.1 | 1 | 1 | 1 | 1 | 0 | 0 | 0 | 0 | 0 | 0 | ? | 143 | Based on *P. farinosa* Fig. 4 |
| Asterinales | *Prillieuxina baccharidincola* VIC 42817 | 1 | 0 | 2 | 0 | 1 | 0 | 1 | ? | 3 | 0 | 1 | 25 | Fig. 7 |
| Venturiales | *Protoventuria barriae* ATCC 90285 | 1 | 0 | 1 | 1 | ? | 0 | 0 | 0 | 0 | 0 | ? | 144 | Figs.1–6; 7; 9; 10 |
| Incertae sedis | *Pseudeurotium hygrophilum* JQ780654.1 | 0 | 0 | 3 | ? | 3 | ? | 0 | 0 | ? | 0 | ? | 145 | Figs. 1–2 |
| Pleosporales | *Pseudocoleophoma polygonicola* KT 731 | 3 | 0 | 1 | 1 | 0 | 0 | 0 | 0 | 0 | 0 | ? | 146 | Fig. 5 |
| Pleosporales | *Pseudodictyosporium wauense* NBRC 30078 | 3 | 0 | ? | ? | ? | ? | 0 | ? | ? | 0 | ? | 56 | Fig. 15 |
| Hysteriales | *Psiloglonium araucanum* CBS 112412 | 1 | 0 | 0 | 1 | 2 | 1 | 0 | 0 | 0 | 0 | ? | 100 | Figs. 8N–8Q |
| Hysteriales | *Psiloglonium clavisporum* GKM L172A | 1 | 0 | 0 | 1 | 2 | 1 | 0 | 0 | 0 | 0 | ? | 100 | Figs. 8E–8H |
| Pleosporales | *Pyrenophora phaeocomes* AFTOL-ID 283 | 1 | 0 | 1 | 1 | 0 | 0 | 0 | 0 | 0 | 0 | ? | 147 | Figs. 1–11 |
| Pyrenulales | *Pyrenula pseudobufonia* AY640962.1 | 3 | 1 | 1 | 1 | 0 | 0 | 0 | 0 | 0 | 0 | ? |  | [link](http://www.waysofenlichenment.net/lichens/Pyrenula%20pseudobufonia) |
| Pyrenulales | *Pyrgillus javanicus* AFTOL-ID 342 | 3 | 1 | 1 | 1 | 0 | 0 | 0 | 0 | 0 | 0 | ? |  | [link](http://lichenportal.org/portal/taxa/index.php?taxon=56108) |
| Capnodiales | *Racodium rupestre* L346 | 0 | 1 | ? | ? | ? | ? | 0 | ? | ? | 0 | ? |  | [link](http://lichenportal.org/portal/taxa/index.php?taxon=118484) |
| Capnodiales | *Rasutoria tsugae* EF1147 | 1 | 0 | 1 | 1 | 0 | 0 | 0 | 0 | 0 | 0 | ? | 148 | none |
| Capnodiales | *Readeriella mirabilis* CBS 116293 | 1 | 0 | 1 | ? | 0 | ? | 0 | ? | ? | 0 | ? | 47 | Fig. 18 |
| Capnodiales | *Recurvomyces mirabilis* CCFEE 5264 | 0 | 0 | ? | ? | ? | ? | 0 | ? | ? | 0 | ? | 71 | Fig. 1 |
| Arthoniales | *Reichlingia zwackhii* KF707637.1 | ? | 1 | 0 | ? | 2 | ? | 0 | 0 | ? | 0 | ? | 149 | Based on *R. syncesioides*, Figs. 2b;  3a–3c |
| Asterotexiales | *Rhagadolobiopsis thelypteridis* EG 156 | 1 | 0 | 2 | 1 | 1 | 1 | 1 | ? | 1 | 0 | 3 | 150 | Fig. 1–2 |
| Incertae sedis | *Rhexothecium globosum* CBS 955.73 | ? | 0 | 3 | 1 | 3 | 0 | 0 | 0 | 0 | 0 | ? | 151 | Fig. 6 |
| Arthoniales | *Roccella fuciformis* AFTOL-ID 126 | 0 | 1 | 0 | ? | 4 | 0 | 0 | 0 | ? | 0 | ? |  | [link](http://www.lichens.lastdragon.org/Roccella_fuciformis.html) |
| Arthoniales | *Roccellographa cretacea* AFTOL-ID 93 | 0 | 1 | 0 | 1 | 2 | 0 | 0 | 0 | 0 | 0 | ? | 152 | none, following genus description |
| Chaetothyriales | *Sarcinomyces petricola* CBS 101157 | 0 | 0 | ? | ? | ? | ? | 0 | ? | ? | 0 | ? | 153 | Figs. 1–11 |
| Incertae sedis | *Saxomyces alpinus* CCFEE 5466 | 0 | 0 | ? | ? | ? | ? | 0 | ? | ? | 0 | ? | 154 | Figs. 3–4 |
| Incertae sedis | *Saxomyces penninicus* CCFEE 5495 | 0 | 0 | ? | ? | ? | ? | 0 | ? | ? | 0 | ? | 154 | Fig. 5 |
| Arthoniales | *Schismatomma decolorans* DUKE-0047570 | 0 | 1 | 0 | ? | 4 | 0 | 0 | 0 | ? | 0 | ? |  | [link](http://lichensmaritimes.org/index.php?task=fiche&lichen=510) |
| Capnodiales | *Schizothyrium pomi* CBS 228.57 | 1 | 0 | 2 | 0 | 1 | 0 | 0 | 4 | 3 | 0 | ? |  | Based on *Schizothyrium gaultheriae* UBC-F3143 |
| Ostropales | *Scolecopeltidium*  sp. 1 UBC-F33033 | 1 | 0 | 2 | 0 | 0 | 0 | 0 | 4 | 3 | 0 | ? |  | Personal collection |
| Ostropales | *Scolecopeltidium*  sp. 2 UBC-F33035 | 1 | 0 | 2 | 0 | 0 | 0 | 0 | 4 | 3 | 0 | ? |  | Personal collection |
| Capnodiales | *Scorias spongiosa* AFTOL-ID 1594 | 1 | 0 | 1 | 1 | 0 | 0 | 0 | 0 | 0 | 0 | ? | 44 | Figs. 12–13 |
| Lichenostigmatales | *Seuratia millardetii* UBC-F33043 | 1 | ? | 0 | 1 | 4 | 0 | 0 | ? | ? | 0 | ? |  | Personal collection |
| Arthoniales | *Simonyella variegata* AFTOL-ID 80 | 0 | 1 | 0 | ? | 4 | 0 | 0 | 0 | ? | 0 | ? |  | [link1](http://www.tropicallichens.net/3719.html), [link2](https://books.google.ca/books?id=-sH9zc8OLOQC&pg=PA349&lpg=PA349&dq=Simonyella+variegata&source=bl&ots=M2eZ8sP0DL&sig=STeejGmQoWp-CKHVISVFSLPlS7U&hl=fr&sa=X&ved=0CFcQ6AEwC2oVChMIvrS1g76iyAIVQqKICh0TnAC4#v=onepage&q=Simonyella%20variegata&f=false) |
| Ostropales | *Sphaeropeziza arctoalpina* Baloch SW057(S) | 4 | 0 | 0 | 1 | 4 | 0 | 0 | 0 | 0 | 0 | ? | 133 | Fig. 5A |
| Capnodiales | *Sphaerulina polyspora* CBS 354.29 | 1 | 0 | 1 | 1 | 0 | 0 | 0 | 0 | 0 | 0 | ? | 155 | Figs. 8;  11–14; 16-17 |
| Onygenales | *Spiromastix warcupii* AFTOL-ID 430 | 0 | 0 | 3 | 1 | 3 | 0 | 0 | 0 | 0 | 0 | ? | 156 | Based on *S.grisea*, Figs. 1–6 |
| Verrucariales | *Staurothele frustulenta* AFTOL-ID 697 | 0 | 1 | 1 | 1 | 0 | 0 | 0 | 0 | 0 | 0 | ? |  | [link](http://www.lichenology.info/cgi-bin/baseportal.pl?htx=atlas&species~=S&abcspec=S&seeall=) |
| Ostropales | *Stictis radiata* AF356663.1 | 3 | 0 | 0 | 1 | 4 | ? | 0 | 0 | 0 | 0 | ? | 157 | Fig. 2I |
| Asterotexiales | *Stictographa lentiginosa* 47621 | 2 | 0 | 0 | 1 | 2 | 1 | 0 | 0 | 0 | 0 | ? | 41 | Fig. 4q |
| Microthyriales | *Stomiopeltis betulae* CBS 114420 | 1 | 0 | ? | ? | ? | 0 | ? | ? | ? | 0 | ? |  | Never illustrated |
| Capnodiales | *Stomiopeltis versicolor* GA3 23C2b | 1 | 0 | ? | ? | ? | ? | ? | ? | ? | 0 | ? |  | Never illustrated |
| Venturiales | *Strigula jamesii* MPN548 | 1 | 1 | 1 | 1 | **0** | 0 | 0 | 0 | 0 | 0 | ? |  | Based on *S. nemathora*. [link](http://www.habitas.org.uk/lichenireland/species.asp?item=19906) |
| Strigulales | *Strigula nemathora* MPN72 | 1 | 1 | 1 | 1 | **0** | 0 | 0 | 0 | 0 | 0 | ? |  | [link](https://www.anbg.gov.au/abrs/lichenlist/VOLUME%2057/Strigula_nemathora_d.html) |
| Strigulales | *Strigula schizospora* MPN73 | 1 | 1 | 1 | 1 | **0** | 0 | 0 | 0 | 0 | 0 | ? |  | Based on *S. nemathora* |
| Dothideales | *Stylodothis puccinioides* CBS 193.58 | 1 | 0 | 1 | 1 | 0 | 0 | 0 | 0 | 0 | 0 | ? | 158 | *Abb.* 144 |
| Dothideales | *Sydowia polyspora* AFTOL-ID 178 | 1 | 0 | 1 | ? | 0 | ? | 0 | ? | ? | 0 | ? | 65 | Pl. 76  Fig. 778 |
| Venturiales | *Sympoventuria capensis* CBS 120136 | 1 | 0 | 1 | 1 | 0 | 0 | 0 | 0 | 0 | 0 | ? | 159 | Fig. 8 |
| Strigulales | *Taeniolella exilis* CBS122902 | 3 | 0 | ? | ? | ? | ? | 0 | ? | ? | 0 | ? | 160 | Figs. 5–6 |
| Asterotexiales | *Taeniolella hawksworthiana* Common 9199B(BR) | 2 | 0 | ? | ? | ? | ? | 0 | ? | ? | 0 | ? | 160 | Figs. 7-8 |
| Asterotexiales | *Taeniolella punctata* Ertz 17390(BR) | 2 | 0 | ? | ? | ? | ? | 0 | ? | ? | 0 | ? | 160 | Figs. 9–11 |
| Asterotexiales | *Taeniolella pyrenulae* Diederich 17075 | 2 | 0 | ? | ? | ? | ? | 0 | ? | ? | 0 | ? | 160 | Figs. 12–13 |
| Asterotexiales | *Taeniolella* sp. Ertz 11026(BR) | 2 | 0 | ? | ? | ? | ? | 0 | ? | ? | 0 | ? | 160 | Figs. 14–15 |
| Asterotexiales | *Taeniolella toruloides* Diederich 17048 | 2 | 0 | ? | ? | ? | ? | 0 | ? | ? | 0 | ? | 160 | Figs. 16–17 |
| Capnodiales | *Teratosphaeria stellenboschiana* CPC 10886 | 1 | 0 | 1 | ? | 0 | ? | 0 | ? | ? | 0 | ? | 128 | Fig. 5 |
| Lichenostigmatales | Teratosphaeriaceae sp. D007 09 | 0 | 0 | ? | ? | ? | ? | 0 | ? | ? | 0 | ? | 161 | none |
| Asterinales | *Thyrinula eucalypti* CPC 12986 | 1 | 0 | ? | ? | ? | ? | ? | ? | ? | 0 | ? | 162 | Figs. 1–7 |
| Venturiales | *Tothia fuscella* CBS 130266 | 1 | 0 | 2 | 0 | 0 | 0 | 1 | 4 | 1 | 0 | ? | 163 | Fig. 2 |
| Ostropales | *Trapelia placodioides* AF274103 | 0 | 1 | 0 | 1 | 4 | 0 | 0 | 0 | 0 | 0 | ? | 15 | Based on *T. involuta*,  Fig. 841 |
| Phaeotrichales | *Trichodelitschia bisporula* CBS 262.69 | ? | 0 | 1 | 1 | 0 | 0 | 0 | 0 | 0 | 0 | ? |  | Based on *T. minuta*, [link](http://ascofrance.fr/recolte/1384/dothideomycetes-pleosporales-phaeotrichaceae-trichodelitschia-minuta) |
| Trypetheliales | *Trypethelium eluteriae* CBS 132375 | 3 | 1 | 1 | 1 | 0 | 0 | 0 | 0 | 0 | 0 | ? | 33 | Figs. 7J; 8E; 10G; 56B–L |
| Tubeufiales | *Tubeufia cerea* AFTOL-ID 1316 | 3 | 0 | 1 | 1 | 0 | 0 | 0 | 0 | 0 | 0 | ? | 164 | Fig. 23 |
| Tubeufiales | *Tubeufia helicomyces* AFTOL-ID 1580 | 3 | 0 | 1 | 1 | 0 | 0 | 0 | 0 | 0 | 0 | ? | 165 | Figs.1–2 |
| Tubeufiales | *Tubeufia paludosa* CBS 120503 | 3 | 0 | 1 | 1 | 0 | 0 | 0 | 0 | 0 | 0 | ? | 164 | Fig. 22 |
| Microthyriales | *Tumidispora shoreae* MFLUCC 12-0409 | 1 | 0 | 2 | 0 | 0 | 0 | 1 | 2 | 2 | 0 | ? | 106 | Fig. 39 |
| Venturiales | *Tyrannosorus pinicola* AFTOL-ID 1235 | ? | 0 | 1 | 1 | 0 | 0 | 0 | 0 | 0 | 0 | ? | 83 | Figs. 6–10 |
| Umbilicariales | *Umbilicaria mammulata* AFTOL-ID 645 | 0 | 1 | 0 | 1 | 4 | 0 | 0 | 0 | 0 | 0 | ? | 15 | Fig. 865 |
| Capnodiales | *Uwebraunia commune* CBS 110747 | 1 | 0 | ? | ? | ? | ? | 0 | ? | ? | 0 | ? | 166 | Figs. 3–6 |
| Venturiales | *Venturia inaequalis* CBS 594.70 | 1 | 0 | 1 | 1 | 0 | 0 | 0 | 0 | 0 | 0 | ? | 116 | Fig. 131 |
| Venturiales | *Venturia pyrina* ICMP 11032 | 1 | 0 | 1 | 1 | 0 | 0 | 0 | 0 | 0 | 0 | ? |  |  |
| Venturiales | *Veronaeopsis simplex* CBS 588.66 | 1 | 0 | ? | ? | ? | ? | 0 | ? | ? | 0 | ? | 167 | Figs. 17C; 35 |
| Tubeufiales | *Wiesneriomyces conjunctosporus* BCC18525 | 1 | 0 | ? | ? | ? | ? | 0 | ? | ? | 0 | ? | 168 | Figs. 1–11 |
| Capnodiales | *Xenomeris juniperi* EF114709.1 | 1 | 0 | 1 | 1 | ? | ? | 0 | 0 | 0 | 0 | ? | 158 | Based on *X. raetica*,  *Abb.* 177 |
| Capnodiales | *Zasmidium anthuriicola* CBS 118742 | 1 | 0 | ? | ? | ? | ? | ? | ? | ? | 0 | ? | 169 | Based on *Z. litseae*, Fig. 2 |
| Zeloasperisporiales | *Zeloasperisporium cliviae* CPC 25145 | 1 | 0 | ? | ? | ? | ? | ? | ? | ? | 0 | ? | 170 | p. 214 |
| Zeloasperisporiales | *Zeloasperisporium eucalyptorum* CBS 124809 | 1 | 0 | ? | ? | ? | ? | ? | ? | ? | 0 | ? | 37 | Fig. 26 |
| Zeloasperisporiales | *Zeloasperisporium ficusicola* MFLUCC 15-0222 | 1 | 0 | 2 | 0 | ? | 0 | 1 | 2 | 2 | 0 | ? | 171 | Fig. 2 |
| Zeloasperisporiales | *Zeloasperisporium searsiae* CPC 25880 | 1 | 0 | ? | ? | ? | ? | ? | ? | ? | 0 | ? | 172 | p. 280 |
| Zeloasperisporiales | *Zeloasperisporium siamense* IFRDCC 2194 | 1 | 0 | 2 | 0 | ? | 0 | 1 | 2 | 2 | 0 | ? | 35 | Fig. 15 |
| Zeloasperisporiales | *Zeloasperisporium wrightiae* MFLUCC 15-0215 | 1 | 0 | 2 | 0 | ? | 0 | 1 | 2 | 2 | 0 | ? | 171 | Fig. 4 |
|  | missing data (%), living taxa | 7.8 | 0.3 | 21 | 27 | 24 | 29 | 3.1 | 33 | 28 | 0 | 90 |  |  |
|  | missing data (%), fossil taxa | 66 | 66 | 0 | 33 | 0 | 0 | 0 | 33 | 33 | 66 | 66 |  |  |

Notes: Characters are: Sub **=** Sporulation substrate; Lic = Lichenized; Spo = Sporocarp type; Low = Differentiated lower wall; Deh = Dehiscence type; Loc = Locule circumference; Rad = Radiating scutellum; Bra = Scutellum branching and septation; Mar = Appressed margin,; App = Lateral appressoria; Ini = Sporocarp initiation. Coding of characters states follows state definitions in Table 2.

Ref is the number of the reference containing the character information and references are listed in full below.

Illustration in Reference specifies the published figure that provided the characters.

**LITERATURE CITED**

1	Mosier, A. C. *et al.* 2016. Fungi contribute critical but spatially varying roles in nitrogen and carbon cycling in acid mine drainage. *Front Microbiol* **7**, 238.

2	Andersen, M. R. *et al.* 2011. Comparative genomics of citric-acid-producing *Aspergillus niger* ATCC 1015 versus enzyme-producing CBS 513.88. *Genome Res* **21**, 885–897.

3	Gostinčar, C. *et al.* 2014. Genome sequencing of four *Aureobasidium pullulans* varieties: biotechnological potential, stress tolerance, and description of new species. *BMC Genomics* **15**, 549–549.

4	Peter, M. *et al.* 2016. Ectomycorrhizal ecology is imprinted in the genome of the dominant symbiotic fungus *Cenococcum geophilum*. *Nature communications* **7**, 12662.

5	McDonald, T. R., Gaya, E. & Lutzoni, F. 2013. Twenty-five cultures of lichenizing fungi available for experimental studies on symbiotic systems. *Symbiosis* **59**, 165–171.

6	Cooke, I. R. *et al.* 2014. Proteogenomic analysis of the *Venturia pirina* (pear scab fungus) secretome reveals potential effectors. *Journal of proteome research* **13**, 3635–3644.

7	Pérez-Ortega, S., Suija, A., Crespo, A. & de los Ríos, A. 2014. Lichenicolous fungi of the genus *Abrothallus* (Dothideomycetes: Abrothallales ordo nov.) are sister to the predominantly aquatic Janhulales. *Fungal Diversity* **64**, 295–304.

8	Diederich, P. 2011. Description of *Abrothallus parmotrematis* sp. nov. (lichenicolous Ascomycota). *Bulletin de la Société des naturalistes luxembourgeois* **112**, 25–34.

9	Stenroos, S. *et al.* 2010. Multiple origins of symbioses between ascomycetes and bryophytes suggested by a five-gene phylogeny. *Cladistics* **26**, 281–300.

10	Sherwood, M. A. 1977. The ostropalean fungi. *Mycotaxon* **5**, 1–277.

11	Baker, B. J., Lutz, M. A., Dawson, S. C., Bond, P. L. & Banfield, J. F. 2004. Metabolically active eukaryotic communities in extremely acidic mine drainage. *Applied and Environmental Microbiology* **70**, 6264–6271.

12	Riddle, L. W. 1920. Observations on the genus *Acrospermum*. *Mycologia* **12**, 175–181.

13	Kohlmeyer, J. & Schatz, S. 1985. *Aigialus* gen.nov. (Ascomycetes) with two new marine species from mangroves. *Transactions of the British Mycological Society* **85**, 699–707.

14	Woudenberg, J. H. C., Groenewald, J. Z., Binder, M. & Crous, P. W. 2013. *Alternaria* redefined. *Studies in Mycology* **75**, 171–212.

15	Brodo, I. M., Sharnoff, S. D. & Sharnoff, S. *Lichen of North America*. (Yale University Press, 2001).

16	Pentecost, A. 2014. Growth and development of ascomata in two species of Arthoniales, *Arthonia calcarea* and *Alyxoria varia* (Lichenized Ascomycota: Arthoniaceae and Lecanographaceae). *Nova Hedwigia* **98**, 41–49.

17	Nelsen, M. *et al.* 2011. New insights into relationships of lichen-forming Dothideomycetes. *Fungal Diversity* **51**, 155–162.

18	Sartoris, G. B. & Kauffman, C. 1925. The development and taxonomic position of *Apiosporina collinsii*. *Papers of the Michigan Academy of Science, Arts and Letters* **5**, 149–162.

19	Sundin, R. & Tehler, A. 1998. Phylogenetic studies of the genus *Arthonia*. *The Lichenologist* **30**, 381–413.

20	Egidi, E. *et al.* 2014. Phylogeny and taxonomy of meristematic rock-inhabiting black fungi in the Dothideomycetes based on multi-locus phylogenies. *Fungal Diversity* **65**, 127–165.

21	de Diego Candela, J. *et al.* 2010. Endocarditis caused by *Arthrographis kalrae*. *The Annals of Thoracic Surgery* **90**, e4–e5.

22	Crous, P. W. *et al.* 2014. Fungal Planet description sheets: 281–319. *Persoonia* **33**, 212–289.

23	Kohlmeyer, J. 1986. *Ascocratera manglicola* gen. et sp. nov. and key to the marine Loculoascomycetes on mangroves. *Canadian Journal of Botany* **64**, 3036–3042.

24	Hofmann, T., Kirschner, R. & Piepenbring, M. 2010. Phylogenetic relationships and new records of Asterinaceae (Dothideomycetes) from Panama. *Fungal Diversity* **43**, 39–53.

25	Guatimosim, E. *et al.* 2014. Towards a phylogenetic reappraisal of Parmulariaceae and Asterinaceae (Dothideomycetes). *Persoonia-Molecular Phylogeny and Evolution of Fungi*.

26	Hyde, K. D. *et al.* 2016. Fungal diversity notes 367–490: taxonomic and phylogenetic contributions to fungal taxa. *Fungal Diversity* **80**, 1–270.

27	Hofmann, T. A. & Piepenbring, M. 2008. New species and records of *Asterina* from Panama. *Mycological Progress* **7**, 87–98.

28	Hofmann, T. A. & Piepenbring, M. 2011. Biodiversity of *Asterina* species on neotropical host plants: New species and records from Panama. *Mycologia* **103**, 1284–1301.

29	Hongsanan, S. *et al.* 2014. Revision of genera in Asterinales. *Fungal Diversity* **68**, 1–68.

30	Guerrero, Y., Hofmann, T. A., Williams, C., Thines, M. & Piepenbring, M. 2011. *Asterotexis cucurbitacearum*, a poorly known pathogen of Cucurbitaceae new to Costa Rica, Grenada and Panama. *Mycology* **2**, 87–90.

31	Liu, J.-K. *et al.* 2011. *Astrosphaeriella* is polyphyletic, with species in *Fissuroma* gen. nov., and *Neoastrosphaeriella* gen. nov. *Fungal Diversity* **51**, 135–154.

32	Nelsen, M. P. *et al.* 2014. Elucidating phylogenetic relationships and genus-level classification within the fungal family Trypetheliaceae (Ascomycota: Dothideomycetes). *Taxon* **63**, 974–992.

33	Aptroot, A. & Lücking, R. 2016. A revisionary synopsis of the Trypetheliaceae (Ascomycota: Trypetheliales). *The Lichenologist* **48**, 763–982.

34	Arx, A. J. v. & Müller, E. 1960. Über die neue Ascomycetengattung *Aulographina*. *Revue de Mycologie* **14**, 330–333.

35	Wu, H.-X. *et al.* 2011. A reappraisal of Microthyriaceae. *Fungal Diversity* **51**, 189–248.

36	Manamgoda, D. S. *et al.* 2014. The genus *Bipolaris*. *Studies in Mycology* **79**, 221–288.

37	Cheewangkoon, R. *et al.* 2009. Myrtaceae, a cache of fungal biodiversity. *Persoonia* **23**, 55–85.

38	Crous, P. W. *et al.* 2016. Fungal Planet description sheets: 469–557. *Persoonia*.

39	Cheewangkoon, R., Groenewald, J., Hyde, K., To-anun, C. & Crous, P. 2012. Chocolate spot disease of *Eucalyptus*. *Mycological Progress* **11**, 61–69.

40	Slippers, B. *et al.* 2004. Combined multiple gene genealogies and phenotypic characters differentiate several species previously identified as *Botryosphaeria dothidea*. *Mycologia* **96**, 83–101.

41	Ertz, D. & Diederich, P. 2015. Dismantling Melaspileaceae: a first phylogenetic study of *Buelliella*, *Hemigrapha*, *Karschia*, *Labrocarpon* and *Melaspilea*. *Fungal Diversity* **71**, 141–164.

42	Sugiyama, J. & Amano, N. 1987. Two metacapnodiaceous sooty moulds from Japan: their identity and behavior in pure culture. *Pleomorphic fungi: the diversity and its taxonomic implications*, 141–156.

43	Seifert, K. A., Morgan-Jones, G., Gams, W. & Kendrick, B. *The genera of hyphomycetes*. (CBS-KNAW Fungal Biodiversity Centre Utrecht, 2011).

44	Chomnunti, P. *et al.* 2011. Capnodiaceae. *Fungal Diversity* **51**, 103–134.

45	Untereiner, W. A. 1997. Taxonomy of selected members of the ascomycete genus *Capronia* with notes on anamorph-teleomorph connections. *Mycologia* **89**, 120–131.

46	Kohlmeyer, J. 1985. *Caryosporella rhizophorae* gen. et sp. nov. (Massariaceae), a marine ascomycete from *Rhizophora* *mangle*. *Proceedings: Plant Sciences* **94**, 355–361.

47	Crous, P. W., Braun, U. & Groenewald, J. Z. 2007. *Mycosphaerella* is polyphyletic. *Studies in Mycology* **58**, 1–32.

48	Massicotte, H. B., Trappe, J. M., Peterson, R. L. & Melville, L. H. 1992. Studies on *Cenococcum geophilum*. II. Sclerotium morphology, germination, and formation in pure culture and growth pouches. *Canadian Journal of Botany* **70**, 125–132.

49	Chomnunti, P. *et al.* 2012. Phylogeny of Chaetothyriaceae in northern Thailand including three new species. *Mycologia* **104**, 382–395.

50	Ainsworth, M. A., Taylor, S. & Cannon, P. F. 2015. Following in the footsteps of Dickie and Leighton: some rarely recorded microfungi on twinflower leaves including *Ceramothyrium linnaeae*, new to Britain. *Field Mycology* **16**, 5–11.

51	Groenewald, J. *et al.* 2013. Species concepts in *Cercospora*: spotting the weeds among the roses. *Studies in Mycology* **75**, 115–170.

52	Selbmann, L., de Hoog, G. S., Mazzaglia, A., Friedmann, E. I. & Onofri, S. 2005. Fungi at the edge of life: cryptoendolithic black fungi from Antarctic desert. *Studies in Mycology*, 1–32.

53	Vikram, H. C., Raj, N. M. & Mathew, D. 2018. New record of *Chaetasbolisia erysiphoides* from cold arid soils of Zanskar (Kargil), India. *Current science* **114**, 25–27.

54	Hongsanan, S., Chomnumti, P., Crous, P. W., Chukeatirote, E. & Hyde, K. D. 2014. Introducing *Chaetothyriothecium*, a new genus of Microthyriales *Phytotaxa* **161**, 157–164.

55	Liu, J. K. *et al.* 2015. Fungal diversity notes 1–110: taxonomic and phylogenetic contributions to fungal species. *Fungal Diversity* **72**, 1–197.

56	Boonmee, S. *et al.* 2016. Dictyosporiaceae fam. nov. *Fungal Diversity*, 1–26.

57	Schubert, K. *et al.* 2007. Biodiversity in the *Cladosporium herbarum* complex (Davidiellaceae, Capnodiales), with standardisation of methods for *Cladosporium* taxonomy and diagnostics. *Studies in Mycology* **58**, 105–156.

58	Sherwood, M. A. 1980. Taxonomic studies in the Phacidales: the genus *Coccomyces* (Rhytismataceae). *Occasional Papers of the Farlow Herbarium of Cryptogamic Botany*, 1–120.

59	Damm, U., Fourie, P. H. & Crous, P. W. 2010. *Coniochaeta* (*Lecythophora*), *Collophora* gen. nov. and *Phaeomoniella* species associated with wood necroses of *Prunus* trees. *Persoonia : Molecular Phylogeny and Evolution of Fungi* **24**, 60–80.

60	Ramaley, A. W. 1996. *Comminutispora* gen. nov. and its *Hyphospora* gen. nov. anamorph. *Mycologia* **88**, 132–136.

61	Bose, T. *Taxonomy and phylogeny of mitosporic Capnodiales and description of a new sooty mold species, Fumiglobus pieridicola, from British Columbia, Canada* MSc thesis, University of British Columbia, (2013).

62	Sterflinger, K. *et al.* 1997. *Coniosporium perforans* and *C. apollinis*, two new rock-inhabiting fungi isolated from marble in the Sanctuary of Delos (Cyclades, Greece). *Antonie van Leeuwenhoek* **72**, 349–363.

63	De Leo, F., Urzi, C. & De Hoog, G. 1999. Two *Coniosporium* species from rock surface. *Studies in Mycology*, 70–79.

64	Baloch, E., Gilenstam, G. & Wedin, M. 2009. Phylogeny and classification of *Cryptodiscus*, with a taxonomic synopsis of the Swedish species. *Fungal Diversity* **38**.

65	Ellis, M. B. & Ellis, J. P. *Microfungi on land plants, an identification handbook*. (The Richmond Publishing Co. Ltd., 1997).

66	Sundin, R. & Tehler, A. 1996. The genus *Dendrographa* (Roccellaceae). *The Bryologist* **99**, 19–31.

67	Zhang, Y. *et al.* 2011. A molecular, morphological and ecological re-appraisal of Venturiales―a new order of Dothideomycetes. *Fungal Diversity* **51**, 249–277.

68	Fernández-Brime, S., Llimona, X., Lutzoni, F. & Gaya, E. 2013. Phylogenetic study of *Diploschistes* (lichen-forming Ascomycota: Ostropales: Graphidaceae), based on morphological, chemical, and molecular data. *Taxon* **62**, 267–280.

69	Hongsanan, S. *et al.* 2016. *Discopycnothyrium palmae* gen. & sp. nov. (Asterinaceae). *Mycotaxon* **131**, 859–869.

70	Thambugala, K. M. *et al.* 2014. Dothideales. *Fungal Diversity* **68**, 105–158.

71	Selbmann, L. *et al.* 2008. Drought meets acid: three new genera in a dothidealean clade of extremotolerant fungi. *Studies in Mycology* **61**, 1–20.

72	Ruibal, C., Platas, G. & Bills, G. F. 2008. High diversity and morphological convergence among melanised fungi from rock formations in the Central Mountain System of Spain. *Persoonia* **21**, 93–110.

73	Hyde, K. D. 1992. The Genus *Saccardoella* from Intertidal Mangrove Wood. *Mycologia* **84**, 803–810.

74	Pang, K.-L. *et al.* 2013. Dyfrolomycetaceae, a new family in the Dothideomycetes, Ascomycota. *Cryptogamie, Mycologie* **34**, 223–232.

75	Rivas Plata, E. *et al.* 2013. A molecular phylogeny of Graphidaceae (Ascomycota, Lecanoromycetes, Ostropales) including 428 species. *MycoKeys* **6**, 55–94.

76	Baloch, E., Lucking, R., Lumbsch, H. T. & Wedin, M. 2010. Major clades and phylogenetic relationships between lichenized and non-lichenized lineages in Ostropales (Ascomycota: Lecanoromycetes). *Taxon* **59**, 1483–1494.

77	Jayawardena, R. S. *et al.* 2014. A re-assessment of Elsinoaceae (Myriangiales, Dothideomycetes). *2014* **176**, 19.

78	Gugnani, H. C. 2003. Ecology and taxonomy of pathogenic aspergilli. *Front Biosci* **8**, 346.

79	Lendemer, J. C. 2007. The Occurrence of *Endocarpon pallidulum* and *Endocarpon petrolepideum* in Eastern North America. *Evansia* **24**, 103–107.

80	Tsuneda, A., Davey, M. L., Hambleton, S. & Currah, R. S. 2008. *Endosporium*, a new endoconidial genus allied to the Myriangiales. *Botany* **86**, 1020–1033.

81	Malloch, D. & Sigler, L. 1988. The Eremomycetaceae (Ascomycotina). *Canadian Journal of Botany* **66**, 1929–1932.

82	Braun, U. *et al.* 2006. Phylogeny and taxonomy of powdery mildew fungi of *Erysiphe* sect. Uncinula on *Carpinus* species. *Mycological Progress* **5**, 139–153.

83	Untereiner, W. A., Straus, N. A. & Malloch, D. 1995. A molecular-morphotaxonomic approach to the systematics of the Herpotrichiellaceae and allied black yeasts. *Mycological Research* **99**, 897–913.

84	Hayward, G. C. 1977. Taxonomy of the lichen families Graphidaceae and Opegraphaceae in New Zealand. *New Zealand Journal of Botany* **15**, 565–584.

85	Lücking, R., Aptroot, A. & Thor, G. 1997. New species or interesting records of foliicolous lichens. II. *Flavobathelium epiphyllum* (lichenized ascomycetes: Melanommatales). *Lichenologist* **29**, 221–228.

86	Bose, T., Reynolds, D. R. & Berbee, M. L. 2014. Common, unsightly and until now undescribed: *Fumiglobus pieridicola* sp. nov., a sooty mold infesting *Pieris japonica* from western North America. *Mycologia* **106**, 746–756.

87	Crous, P. W. *et al.* 2011. Fungal Planet description sheets: 69–91. *Persoonia : Molecular Phylogeny and Evolution of Fungi* **26**, 108–156.

88	Schubert, K., Rischel, A. & Braun, U. 2013. A monograph of *Fusicladium* s. lat. (hyphomycetes). *Schlechtendalia* **9**, 1–132.

89	Pirozynski, K. A. 1971. Note on *Geastrumia polystigmatis*. *Mycologia* **63**, 897–901.

90	Barr, M. E. 1968. The Venturiaceae in North America. *Canadian Journal of Botany* **46**, 799–864.

91	Boehm, E. W., Marson, G., Mathiassen, G. H., Gardiennet, A. & Schoch, C. L. 2015. An overview of the genus *Glyphium* and its phylogenetic placement in Patellariales. *Mycologia* **107**, 607–618.

92	Kauff, F. & Büdel, B. 2005. Ascoma ontogeny and apothecial anatomy in the Gyalectaceae (Ostropales, Ascomycota) support the re-establishment of the Coenogoniaceae. *The Bryologist* **108**, 272–281.

93	Hyde, K. D. 1992. *Julella avicenniae* (Borse) comb. nov. (Thelenellaceae) from intertidal mangrove wood and miscellaneous fungi from the NE coast of Queensland. *Mycological Research* **96**, 939–942.

94	Kohlmeyer, J., Volkmann-Kohlmeyer, B. & Eriksson, O. E. 1996. Fungi on *Juncus roemerianus*. 8. New bitunicate ascomycetes. *Canadian Journal of Botany* **74**, 1830–1840.

95	Tsui, C. K. M. & Berbee, M. L. 2006. Phylogenetic relationships and convergence of helicosporous fungi inferred from ribosomal DNA sequences. *Molecular Phylogenetics and Evolution* **39**, 587–597.

96	Castañeda Ruiz, R. F. *Deuteromycotina de Cuba. Hyphomycetes III*. (Instituto de Investigaciones, “Alejandro de Humboldt”, 1985).

97	Decock, C., Robert, V. & Masuka, A. J. 1998. *Heliocephala zimbabweensis* sp. nov. from Southern Africa. *Mycologia* **90**, 330–333.

98	Diederich, P. & Wedin, M. 2000. The species of *Hemigrapha* (lichenicolous Ascomycetes, Dothideales) on Peltigerales. *Nordic Journal of Botany* **20**, 203–214.

99	Yang, H. L. *et al.* 2010. Novel fungal genera and species associated with the sooty blotch and flyspeck complex on apple in China and the USA. *Persoonia* **24**, 29–37.

100	Boehm, E. W. A. *et al.* 2009. A molecular phylogenetic reappraisal of the Hysteriaceae, Mytilinidiaceae and Gloniaceae (Pleosporomycetidae, Dothideomycetes) with keys to world species. *Studies in Mycology* **64**, 49–83–S43.

101	Seaver, F. J. 1910. Iowa Discomycetes. *Bulletin of the Laboratories of Natural History of the State University of Iowa* **6**, 41–131.

102	Yacharoen, S. *et al.* 2015. Patellariaceae revisited. *Mycosphere* **6**, 290–326.

103	Guatimosim, E., Schwartsburd, P. B. & Barreto, R. W. 2014. A new *Inocyclus* species (Parmulariaceae) on the neotropical fern *Pleopeltis astrolepis*. *IMA Fungus* **5**, 51–55.

104	Crous, P. W., Barreto, R. W., Alfenas, A. C., Alfenas, R. F. & Groenewald, J. Z. 2010. What is *Johansonia*? *IMA fungus* **1**, 117–122.

105	Slippers, B. *et al.* 2013. Phylogenetic lineages in the Botryosphaeriales: a systematic and evolutionary framework. *Studies in Mycology* **76**, 31–49.

106	Ariyawansa, H. A. *et al.* 2015. Fungal diversity notes 111–252—taxonomic and phylogenetic contributions to fungal taxa. *Fungal Diversity* **75**, 27–274.

107	Raja, H. A. & Shearer, C. A. 2008. Freshwater ascomycetes: new and noteworthy species from aquatic habitats in florida. *Mycologia* **100**, 467–489.

108	Ahn, Y.-m. & Shearer, C. A. 1999. Taxonomic revision of *Leptosphaeria vagabunda* and four infraspecific taxa. *Mycologia* **91**, 684–693.

109	Srivastava, R. C. 1982. Notes on two interesting fungi from india. *Archiv für Protistenkunde* **125**, 331–333.

110	Lawrey, J. D. *et al.* 2011. The obligately lichenicolous genus *Lichenoconium* represents a novel lineage in the Dothideomycetes. *Fungal Biology* **115**, 176–187.

111	Muggia, L., Gueidan, C., Knudsen, K., Perlmutter, G. & Grube, M. 2013. The lichen connections of black fungi. *Mycopathologia* **175**, 523–535.

112	Zhang, Y., Crous, P. W., Schoch, C. L. & Hyde, K. D. 2012. Pleosporales. *Fungal Diversity* **53**, 1–221.

113	Mathiassen, G. H., Granmo, A. & Rämä, T. 2015. *Lophium elegans* (Ascomycota), a rare European species.

114	Boehm, E. W. A., Schoch, C. L. & Spatafora, J. W. 2009. On the evolution of the Hysteriaceae and Mytilinidiaceae (Pleosporomycetidae, Dothideomycetes, Ascomycota) using four nuclear genes. *Mycological Research* **113**, 461–479.

115	Hofmann, T. A. *Plant parasitic Asterinaceae and Microthyriaceae from the Neotropics (Panama)* Ph.D. thesis, Johann Wolfgang Goethe-University, (2009).

116	Hyde, K. D. *et al.* 2013. Families of Dothideomycetes. *Fungal Diversity* **63**, 1–313.

117	Frisch, A., Thor, G., Ertz, D. & Grube, M. 2014. The arthonialean challenge: restructuring Arthoniaceae. *Taxon* **63**, 727–744.

118	Frank, J. *et al.* 2010. *Microcyclospora* and *Microcyclosporella*: novel genera accommodating epiphytic fungi causing sooty blotch on apple. *Persoonia* **24**, 93–105.

119	Raja, H. A. *et al.* 2013. Freshwater ascomycetes: *Minutisphaera* (Dothideomycetes) revisited, including one new species from Japan. *Mycologia* **105**, 959–976.

120	Tibpromma, S. *et al.* 2017. Fungal diversity notes 491–602: taxonomic and phylogenetic contributions to fungal taxa. *Fungal Diversity* **83**, 1–261.

121	Lücking, R. & Sérusiaux, E. 1996. *Musaespora kalbii* (lichenized Ascomycetes: Melanommatales), a new foliicolous lichen with a pantropical distribution. *Nordic Journal of Botany* **16**, 661–668.

122	Mapook, A. *et al.* 2016. Muyocopronales, ord. nov.,(Dothideomycetes, Ascomycota) and a reappraisal of *Muyocopron* species from northern Thailand. *Phytotaxa* **265**, 225–237.

123	Tibpromma, S. *et al.* 2016. *Muyocopron garethjonesii* sp. nov. (Muyocopronales, Dothideomycetes) on *Pandanus* sp. *Mycosphere* **7**, 1480–1489.

124	Sutton, B. C. 1973. *Pucciniopsis*, *Mycoleptodiscus* and *Amerodiscosiella*. *Transactions of the British Mycological Society* **60**, 525–536.

125	Nelsen, M. P. *et al.* 2009. Unravelling the phylogenetic relationships of lichenised fungi in Dothideomyceta. *Studies in Mycology* **64**, 135–144–S134.

126	Crous, P. W. *et al.* 2009. Phylogenetic lineages in the Capnodiales. *Studies in Mycology* **64**, 17–47–S17.

127	Kohlmeyer, J. 1966. Neue Meerespilze an Mangroven. *Berichte der Deutschen Botanischen Gesellschaft* **79**, 27–37.

128	Crous, P. W., Wingfield, M. J., Mansilla, J. P., Alfenas, A. C. & Groenewald, J. Z. 2006. Phylogenetic reassessment of *Mycosphaerella* spp. and their anamorphs occurring on *Eucalyptus*. II. *Studies in Mycology* **55**, 99–131.

129	Lohman, M. L. 1932. Three new Species of *Mytilidion* in the proposed subgenus, Lophiopsis. *Mycologia* **24**, 477–484.

130	Raja, H. A., Miller, A. N. & Shearer, C. A. 2012. Freshwater ascomycetes: Natipusillaceae, a new family of tropical fungi, including *Natipusilla bellaspora* sp. nov. from the Peruvian Amazon. *Mycologia* **104**, 569–573.

131	Ferrer, A., Miller, A. N. & Shearer, C. A. 2011. *Minutisphaera* and *Natipusilla*: two new genera of freshwater Dothideomycetes. *Mycologia* **103**, 411–423.

132	Wolf, F. T. & Wolf, F. A. 1939. A study of *Botryosphaeria* *ribis* on willow. *Mycologia* **31**, 217–227.

133	Baloch, E., Gilenstam, G. & Wedin, M. 2013. The relationships of *Odontotrema* (Odontotremataceae) and the resurrected *Sphaeropezia* (Stictidaceae)—new combinations and three new *Sphaeropezia* species. *Mycologia* **105**, 384–397.

134	Crous, P. W. *et al.* 2016. Fungal Planet description sheets: 400–468. *Persoonia* **36**, 316–458.

135	Crous, P. W. 2009. Taxonomy and phylogeny of the genus *Mycosphaerella* and its anamorphs. *Fungal Diversity* **38**, 1–24.

136	Williamson, S. M., Hodges, C. S. & Sutton, T. B. 2004. Re-examination of *Peltaster fructicola*, a member of the apple sooty blotch complex. *Mycologia* **96**, 885–890.

137	Pennycook, S. & McKenzie, E. 2002. *Scoleciasis atkinsonii*, an earlier name for *Phaeophleospora hebes*; and a note on GH Cunningham's epithets *hebe* and *pseudopanax*. *Mycotaxon* **82**, 145–146.

138	Yang, H., Chomnumti, P., Aryiawansa, H., Wu, H.-x. & Hyde, K. D. 2014. The genus *Phaeosaccardinula* (Chaetothyriales) from Yunnan, China, introducing two new species. *Chiang Mai Journal of Science* **41**, 873–884.

139	Tsuneda, A., Tsuneda, I. & Currah, R. S. 2004. Endoconidiogenesis in *Endoconidioma populi* and *Phaeotheca fissurella*. *Mycologia* **96**, 1136–1142.

140	Cain, R. F. 1956. Studies of coprophilous ascomycetes: II *Phaeotrichum*, a new cleistocarpous genus in a new family, and its relationships. *Canadian Journal of Botany* **34**, 675–687.

141	Takashio, M. & Vanbreuseghem, R. 1971. Production of ascospores by *Piedraia hortai* in vitro. *Mycologia* **63**, 612–618.

142	Inderbitzin, P., Mehta, Y. R. & Berbee, M. L. 2009. *Pleospora* species with *Stemphylium* anamorphs: a four locus phylogeny resolves new lineages yet does not distinguish among species in the *Pleospora herbarum* clade. *Mycologia* **101**, 329–339.

143	Nelsen, M. P. *et al.* 2014. Molecular phylogeny reveals the true colours of Myeloconidaceae (Ascomycota: Ostropales). *Australian Systematic Botany* **27**, 38–47.

144	Carris, L. M. & Poole, A. P. 1993. A new species of *Protoventuria* on leaves of *Vaccinium macrocarpon*. *Mycologia* **85**, 93–99.

145	Sogonov, M. V., Schroers, H.-J., Gams, W., Dijksterhuis, J. & Summerbell, R. C. 2005. The hyphomycete *Teberdinia hygrophila* gen. nov., sp. nov. and related anamorphs of *Pseudeurotium* species. *Mycologia* **97**, 695–709.

146	Tanaka, K. *et al.* 2015. Revision of the Massarineae (Pleosporales, Dothideomycetes). *Studies in Mycology* **82**, 75–136.

147	Shoemaker, R. A. 1961. *Pyrenophora phaeocomes* (Reb. ex Fr.) Fr. *Canadian Journal of Botany* **39**, 901–908.

148	Dearness, J. 1924. New and Noteworthy Fungi: III. *Mycologia* **16**, 143–176.

149	Frisch, A., Thor, G. & Sheil, D. 2014. Four new Arthoniomycetes from Bwindi Impenetrable National Park, Uganda–supported by molecular data. *Nova Hedwigia* **98**, 295–312.

150	Guatimosim, E., Pinto, H. J., Barreto, R. W. & Prado, J. 2014. *Rhagadolobiopsis*, a new genus of Parmulariaceae from Brazil with a description of the ontogeny of its ascomata. *Mycologia* **106**, 276–281.

151	Samson, R. A. & Mouchacca, J. 1975. Two new soil-borne cleistothecial ascomycetes. *Canadian Journal of Botany* **53**, 1634–1639.

152	Tehler, A. & Irestedt, M. 2007. Parallel evolution of lichen growth forms in the family Roccellaceae (Arthoniales, Ascomycota). *Cladistics* **23**, 432–454.

153	Wollenzien, U., de Hoog, G. S., Krumbein, W. & Uijthof, J. M. J. 1997. *Sarcinomyces petricola*, a new microcolonial fungus from marble in the Mediterranean basin. *Antonie van Leeuwenhoek* **71**, 281–288.

154	Selbmann, L. *et al.* 2014. Mountain tips as reservoirs for new rock-fungal entities: *Saxomyces* gen. nov. and four new species from the Alps. *Fungal Diversity* **65**, 167–182.

155	Wolf, F. A. 1925. Some undescribed fungi on sourwood, *Oxydendron arboreum* (L.) DC. *Journal of the Elisha Mitchell Scientific Society* **41**, 94–99.

156	Currah, R. & Locquin-Linard, M. 1988. *Spiromastix grisea* sp. nov. and its relationship to other Onygenaceae with helical appendages. *Canadian Journal of Botany* **66**, 1135–1137.

157	Wedin, M., Döring, H. & Gilenstam, G. 2006. *Stictis* s. lat. (Ostropales, Ascomycota) in northern Scandinavia, with a key and notes on morphological variation in relation to lifestyle. *Mycological Research* **110**, 773–789.

158	Müller, E. & von Arx, J. A. 1962. Die Gattungen der didymosporen Pyrenomyceten. *Beiträge zur Kryptogamenflora der Schweiz* **11**, 1–922.

159	Crous, P. W., Mohammed, C., Glen, M., Verkley, G. J. M. & Groenewald, J. Z. 2007. *Eucalyptus* microfungi known from culture. 3. *Eucasphaeria* and *Sympoventuria* genera nova, and new species of *Furcaspora*, *Harknessia*, *Heteroconium* and *Phacidiella*. *Fungal Diversity* **25**, 19–36.

160	Ertz, D. *et al.* 2016. Contribution to the phylogeny and taxonomy of the genus *Taeniolella*, with a focus on lichenicolous taxa. *Fungal biology* **120**, 1416–1447.

161	Ruibal, C. *et al.* 2009. Phylogeny of rock-inhabiting fungi related to Dothideomycetes. *Studies in Mycology* **64**, 123–133.

162	Wall, E. & Keane, P. J. 1984. Leaf spot of *Eucalyptus* caused by *Aulographina eucalypti*. *Transactions of the British Mycological Society* **82**, 257–273.

163	Wu, H., Jaklitsch, W. M., Voglmayr, H. & Hyde, K. D. 2012. Epitypification, morphology, and phylogeny of *Tothia fuscella*. *Mycotaxon* **118**, 203–211.

164	Boonmee, S. *et al.* 2014. Tubeufiales, ord. nov., integrating sexual and asexual generic names. *Fungal Diversity* **68**, 239–298.

165	Webster, J. 1951. Graminicolous pyrenomycetes: I. The conidial stage of *Tubeufia helicomyces*. *Transactions of the British Mycological Society* **34**, 304–308.

166	Crous, P. W., Groenewald, J. Z., Mansilla, J. P., Hunter, G. C. & Wingfield, M. J. 2004. Phylogenetic reassessment of *Mycosphaerella* spp. and their anamorphs occurring on *Eucalyptus*. *Studies in Mycology* **50**, 195–214.

167	Arzanlou, M. *et al.* 2007. Phylogenetic and morphotaxonomic revision of *Ramichloridium* and allied genera. *Studies in Mycology* **58**, 57–93.

168	Kuthubutheen, A. J. & Nawawi, A. 1988. A new species of *Wiesneriomyces* (Hyphomycetes) from submerged decaying leaves. *Transactions of the British Mycological Society* **90**, 619–625.

169	Zhao, W. *et al.* 2016. A new species of *Zasmidium* associated with sooty blotch and flyspeck. *Phytotaxa* **258**, 190–194.

170	Crous, P. W. *et al.* 2015. Fungal Planet description sheets: 320-370. *Persoonia* **34**, 167–266.

171	Hongsanan, S. *et al.* 2015. Zeloasperisporiales ord. nov., and two new species of *Zeloasperisporium*. *Cryptogamie, Mycologie* **36**, 301–317.

172	Crous, P. W. *et al.* 2015. Fungal Planet description sheets: 371–399. *Persoonia* **35**, 264–327.
