## Supplemental Data 2 for "Character evolution of modern fly-speck fungi and implications for interpreting thyriothecial fossils"

Appendix S5. Consensus tree from MrBayes analysis.
Results from four runs of eight chains each, running 160 million generations and sampling every 5000 trees, from which 50% of the samples were discarded as burn-in. Posterior probabilities mapped on branches, clades are labeled and ordered as in Fig. 1.

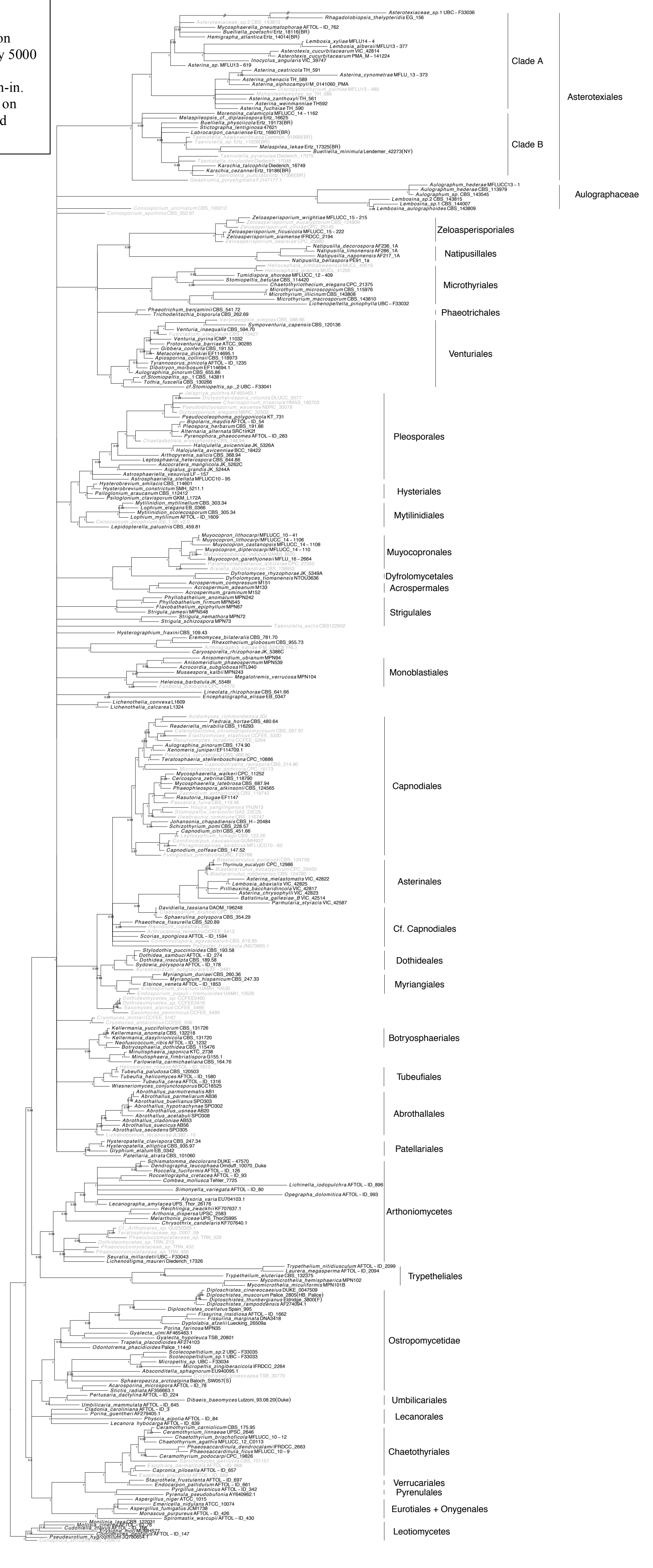

Appendix S6. Maximum likelihood phylogeny labeled with codes for nodes in ancestral state analysis. Pink diamonds indicate taxa forming thyriothecia. Most likely tree obtained out of 4000 independent ML searches of a 4552 bp alignment of LSU and SSU rDNA data for 320 taxa, with bootstrap support > 70% in red and posterior probability > 0.95 in black.

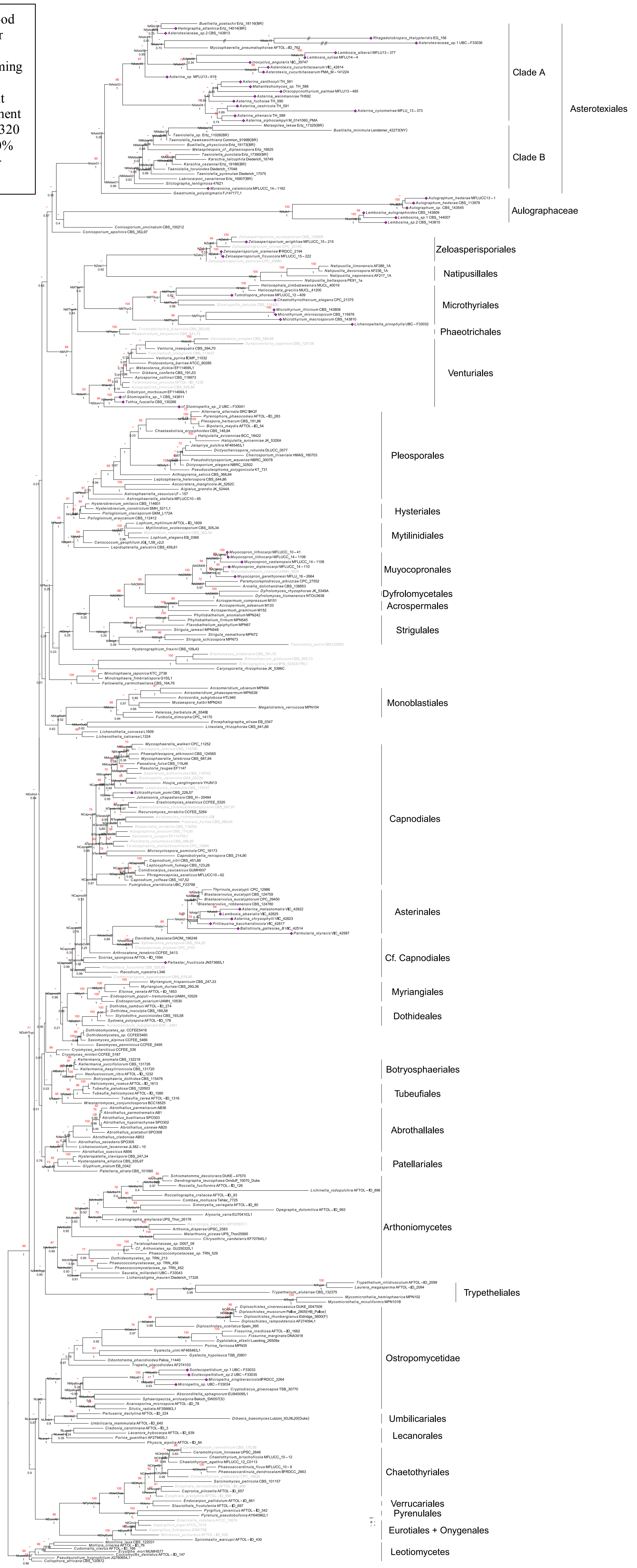

**Appendices S8–S18 Ancestral character state reconstructions**
Ancestral character state reconstructions over most likely topology for 11 morphological characters: proportional likelihoods from MK1 analysis are reported in red pie charts, and posterior probabilities from BayesTraits V3 analysis reported in yellow pie charts. Both were mapped for nodes that could be reconstructed in each tree. In each tree, names in grey indicate taxa that could not be coded for the character, with ancestral states that could not be reconstructed. Bootstrap support > 0.7 is reported in red and posterior probabilities > 0.95 in black.

Appendix S8. Ancestral character state reconstructions for sporulation substrate (Sub).

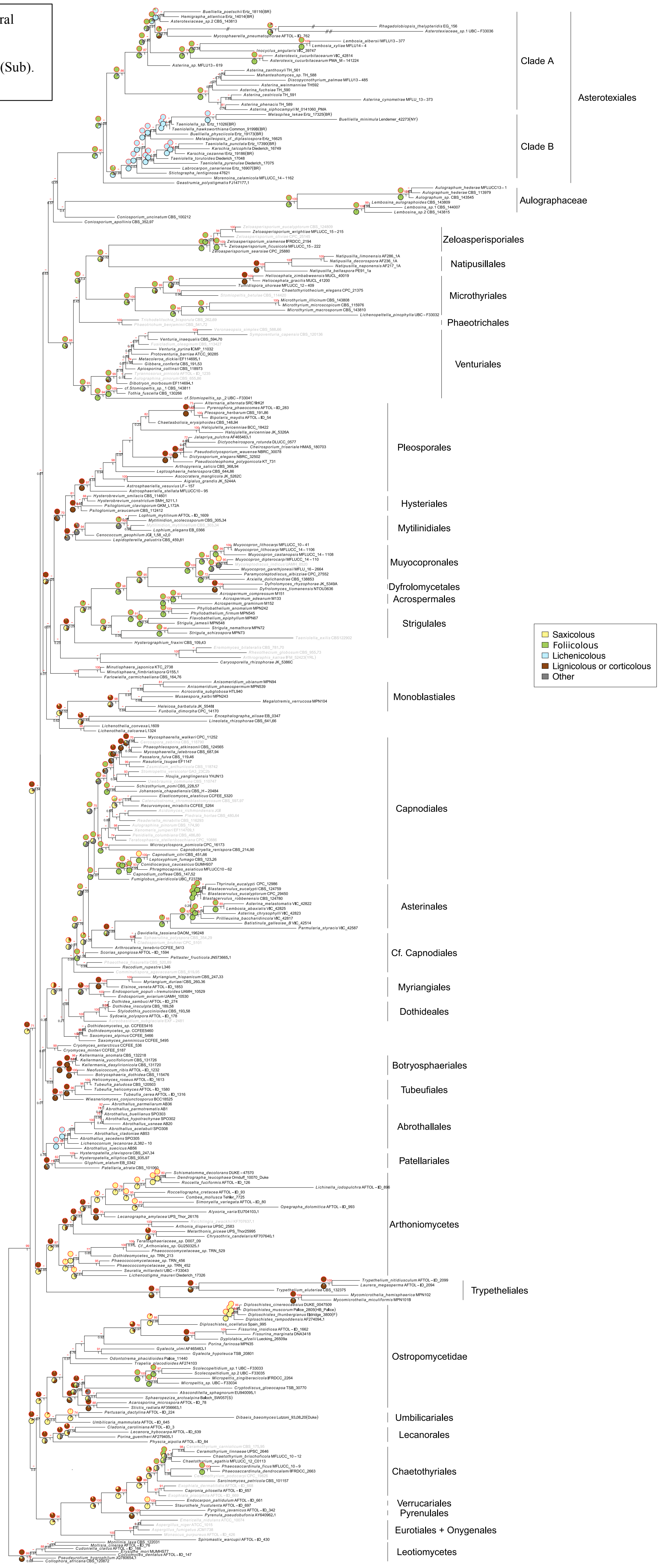

Appendix S9. Ancestral character states reconstruction as lichenized or non-lichenized (Lic).

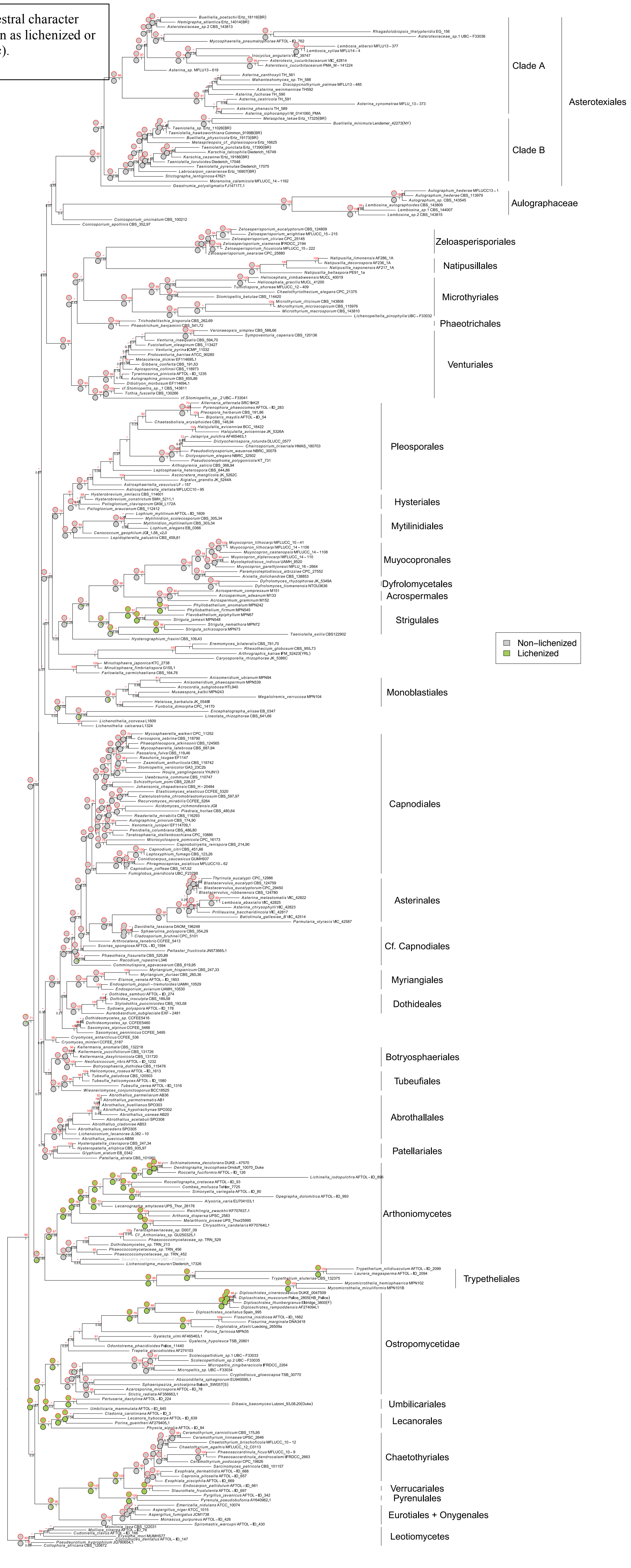

Appendix S10. Ancestral character state reconstruction for sporocarp type (Spo).

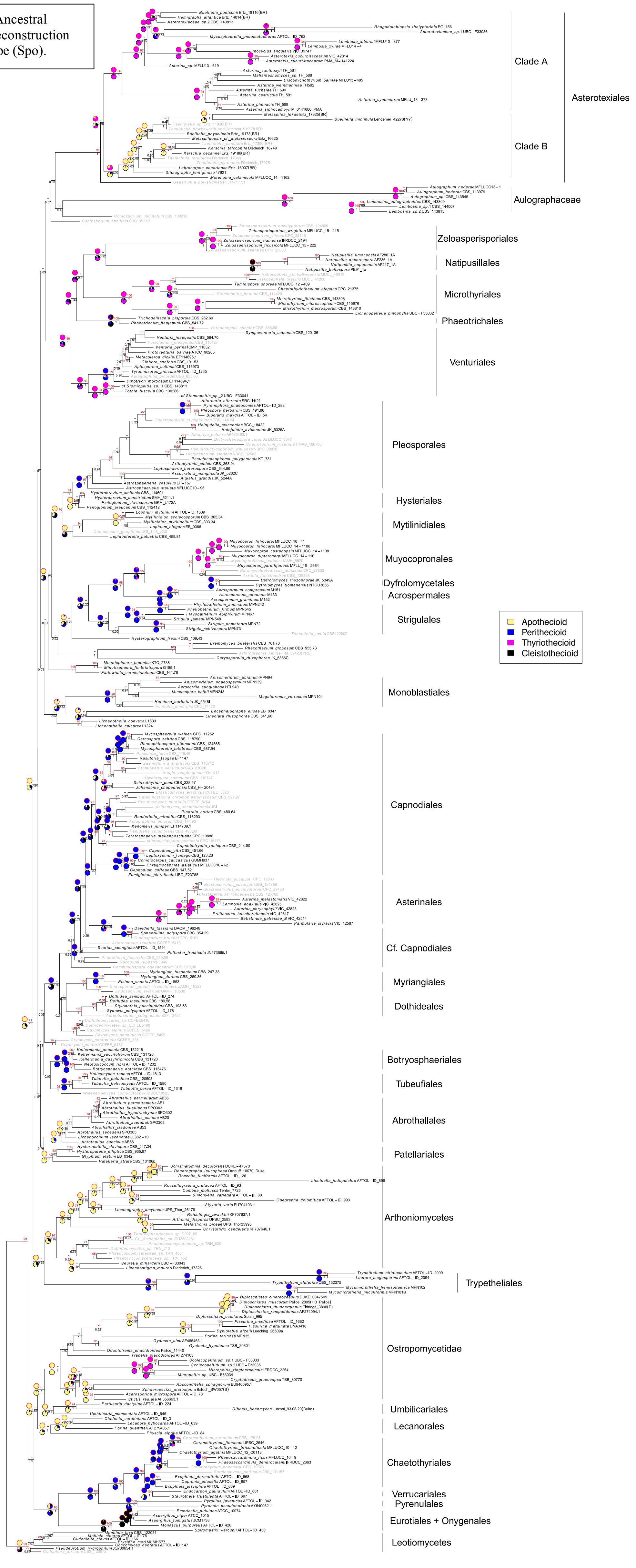

Appendix S11. Ancestral character state reconstruction for differentiated or undifferentiated lower wall of sporocarp (Low).

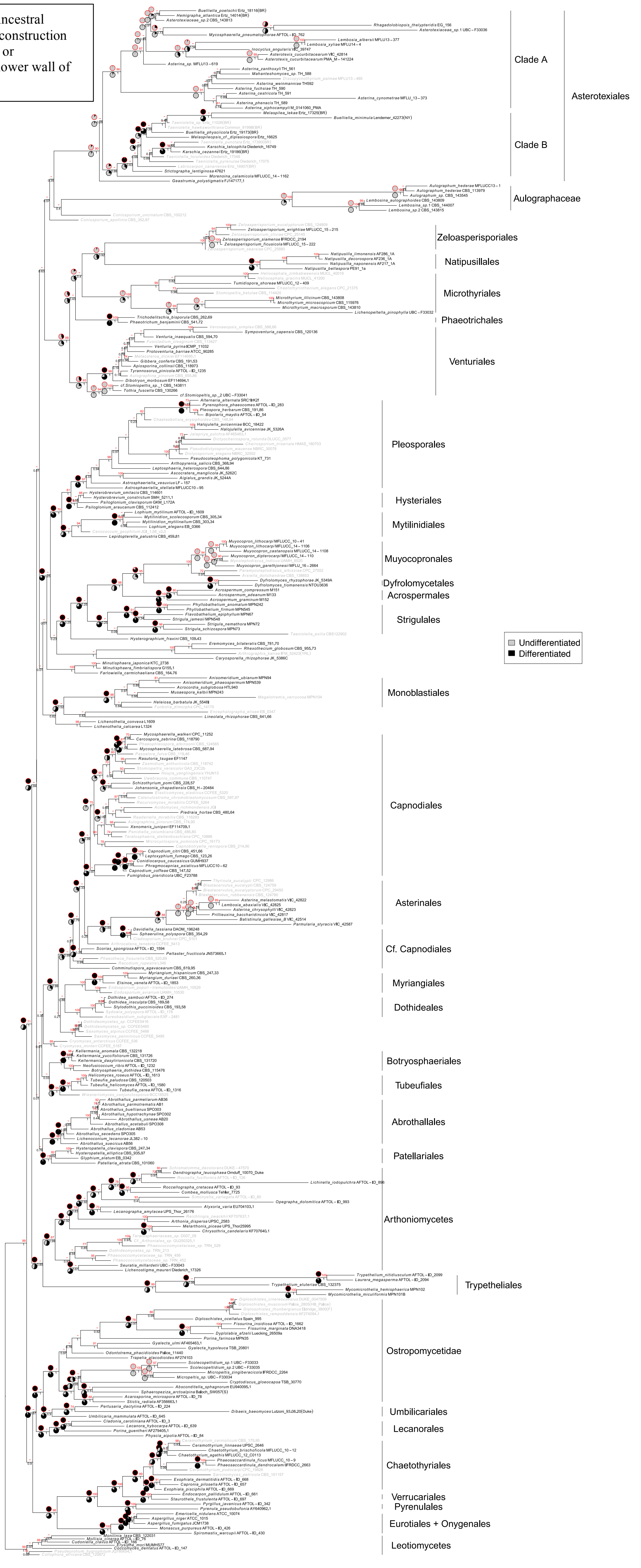

Appendix S12. Ancestral character state reconstruction for sporocarp dehiscence (Deh).

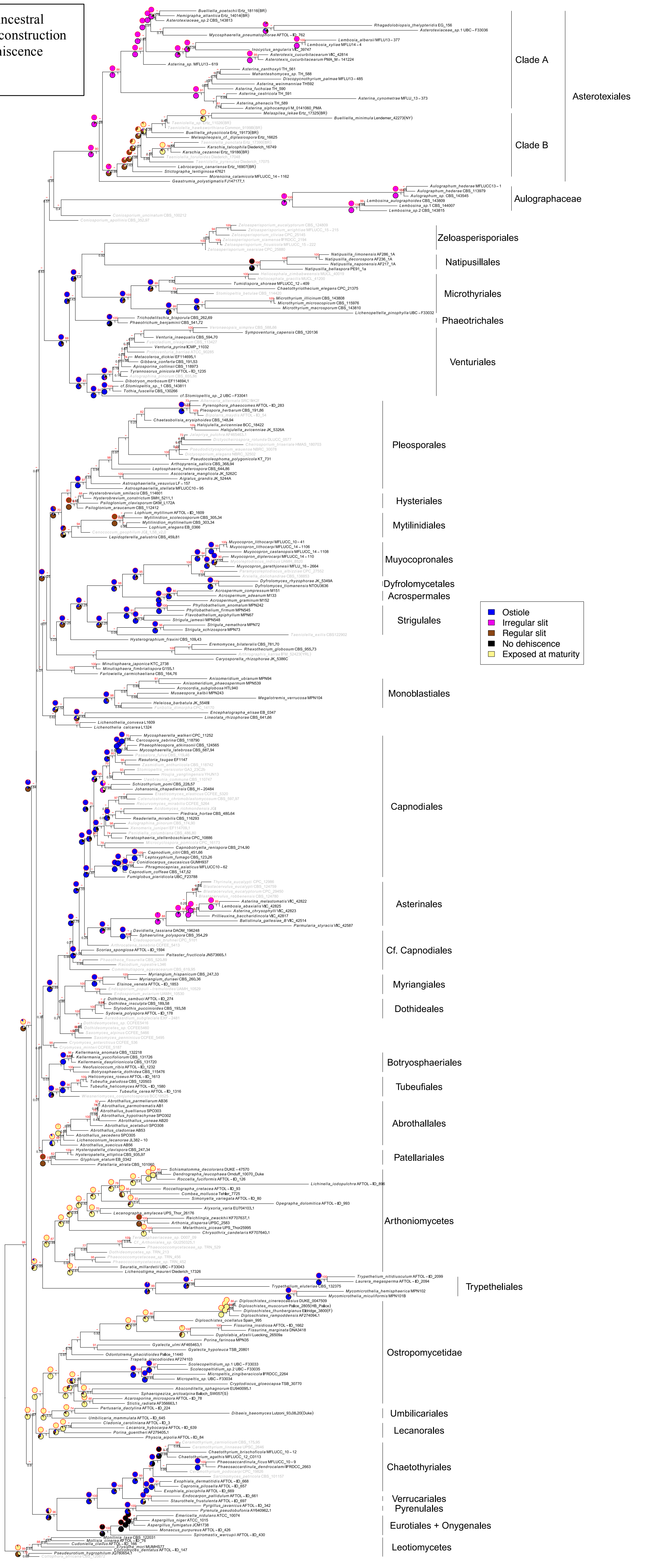

Appendix S13. Ancestral character state reconstruction for circular or elongate locule circumference (Loc).

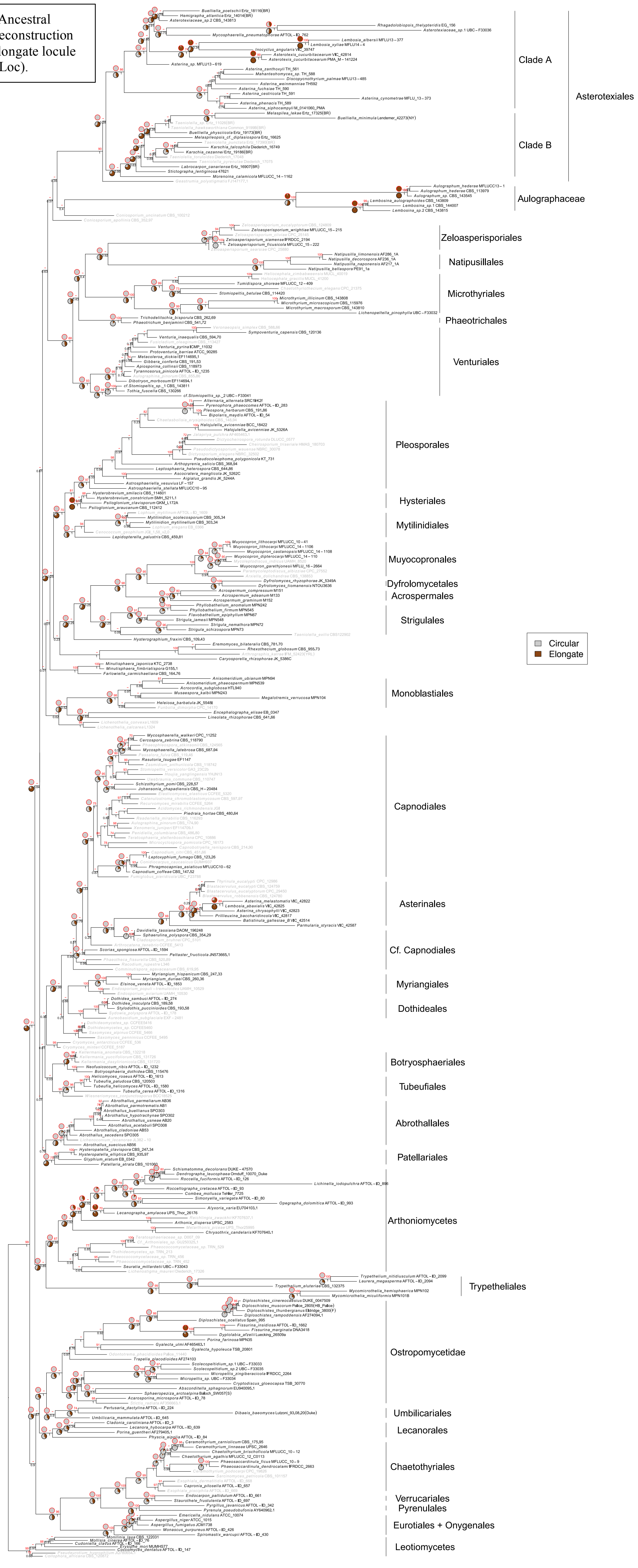

Appendix S14. Ancestral character state reconstruction for presence or absence of a radiate sporocarp (Rad).

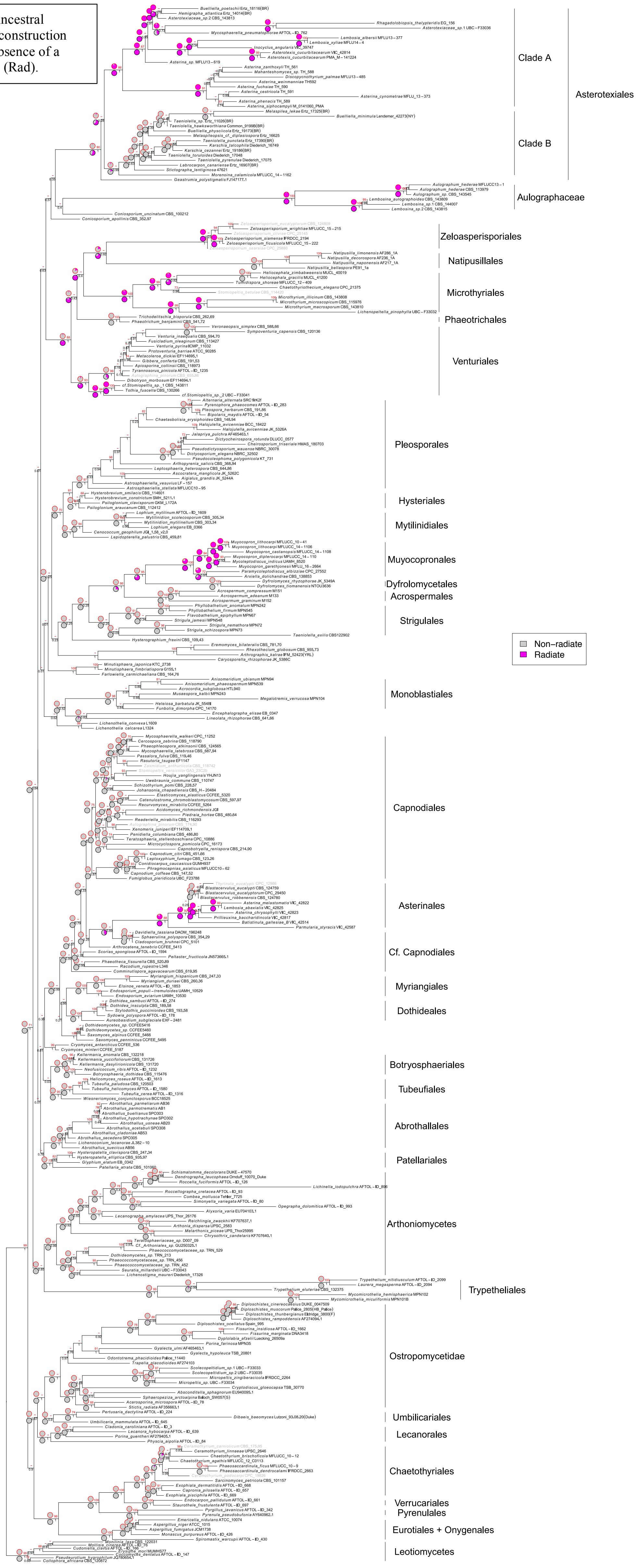

Appendix S15. Ancestral character state reconstruction for scutellum branching (Bra).

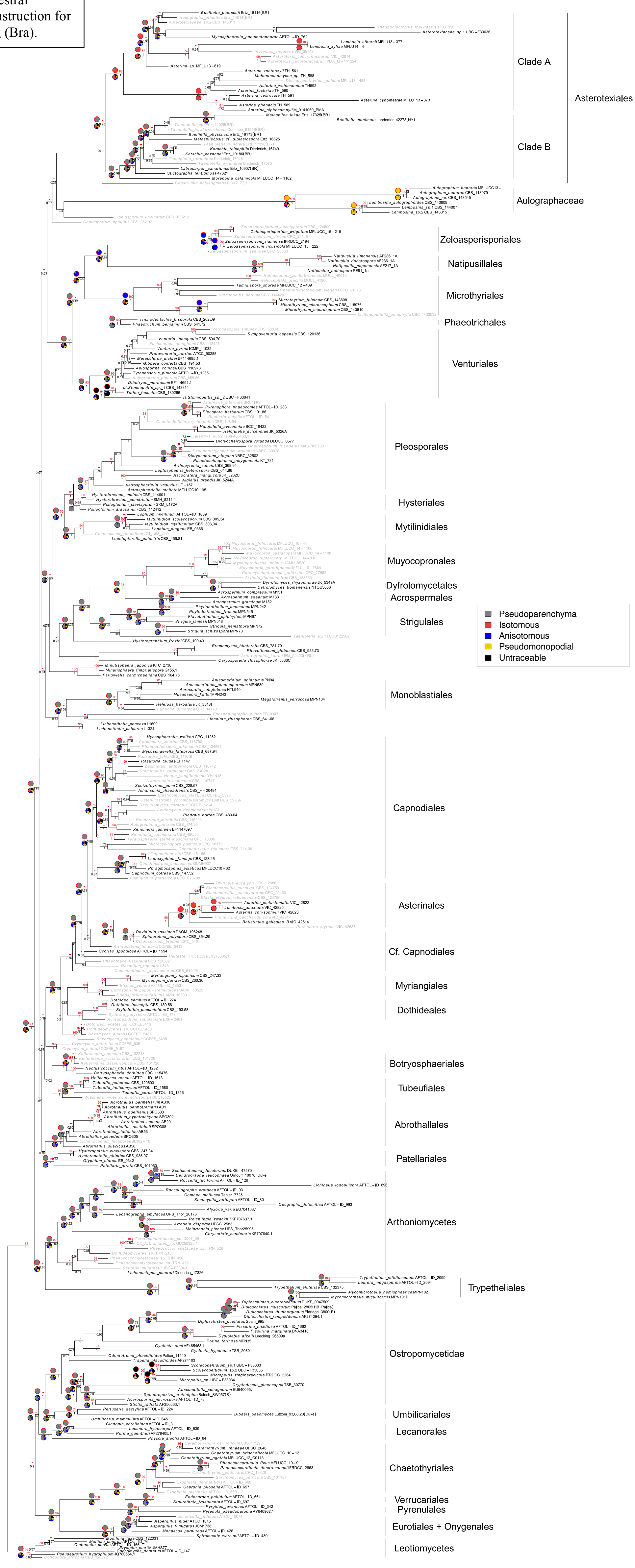

Appendix S16. Ancestral character state reconstruction for margin of sporocarp (Mar).

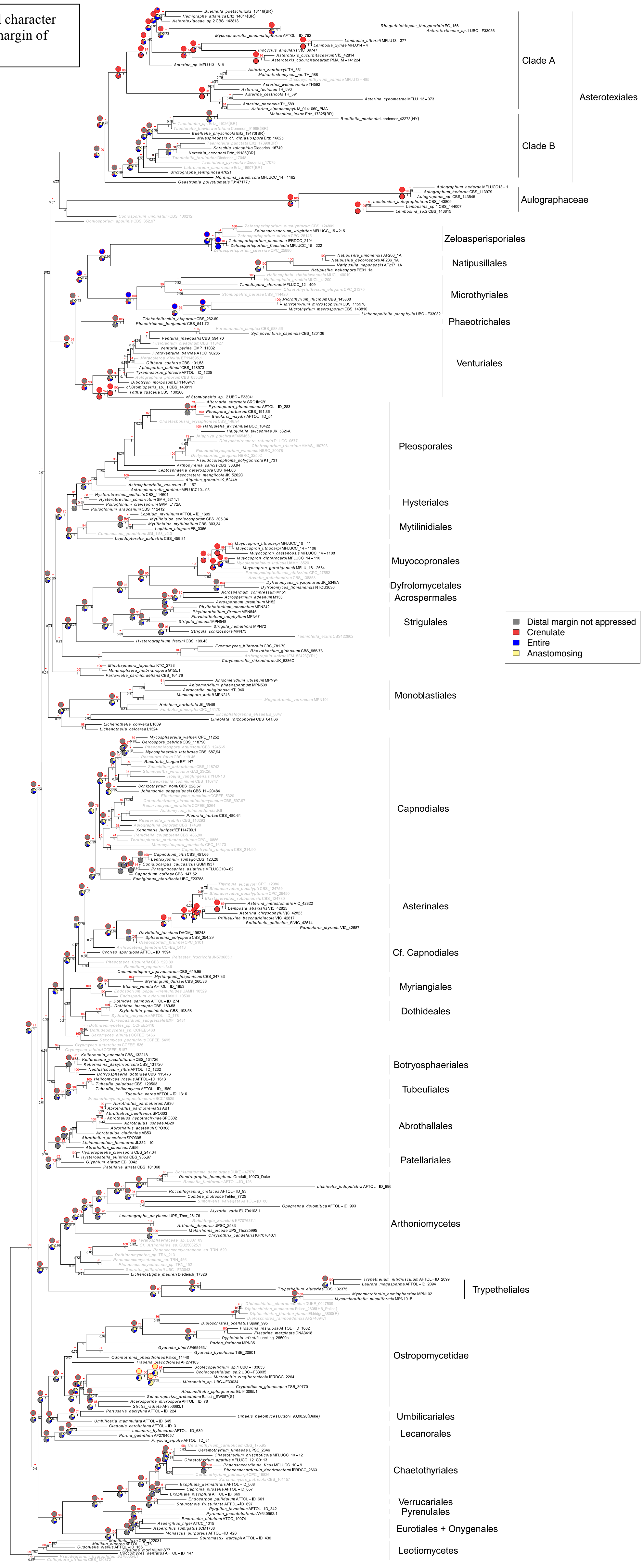

**Appendix S17**. Ancestral character state reconstruction for the presence or absence of lateral appressoria on superficial mycelium (App).

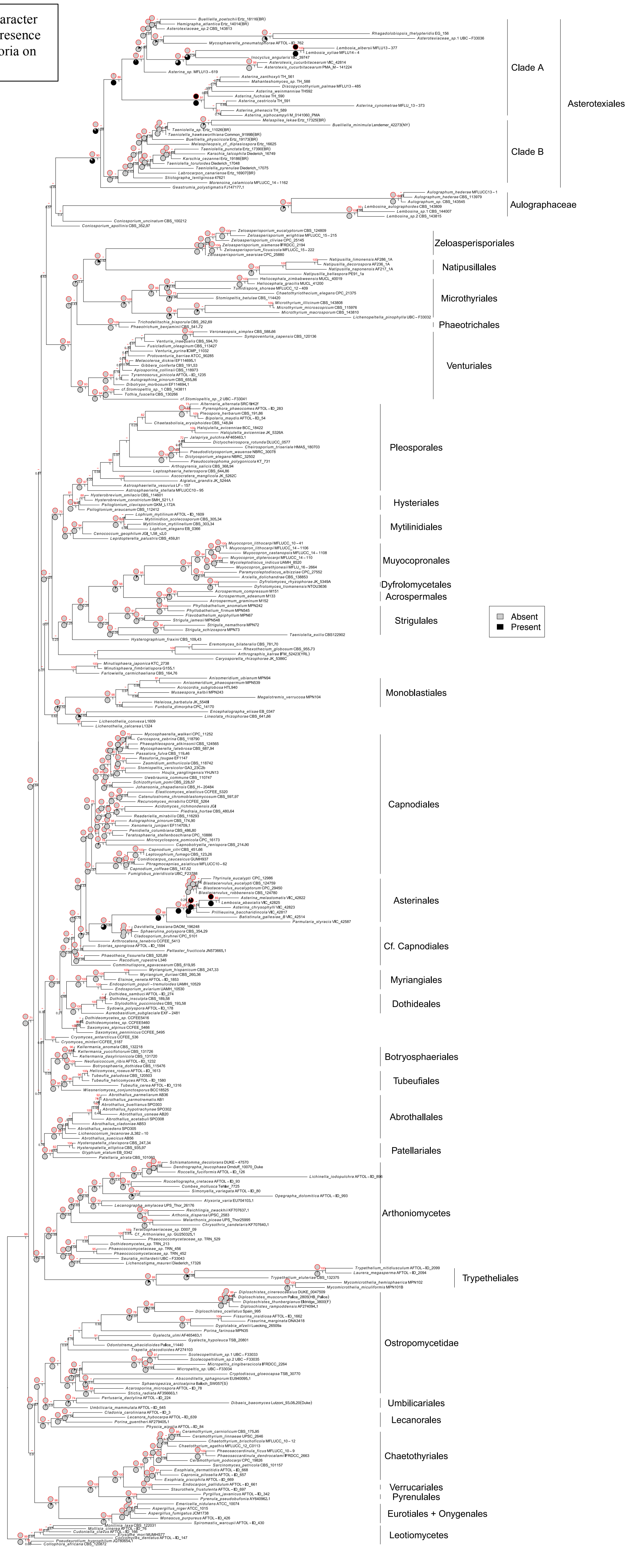

Appendix S18. Ancestral character state reconstruction for Sporocarp initiation (Ini); terminal at tip of generator hypha, intercalary below generator hypha, from spore, from host stomata or from multiple generator hyphae.

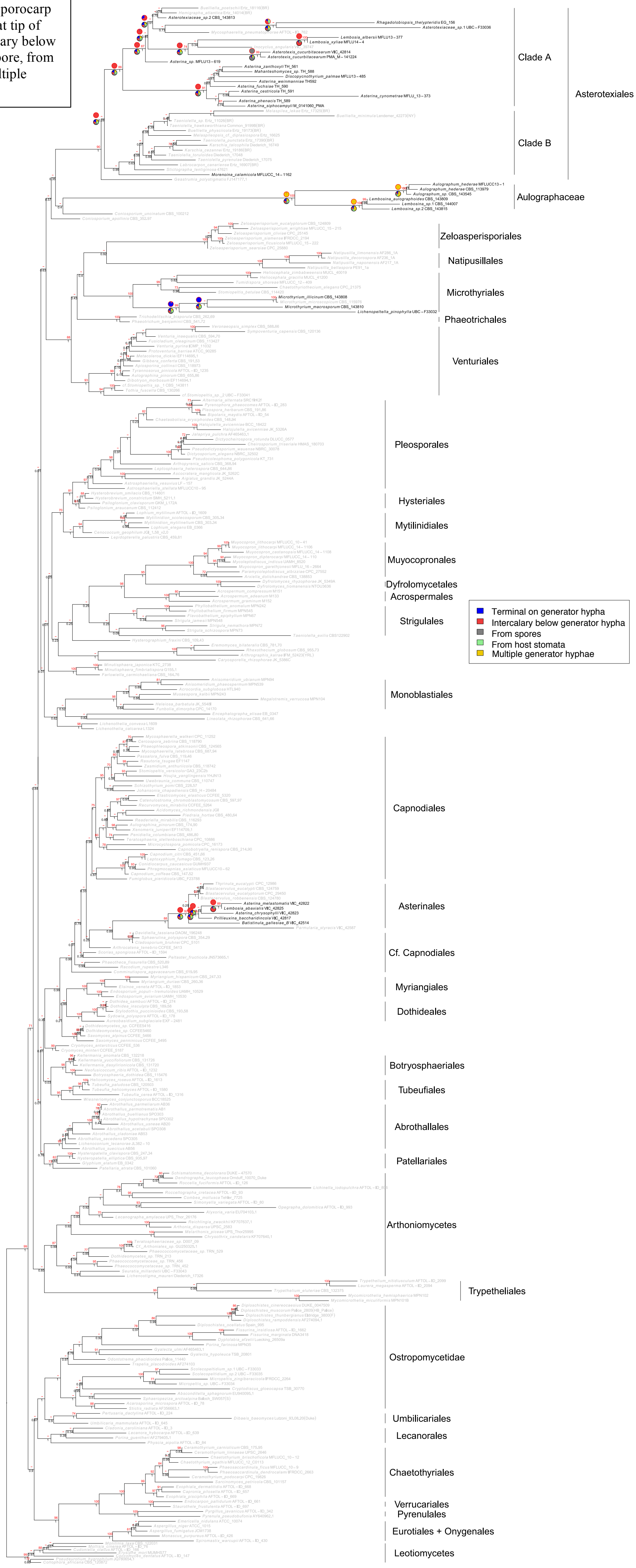

Appendix S19. Consensus of 28 equally most parsimonious phylogenetic positions of the Triassic "fungal thallus" from India described in Mishra et al. (2018).

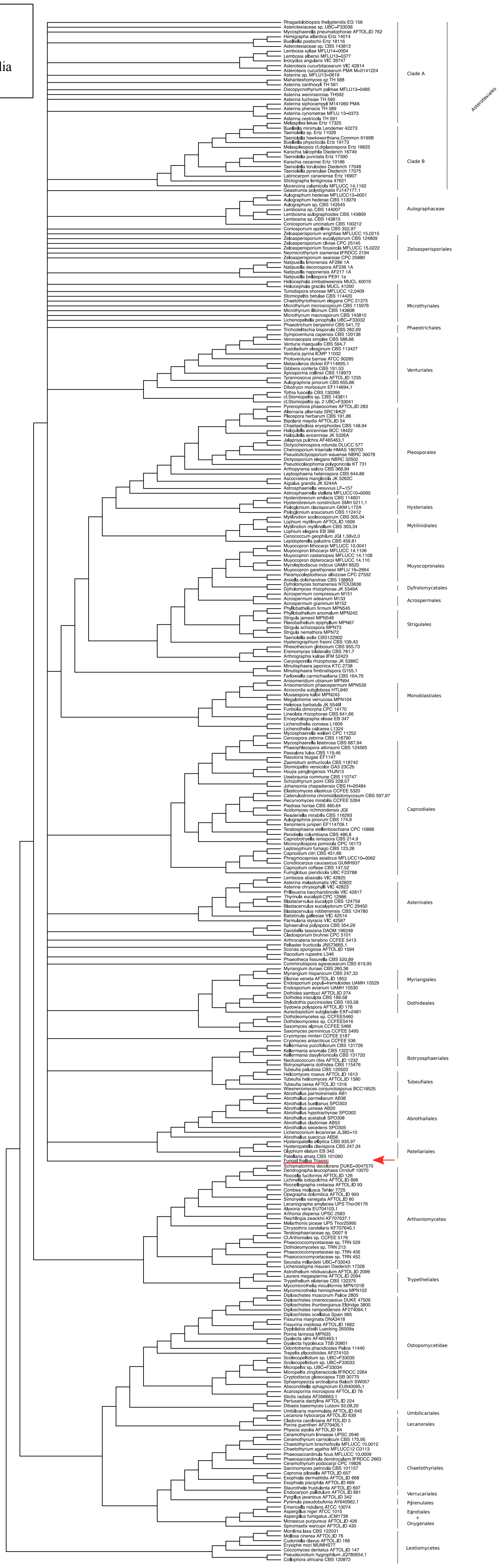

Appendix S20. Consensus of 16 equally most parsimonious phylogenetic positions of *Trichothyrites setifer* from the Eocene of India described in Monga et al. (2015).

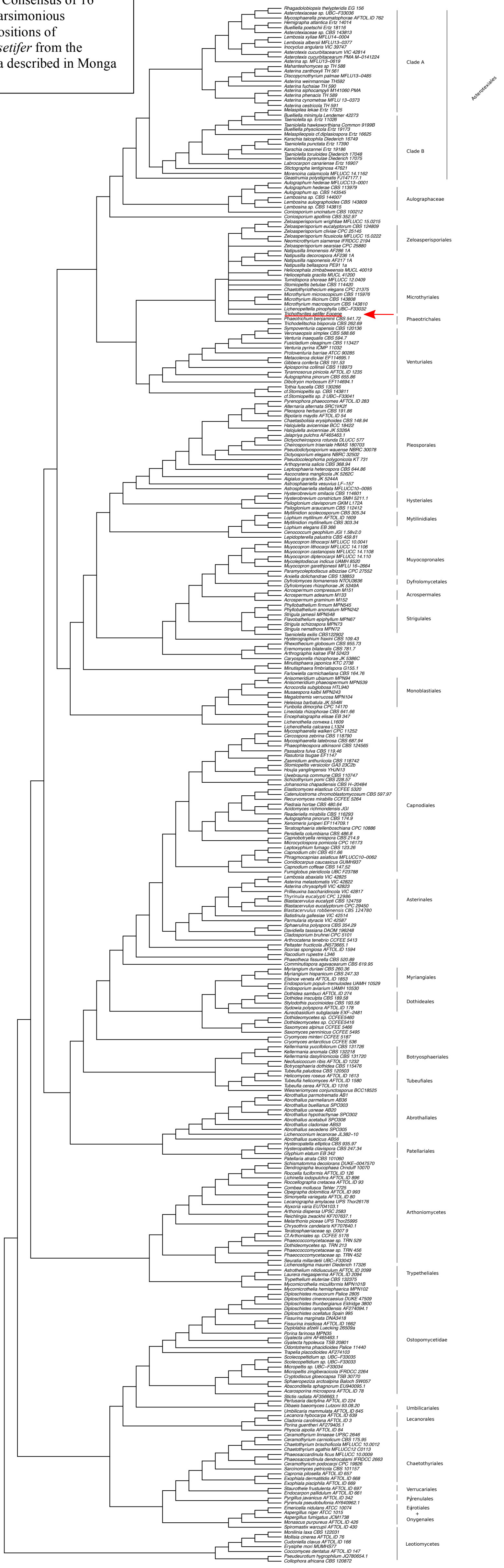

Appendix S21. Consensus of 23 equally most parsimonious phylogenetic positions of *Asterina eocenica* from the Eocene of USA, described in Dilcher (1965).

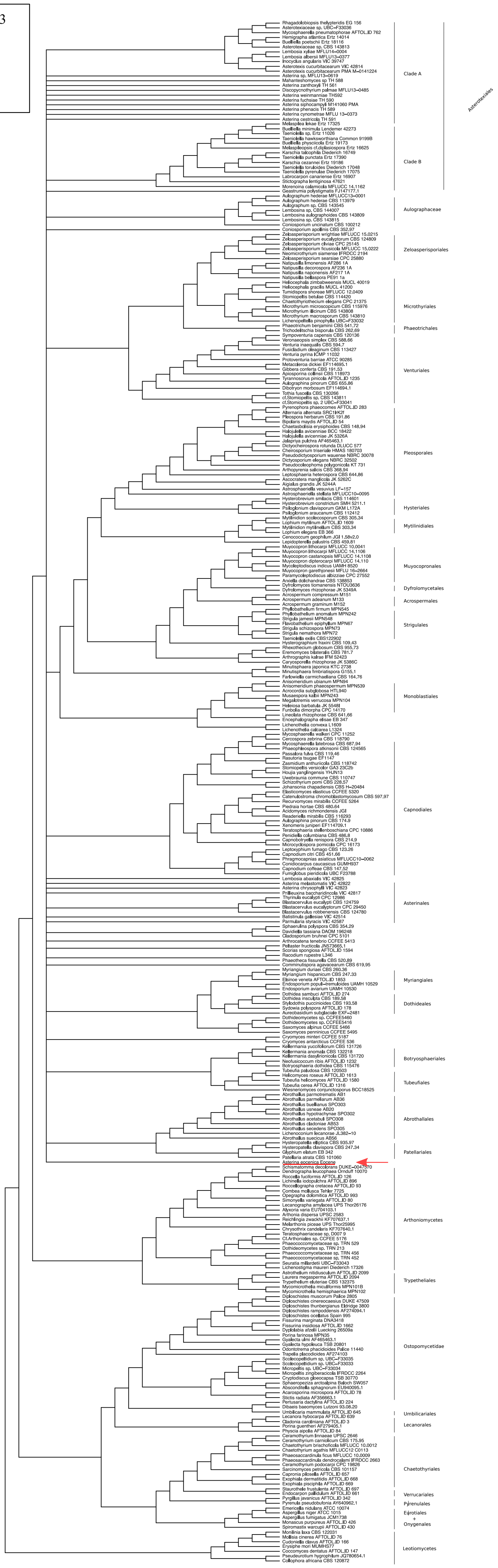

Appendix S22. Consensus of the 18 equally most parsimonious phylogenetic trees placing *Asterina eocenica* with taxa in Asterotexiales.

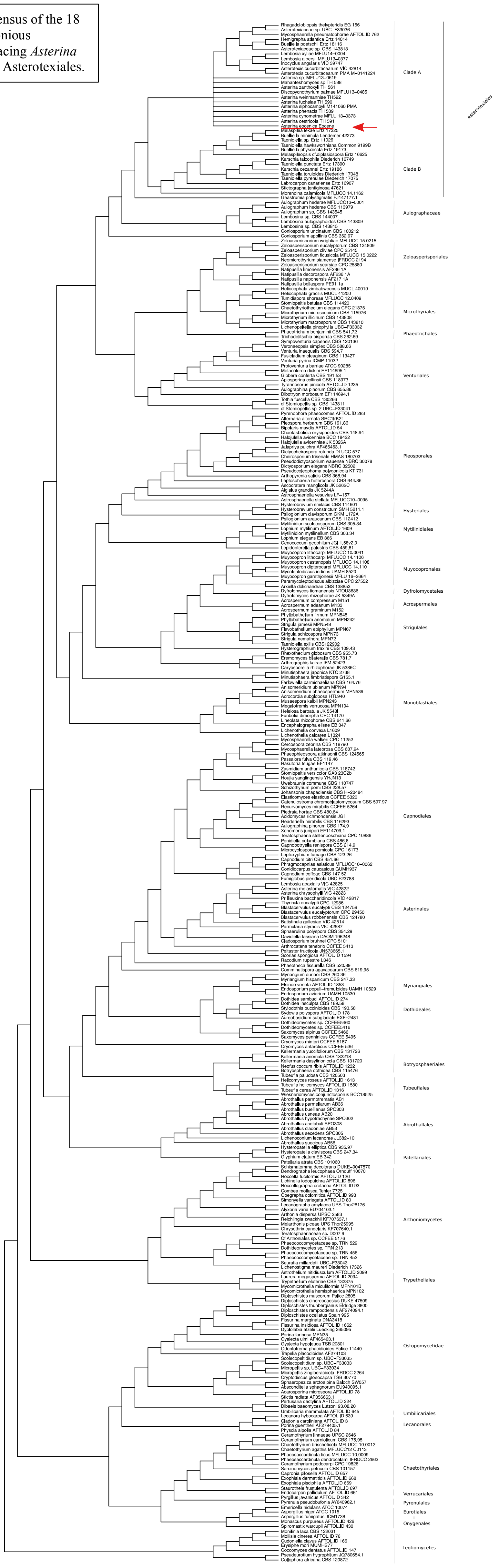

Appendix S23. Consensus of the 5 equally most parsimonious phylogenetic trees placing *Asterina eocenica* with taxa in Asterinales.

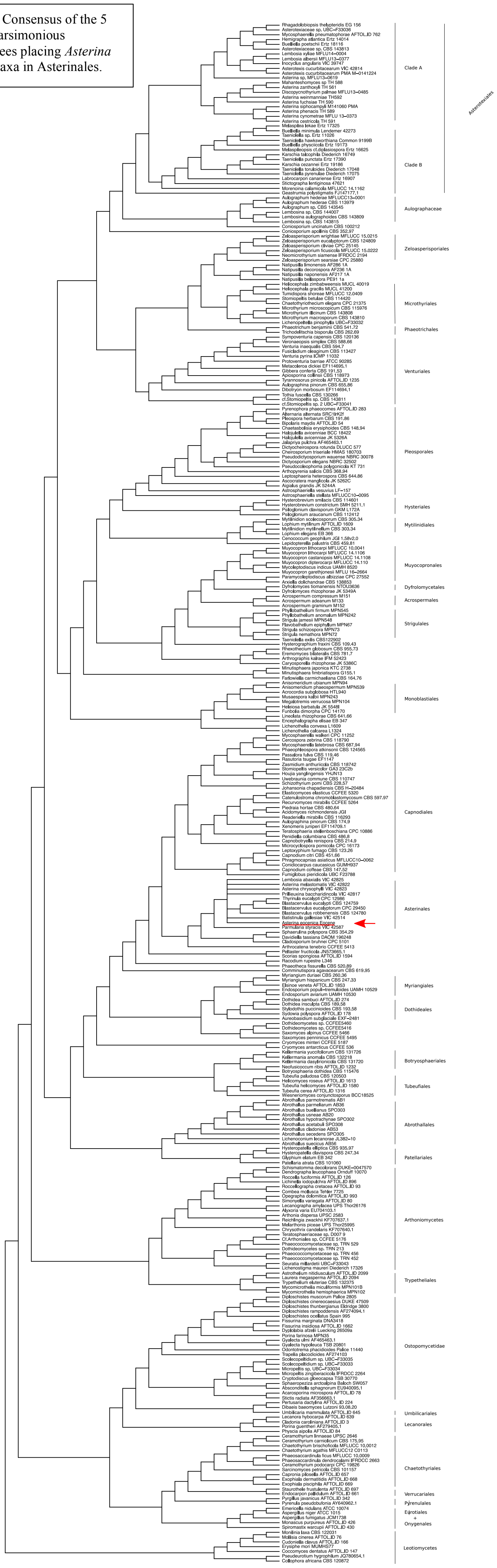
